# Design of SARS-CoV-2 protease inhibitors with improved affinity and reduced sensitivity to mutations

**DOI:** 10.1101/2023.07.19.549739

**Authors:** Michael Westberg, Yichi Su, Xinzhi Zou, Pinghan Huang, Arjun Rustagi, Jaishree Garhyan, Puja Bhavesh Patel, Daniel Fernandez, Yan Wu, Lin Ning, Aimee Beck, Marwah Karim, Chenzhou Hao, Panatda Saenkham-Huntsinger, Vivian Tat, Aleksandra Drelich, Bi-Hung Peng, Shirit Einav, Chien-Te K. Tseng, Catherine Blish, Michael Z. Lin

**Affiliations:** Department of Neurobiology, Stanford University; Stanford, CA, USA; Department of Chemistry, Aarhus University; Aarhus, Denmark; Department of Bioengineering, Stanford University; Stanford, CA, USA; Department of Microbiology and Immunology, The University of Texas Medical Branch; Galveston, TX, USA; Department of Medicine, Stanford University; Stanford, CA, USA; Stanford In Vitro Biosafety Level 3 Service Center, Stanford University; Stanford, CA, USA; Program in Chemistry, Engineering, and Medicine for Human Health (ChEM-H), Stanford University; Stanford, CA, USA; Stanford ChEM-H, Macromolecular Structure Knowledge Center, Stanford University; Stanford, CA, USA; Department of Pathology, The University of Texas Medical Branch; Galveston, TX, USA; Department of Neuroscience, Cell Biology, and Anatomy, The University of Texas Medical Branch; Galveston, TX, USA; Department of Microbiology and Immunology, Stanford University; Stanford, CA, USA; Chan Zuckerberg Biohub; San Francisco, CA, USA; Department of Chemical and Systems Biology, Stanford University; Stanford, CA, USA

## Abstract

Inhibitors of the SARS-CoV-2 main protease (M^pro^) such as nirmatrelvir (NTV) and ensitrelvir (ETV) have proven effective in reducing the severity of COVID-19, but the presence of resistance-conferring mutations in sequenced viral genomes raises concerns about future drug resistance. Second-generation oral drugs that retain function on these mutants are thus urgently needed. We hypothesized that the covalent HCV protease inhibitor boceprevir (BPV) could serve as the basis for orally bioavailable drugs that inhibit SARS-CoV-2 M^pro^ more tightly than existing drugs. Performing structure-guided modifications of BPV, we developed a picomolar-affinity inhibitor, ML2006a4, with antiviral activity, oral pharmacokinetics, and therapeutic efficacy similar or superior to NTV. A crucial feature of ML2006a4 is a novel derivatization of the ketoamide reactive group that improves cell permeability and oral bioavailability. Finally, ML2006a4 is less sensitive to several mutations that cause resistance to NTV or ETV and occur in the natural SARS-CoV-2 population. Thus, anticipatory drug design can preemptively address potential resistance mechanisms.

## Introduction

In the last two decades, three coronaviruses, SARS-CoV-1, MERS-CoV, and SARS-CoV-2, have emerged from animal reservoirs to cause lethal respiratory illnesses in humans. In particular, more than 6.9 million people have already died from COVID-19 (*1*). SARS-CoV-2 has repeatedly evolved to spread more quickly and evade immunity from prior infections or vaccines. While previous infection or three doses of vaccines provide high levels of protection against death, respectively (*2, 3*), the persistently high prevalence and intrinsic virulence of SARS-CoV-2 continue to cause morbidity and mortality, especially in older patients with preexisting medical conditions. Thus, oral anti-coronavirus drugs administered on an outpatient basis to prevent progression to severe disease are desirable.

The coronavirus main protease (M^pro^) is a major target for antiviral drugs (*4*–*6*). M^pro^ is essential for viral replication as it cleaves the large non-structural coronavirus polyprotein into smaller fragments that then assemble to form the RNA replicase (*4*–*6*). Prior to SARS-CoV-2 emergence, experimental inhibitors of M^pro^ were found to protect mice and cats from lethal coronavirus diseases (*7*–*9*). The SARS-CoV-2 M^pro^ inhibitor nirmatrelvir (NTV) (*10*) was proven to reduce hospitalization risk by 89% in unvaccinated COVID-19 patients, leading to its emergency approval in multiple countries (*11, 12*). A non-covalent non-peptidomimetic M^pro^ inhibitor, ensitrelvir (ETV) (*13*), has also shown clinical efficacy and is approved for use in Japan (*14*–*16*). Other oral M^pro^ inhibitors that have entered clinical trials include pomotrelvir (PTV), leritrelvir (*17*), simnotrelvir, EDP-235, and HS-10517 (GDDI-4405). Molecular structures have not been revealed for the last two, and published data are overall sparse for all these inhibitors. Nevertheless, the success of NTV and ETV demonstrate the utility of targeting M^pro^ to treat coronavirus disease.

A major concern, however, is the potential for emergence of drug-resistant SARS-CoV-2 mutants. Genomic surveillance of SARS-CoV-2 has been performed on a large scale since the beginning of the pandemic, enabling the detection of new alleles while they are still rare (*18*–*24*). This may guide the preemptive development of new drugs to address mutants that resist available treatments. Examples of concerning M^pro^ mutations detected via genomic surveillance and laboratory experiments are S144A, E166X, and A173V, which decrease NTV or ETV potency while maintaining viable virus replication (*16, 18, 19, 25*–*34*).

One drug design strategy to reduce the risk of resistance is to mimic the interactions between the substrate and the highly conserved atoms of the enzyme to the greatest extent possible (*35*). Such substrate mimicry reduces the risk of escape mutants since the mutations will both affect inhibitor and substrate binding, with the latter reducing virus fitness. Thus, inhibitors that form strong interactions with essential active site residues of M^pro^ provide a potential route to efficiently maximize ligand binding affinity and minimize the risk of escape mutants.

NTV is a covalent M^pro^ inhibitor, using a nitrile as an electrophilic group (or “warhead”) to react with the nucleophilic C145 in the active site (*10*). Prior to NTV, the only warhead used in approved viral protease inhibitors was the ketoamide group of the hepatitis C virus (HCV) drugs telaprevir (TPV), boceprevir (BPV), and narlaprevir (NPV). These drugs are orally bioavailable and safe, even when taken continuously for > 6 months (*36, 37*). In addition, ketoamides form stable adducts mimicking the tetrahedral transition states and hemiketal/hemithioketal intermediates of substrate proteolysis (*38*–*40*). Indeed, a series of ketoamide-based inhibitors developed for SARS-CoV-1 M^pro^, culminating in 13b, were shown to be active against SARS-CoV-2 M^pro^ (*41, 42*). However, 13b demonstrates a relatively high 50% effective antiviral concentration (EC_50_) of ∼5 µM (*42*), compared to ∼0.1 µM for nirmatrelvir (*10*). Another noteworthy class of reversible covalent inhibitors are aldehydes or their prodrugs, such as GC376 (*6, 43*–*45*). However, aldehydes are considered non-ideal drug candidates due to their metabolic instability, low oral bioavailability, and toxicity caused by off-target reactions (*46*–*48*), including their non-specific inhibition of other cysteine proteases (*49, 50*).

In this study, we address the questions of whether ketoamide inhibitors of M^pro^ can be designed with similar potency and oral bioavailability as NTV, and if they offer an alternative for NTV- or ETV-resistant viral mutants. We hypothesized that BPV supplies a preorganized scaffold and a tight-binding warhead suitable for the generation of robust M^pro^ inhibitors. Our first compound based on this hypothesis, ML1000, efficiently inhibited SARS-CoV-2 M^pro^ with a 50% inhibitory concentration (IC_50_) of 20 nM, but was limited in permeability. We performed structure-guided modifications on side groups and on the ketoamide warhead to improve affinity and permeability. The resulting lead compound ML2006a4 is a picomolar-affinity SARS-CoV-2 M^pro^ inhibitor with oral pharmacokinetics and antiviral efficacy in mice similar or superior to NTV. ML2006a4 binding to M^pro^ is impacted to a lesser degree by several mutations that reduce potency of NTV or ETV, suggesting it may be useful if resistance to these M^pro^ inhibitors arises. Thus, rational antiviral drug design can be used to preemptively address future resistance mutations.

## Results

### HCV protease inhibitors show activity against M^pro^

At the onset of the SARS-CoV-2 pandemic, we asked whether HCV protease inhibitors could be repurposed to inhibit M^pro^. We manually docked BPV into the SARS-CoV-2 M^pro^ active site with good shape complementarity by its P1, P2, and P4 groups (**Fig. 1A**). Strikingly, the bicyclic P2 proline analogue in BPV filled the deep S2 pocket in M^pro^, while its backbone φ and Ψ angles retrace those of the P2 Leu or Phe residues of earlier M^pro^ inhibitors (*42, 51*) (**Fig. 1A** and fig. S1). The distinct bicyclic proline analogue of TPV could also fill S2 while retracing the M^pro^ inhibitor backbone conformation (fig. S1).

**Fig. 1.**
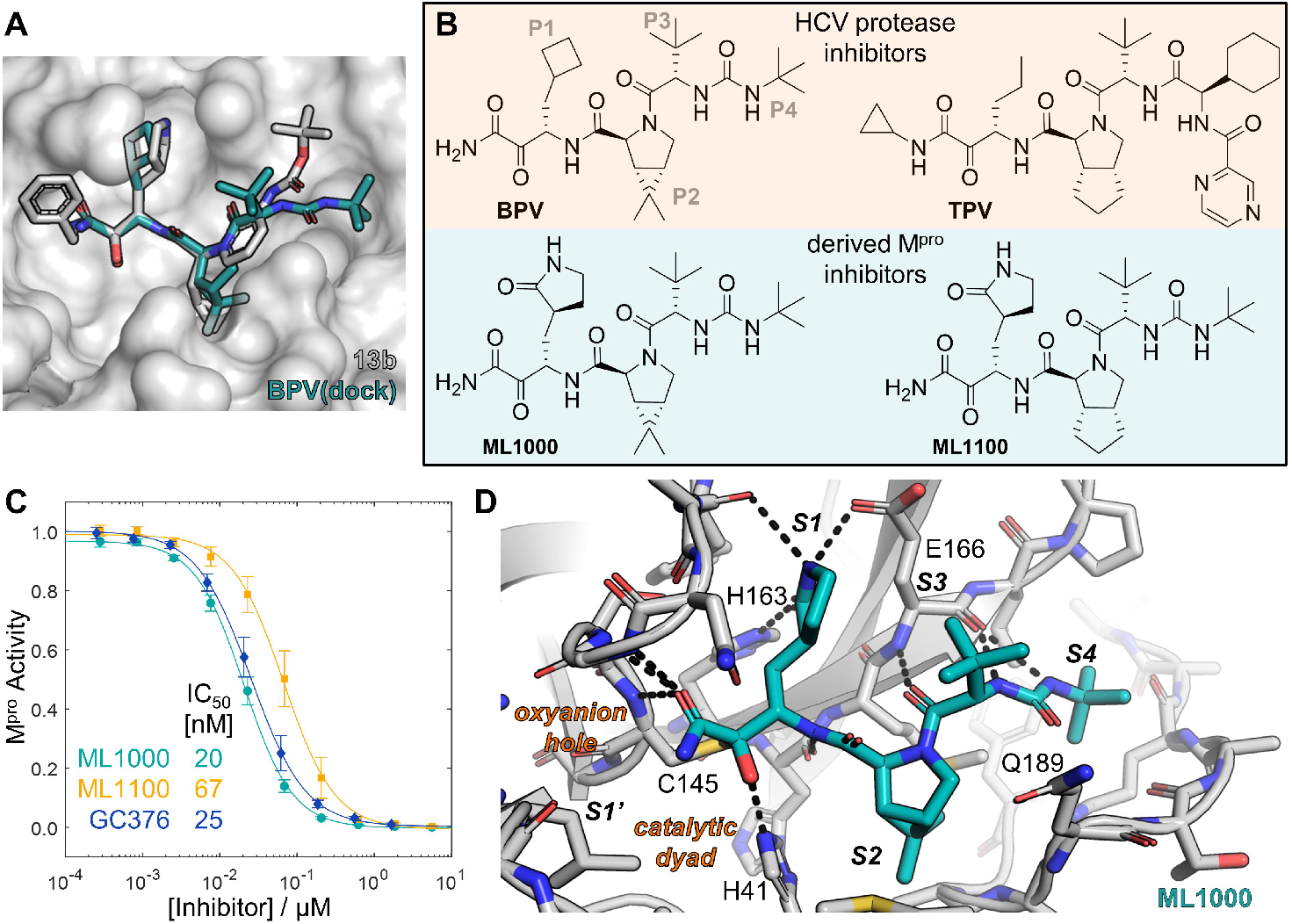
M^pro^ inhibitors derived from HCV protease inhibitors. (**A**) Manual rigid docking of BPV into a 13b-M^pro^ co-crystal structure (PDB 6Y2G, protomer B). Using PyMOL, BPV was manually placed into the SARS-CoV-2 M^pro^ active site and unconstrained bonds were rotated for optimal complementarity. (**B**) Structures of BPV and TPV and the derived ML1000 and ML1100. ML1000 is essentially BPV with a P1 γ-lactam. ML1100 replaces the bicyclic P2 proline analog of BPV with that of TPV. (**C**) Inhibition of M^pro^-coil activity by GC376, ML1000, and ML1100 at 25 °C after 1 h of preincubation. Error bars are SD (*N = 3*). (**D**) Crystal structure of the ML1000-M^pro^ adduct confirms the expected binding mode with H-bond interactions highlighted. Nucleophilic attack of C145 at the ketoamide warhead of ML1000 creates a hemithioketal and allows for additional non-covalent interactions with H41 and the oxyanion hole defined by the backbone NHs of residues 143-145.

Other aspects of BPV are also compatible with M^pro^ binding. The P3 group of M^pro^ substrates face out into solution (*52*), as does the t-butyl group in the analogous position of BPV, TPV, and NPV. The P3 backbone NH in all three inhibitors can form a hydrogen bond (H-bond) with the backbone carbonyl of M^pro^ E166. However, the P4 groups of TPV and NPV appear too large for the S4 pocket (fig. S1). We thus hypothesized that BPV and, to a lesser extent, NPV and TPV, may be able to inhibit SARS-CoV-2 M^pro^.

After finding that BPV and TPV inhibited SARS-CoV-2 M^pro^ in mammalian cells and enzymatic assays (fig. S2), we measured IC_50_ values for BPV, TPV, NPV, and other proposed M^pro^ inhibitors (fig. S3). BPV, TPV, and NPV all showed activity against M^pro^, with BPV being the most potent (IC_50_ = 6.2 µM). In the course of our study, at least three other groups reported BPV inhibiting SARS-CoV-2 M^pro^ at low micromolar concentrations (*53*–*55*). Two of these studies included crystal structures of the BPV-M^pro^ adduct that validate our manually docked model (*54, 55*).

### A boceprevir-based M^pro^ inhibitor

We next sought to design inhibitors that combine the bicyclic P2 rings of BPV or TPV with side groups optimized for M^pro^ binding. A P1 Gln residue is strongly preferred by all CoV M^pro^ species (*56*). This preference is conserved in the related enterovirus 3C proteases such as human rhinovirus (HRV) protease and even with the more distantly related plant potyvirus proteases (*57, 58*). Beginning with the HRV protease inhibitor rupintrivir, enterovirus and CoV protease inhibitors have incorporated a γ- or δ-lactam group at P1 as a Gln mimic (*59*–*61*). We initially designed two compounds with a γ-lactam at P1 and a bicyclic ring at P2 (*62*): ML1000, with the P2, P3, P4, and ketoamide groups of BPV, and ML1100 with the P2 group of TPV and the P3, P4, and ketoamide groups of BPV (**Fig. 1B**). We retained the P3 t-butyl of BPV as it was preferred at this position in substrate screens (*63*), and as its lipophilicity could aid permeability.

ML1000 and ML1100 were potent M^pro^ inhibitors in solution, with measurable IC_50_ values limited by the ∼100 nM M^pro^ concentrations required to detect activity on a FRET substrate. To maintain M^pro^ activity at low nanomolar concentrations we engineered a M^pro^-coil fusion protein for avidity-enhanced dimerization and thereby measurable activity at 1 nM. In this assay, ML1000 and ML1100 produced IC_50_ values of 20 nM and 67 nM, respectively (**Fig. 1C** and fig. S4). A ML1000-M^pro^ crystal structure confirmed the P1 γ-lactam was positioned similarly to other M^pro^ inhibitors, with the other atoms matching the manually docked BPV structure (**Fig. 1D**).

However, ML1000 and ML1100 incorporate 6 H-bond donors (HBDs) and 6 H-bond acceptors (HBAs), an unfavorable number for membrane permeability (*64*). Indeed, their apparent permeability (P_app_) in Caco-2 cell layers was low (table S1), and they failed to inhibit replication of SARS-CoV-2 (USA-WA1/2020 strain expressing NLuc) in cell culture even at 100 µM (fig. S5 and S6). Nevertheless, we hypothesized the ketoamide-based inhibitors would be useful for maximizing drug-target interactions, as the warhead allows P1’ extensions and, after reaction, mimics the tetrahedral transition state and hemithioketal substrate intermediate with multiple H-bonds to M^pro^ (**Fig. 1D**). Thus, we chose ML1000 as a lead compound for developing high-affinity oral M^pro^ inhibitors that are robust to protease mutation.

### Optimizing permeability and potency

We performed stepwise optimization of ML1000 to improve permeability and potency (**Fig. 2, A and B**). To aid in drug screening, we also devised a bioluminescent assay for efficiently measuring intracellular inhibition of M^pro^ (i.e., IC_50,cell_). In this assay, M^pro^ matures autocatalytically and cleaves a luciferase; M^pro^ inhibition thus leads to luminescence increases (fig. S7). A larger IC_50_/IC_50,cell_ ratio indicates better permeability.

**Fig. 2.**
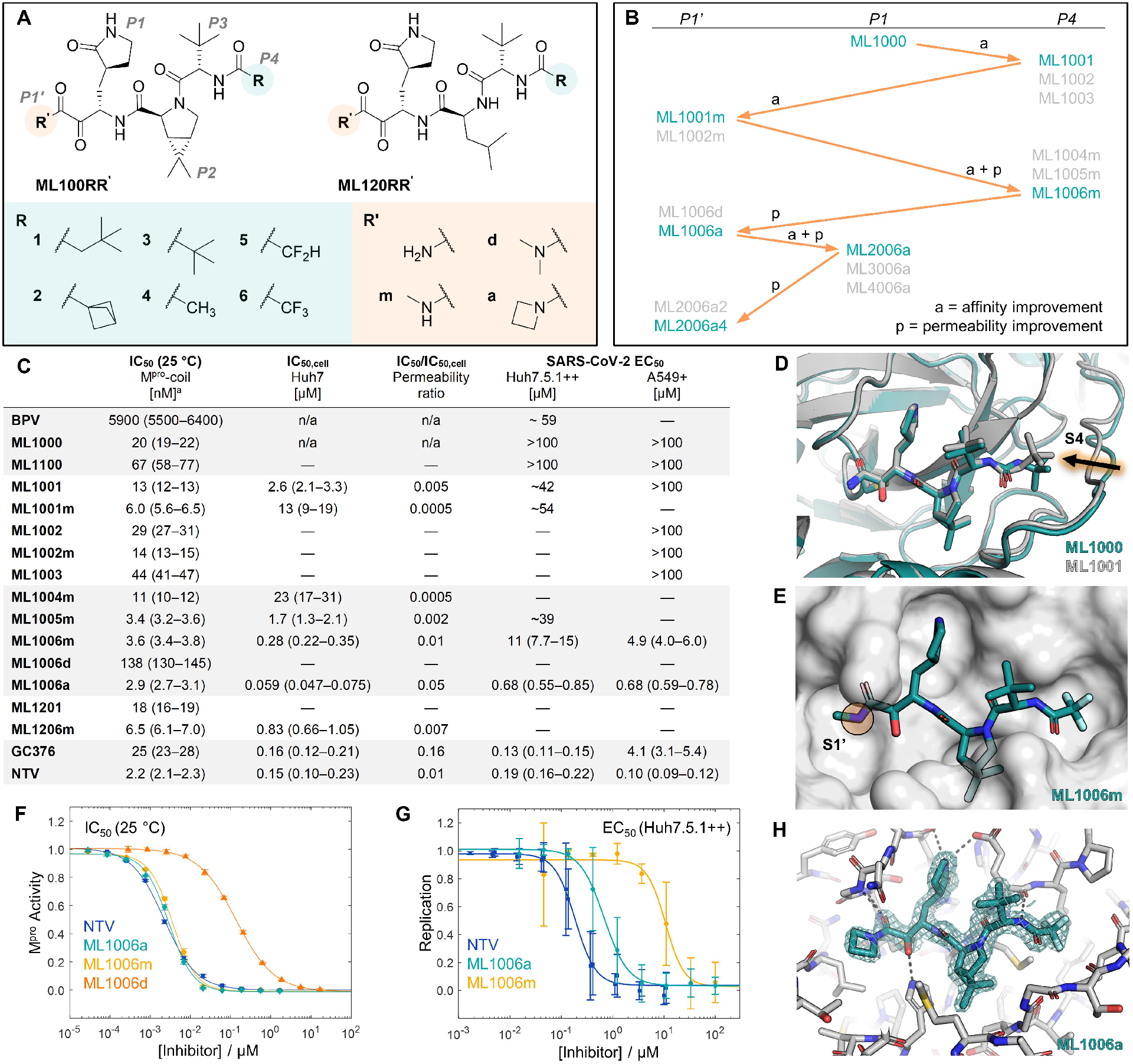
Optimizing permeability and potency of ML1000. (**A**) M^pro^ inhibitors based on ML1000. Additionally, select non-constrained inhibitors (P2 = Leu) were synthesized to assess the effects of P2 rigidification. (**B**) Progressing from ML1000, stepwise optimization of P4 (R), P1’ (R’), and P1 was performed. (**C**) Inhibitor activity against purified M^pro^-coil (IC_50_), intracellular M^pro^ (IC_50,cell_), and SARS-CoV-2 replication (EC_50_). CI_95_ are reported in parentheses and data, fits, and statistical details are found in fig. S4 to S7. ^a^IC_50_ data was measured at 25 °C after 1 h of preincubation. In contrast to NTV, the ketoamides have not reached equilibrium. (**D**) Co-crystals of M^pro^ with ML1000 or ML1001 show that ML1000 expands the S4 pocket while the flexible ML1001 P4 cap swings out of the contracted S4 pocket. (**E**) A M^pro^-ML1006m co-crystal highlights limited space for *N,N*-substitution and extension of the ketoamide into S1’. (**F**) M^pro^-coil inhibition curves show significantly reduced affinity for ML1006d but not ML1006a. (**G**) Inhibition of SARS-CoV-2 replication in Huh7.5.1++ improves significantly from ML1006m to ML1006a. (**H**) The M^pro^-ML1006a adduct structure confirms the presence of the expected planar azetidinylated ketoamide. Error bars are SD.

First, in ML1001 through ML1003, we exchanged the rigid and bulky urea of ML1000 (fig. S8) for an amide with various alkyl groups, thereby removing a HBD and optimizing the fit of the P4 cap (**Fig. 2A**). ML1001 exhibited the lowest IC_50_, but IC_50,cell_ and antiviral EC_50_ remained high (**Fig. 2C**).

Second, to better mimic P1’ substrate interactions and to improve permeability, we replaced a ketoamide hydrogen in ML1001 and ML1002 with N-methyl, creating ML1001m and ML1002m. While this lowered IC_50_, suggesting the methyl interacts favorably with the S1’ pocket, it did not improve IC_50,cell_ or antiviral EC_50_ (**Fig. 2C**).

Third, we shrunk P4, as the ML1001-M^pro^ crystal structure revealed the 2,2-dimethylpropyl was not fully buried inside the S4 pocket (**Fig. 2D**). We synthesized ML1004m through ML1006m with methyl, difluoromethyl, and trifluoromethyl P4 caps (**Fig. 2, A and B**). As a benchmark, we also synthesized NTV, which at this time in our study was revealed to be identical with ML1001 in P1, P2, and P3 (*10*). Difluoromethyl and trifluoromethyl groups (ML1005m and ML1006m) produced IC_50_ values of 3.4 and 3.6 nM, respectively, lower than the 11 nM of the methyl (ML1004m) (**Fig. 2C**). We selected ML1006m as the new lead based on IC_50,cell_, but its antiviral EC_50_ remained > 4 µM (**Fig. 2C**).

Fourth, we removed all HBDs from the ketoamide to boost lipophilicity and thereby permeability. As a ML1006m-M^pro^ crystal structure revealed limited space around the ketoamide (**Fig. 2E**), we added minimally sized ‘a’ and ‘d’ substitutions (**Fig. 2A**). ML1006d contains a *N,N*-dimethylketoamide, but IC_50_ increased to 138 nM (**Fig. 2F**), possibly due to methyl-carbonyl intramolecular clashes and associated twisting within the ketoamide (*65, 66*). ML1006a, in which we added a strained four-membered azetidine ring as a compact non-clashing *N,N*-substitution, maintained an IC_50_ of 2.9 nM (**Fig. 2F**). ML1006a achieved antiviral EC_50_ of 0.68 µM, a clear improvement over ML1006m (**Fig. 2G**), but still inferior to NTV. A ML1006a-M^pro^ structure revealed the expected planar azetidinyl ketoamide in the S1’ pocket (**Fig. 2H**).

Fifth, we remodified P1 to improve lipophilicity. ML2006a enlarged P1 by one carbon atom to form a δ-lactam as seen in other viral protease inhibitors (*60, 67*–*69*), while ML3006a and ML4006a incorporated “inverted lactams” to remove a HBD (**Fig. 3A**). ML2006a improved on IC_50,_ IC_50,cell_ and antiviral EC_50_ (**Fig. 3, B and C**, and fig. S9). ML3006a and ML4006a were more permeable based on the IC_50_/IC_50,cell_ ratio and P_app_ measurements (**Fig. 3C** and table S1), but IC_50_, IC_50,cell_, and EC_50_ were all increased (**Fig. 3, B and C**), as expected from the removal of a strong H-bond with E166 (fig. S10).

**Fig. 3.**
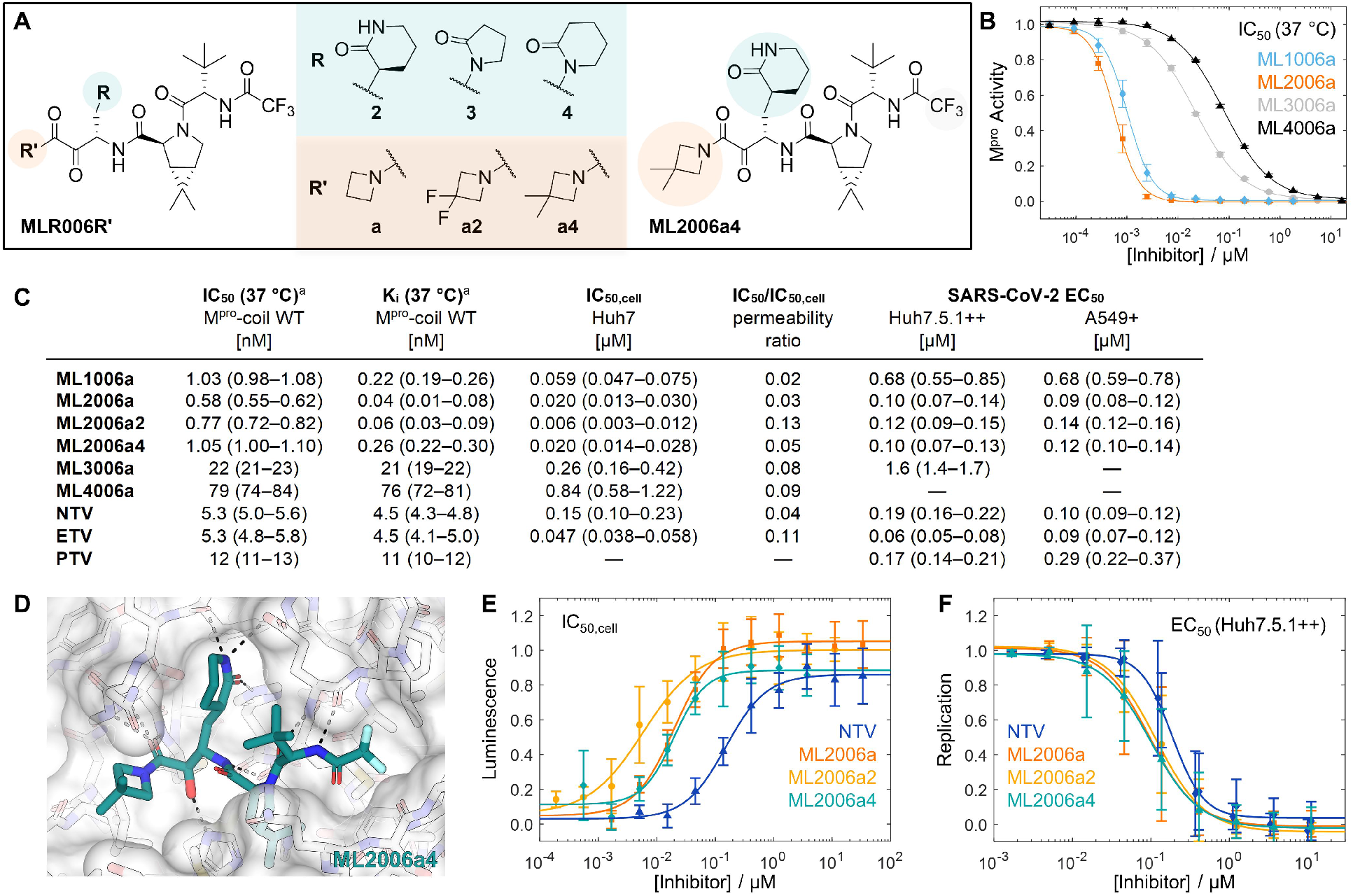
Discovery of SARS-CoV-2 M^pro^ inhibitors with higher potency than NTV in vitro. (**A**) P1 lactamyls and azetidine substitutions were tested to improve permeability. (**B**) IC_50_ curves showing improved M^pro^ inhibition by ML2006a, but loss of activity for ML3006a and ML4006a. (**C**) Tabulation of inhibitor activity against purified M^pro^-coil (IC_50_ and K_i_), intracellular M^pro^ (IC_50,cell_), and SARS-CoV-2 replication (EC_50_). CI_95_ are reported in parentheses and data, fits, and statistical details are found in fig. S5, S6, S7, S9, and S13. ^a^Measured at 37 °C after 3 h of preincubation. (**D**) Crystal structure of the covalent adduct of ML2006a4 and SARS-CoV-2 M^pro^. (**E**) ML2006a derivatives have improved permeability and effectively inhibit intracellular M^pro^ activity and luciferase proteolysis. (**F**) SARS-CoV-2 replication in Huh7.5.1++ cell cultures is effectively inhibited by ML2006a derivatives. Error bars are SD.

In a final step to improve lipophilicity, we tested difluoro and dimethyl 3,3-substitutions of the azetidine of ML2006a, creating ML2006a2 and ML2006a4 (**Fig. 3A**). The larger 3,3-dimethyl azetidinyl of ML2006a4 is just contained within S1’ (**Fig. 3D** and fig. S10). These compounds retained the low IC_50_ and IC_50,cell_ of ML2006a (**Fig. 3, C to E**) while improving permeability (**Fig. 3C** and table S1). Gratifyingly, ML2006a, ML2006a2, and ML2006a4 achieved antiviral EC_50_ values of 100–120 nM in Huh7.5.1-ACE2-TMPRSS2 (Huh7.5.1++) cells, lower than the 190 nM of NTV (**Fig. 3C and F**). In A549-ACE2 (A549+) and Calu-3 cells, their EC_50_ values were similar to NTV (**Fig. 3C** and fig. S11).

### Orally dosable ketoamide

We performed in vitro ADME assays at several steps during inhibitor optimization (supplementary text). Like NTV (*10*), the ketoamide-based inhibitors also undergo degradation by CYP3A, an effect counteracted by ritonavir (RTV) (table S1). We measured pharmacokinetics (PK) in mice for ML1006m, ML1006a, ML2006a, ML2006a4, and NTV (**Fig. 4, A and B)**, all formulated in a saline solution with PEG-300, DMSO, and Tween-80 (PDT formulation). As expected, pre-dosing of RTV lowers clearance of all five compounds. Oral bioavailability (F) improves throughout the ketoamide series, culminating with 27% or 115% for ML2006a4 without or with RTV, respectively. While ML2006a2 has antiviral potency and in vitro ADME properties comparable to ML2006a4, including human microsome and plasma stability, its low stability in mouse plasma precluded *in vivo* characterization (table S1). When dosed orally at 20 mg/kg with RTV in PDT, mouse plasma concentrations of NTV and ML2006a4 exceeded plasma protein binding-adjusted antiviral EC_90_ and EC_50_ (EC_90,p_ and EC_50,p_) levels for ∼8 h and ∼12 h after administration, respectively (**Fig. 4B**).

**Fig. 4.**
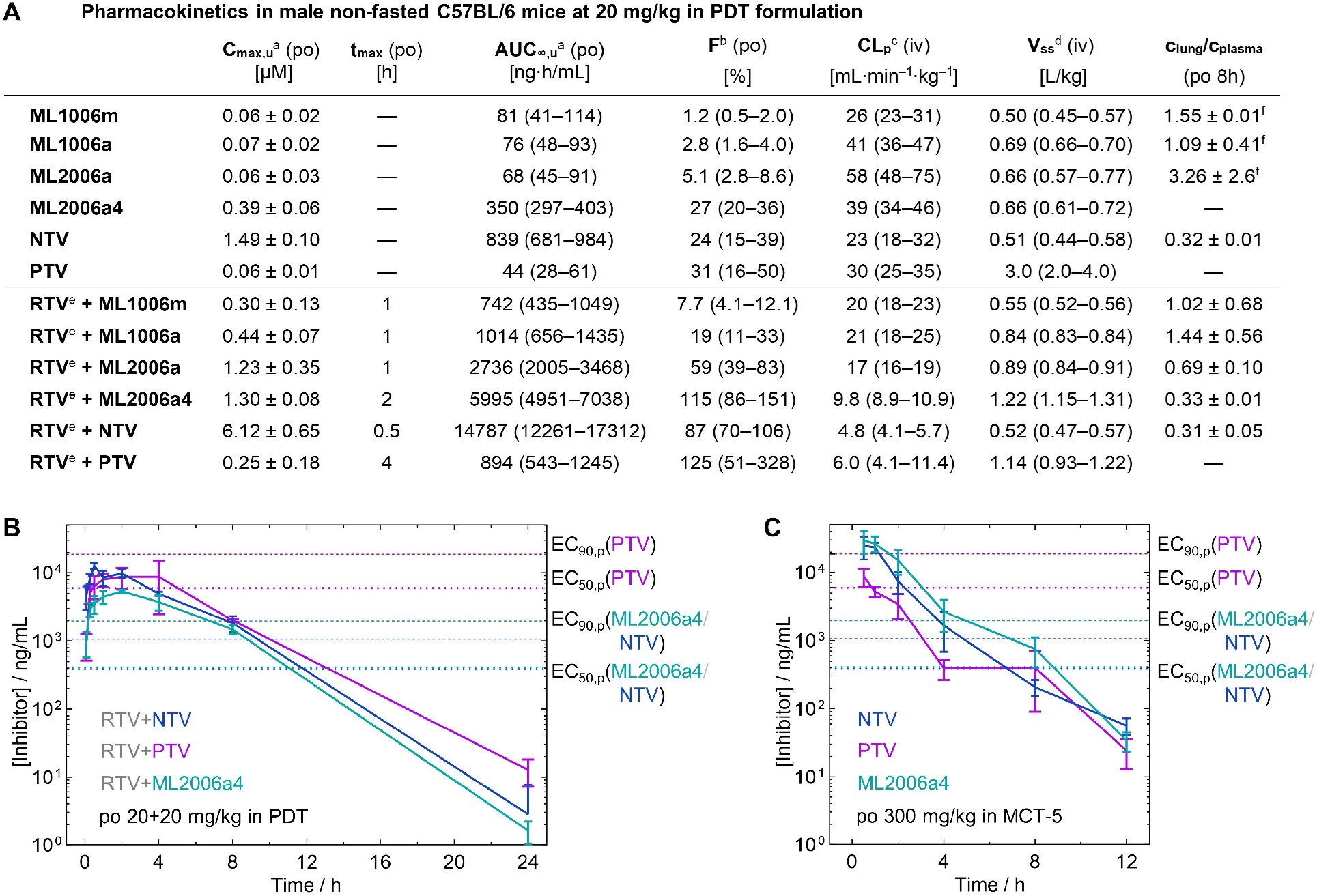
ML2006a4 is orally bioavailable in mice. (**A**) Key parameters from non-compartmental analysis of pharmacokinetic studies performed in mice with oral (po) dosing at 20 mg/kg and intravenous (iv) dosing at 2 mg/kg in a saline solution with 40% PEG300, 10% DMSO, and 5% Tween-80 (PDT). In parentheses, parameter intervals calculated using non-compartmental analysis of the plasma concentration mean ± SD (*N = 3*). For direct experimental observations, ± SD are reported. ^a^Calculated using f_u_ reported in table S1. ^b^Oral bioavailability. ^c^Plasma clearance rate. ^d^Volume of distribution. ^e^RTV dosed po at 20 mg/kg 30 min prior to test compound. ^f^(*N = 2*) (**B**) Inhibitor plasma concentration monitored for 24 h after administration of a 20 mg/kg RTV pre-dose followed by single-doses of 20 mg/kg ML2006a4, NTV, or PTV solutions in PDT. The overall profile is similar for ML2006a4 and NTV that each maintain concentrations above EC_90,p_ measured in Huh7.5.1++ cells for ∼8 h. In contrast, PTV never reaches EC_90,p_. (**C**) Inhibitor plasma concentrations monitored for 12 h after single-agent administration of 300 mg/kg ML2006a4, NTV, or PTV suspensions in 0.5% methylcellulose and 5% Tween-80 (MCT-5).

As NTV demonstrated initial evidence of efficacy in mice at a high dose without RTV (10), we also asked how ML2006a4 would compare to NTV in PK with a high-dose single-agent regimen. When administered orally at 300 mg/kg in aqueous suspensions with methylcellulose and Tween-80, conditions similar to those used previously for NTV (10), mouse plasma concentrations of ML2006a4 and NTV were comparable at most time points, although NTV fell below EC_50,p_ before 8 h, while ML2006a4 reached EC_50,p_ at 8 h (**Fig. 4C)**.

We also obtained ADME and PK data for PTV, a nitrile-based M^pro^ inhibitor that was recently tested in humans as a RTV-free COVID-19 treatment but failed to achieve the clinical endpoints. The clearance of PTV by human liver microsomes was higher than that of NTV or ML2006a4, suggesting that PTV is not better suited for a single-agent regimen than NTV or ML2006a4 (table S1). In addition, metabolism of PTV was only partly suppressed by RTV (table S1), indicating that enzymes beyond CYP3A contribute to its degradation. In mice, total plasma concentrations of PTV closely matched NTV and ML2006a4 when dosed orally with RTV at 20+20 mg/kg in the PDT formulation (**Fig. 4B**). However, due to a high level of plasma protein binding (table S1), EC_50,p_ is substantially higher for PTV than for NTV and ML2006a4, leading to rapid decay to plasma concentrations below EC_50,p_ (**Fig. 4B**). In addition, when administered at 300 mg/kg without RTV, PTV plasma levels were lower than those of NTV and ML2006a4, and dropped below EC_50,p_ after 1 h. Thus, our data indicates that PTV is an inferior SARS-CoV-2 antiviral compared to NTV and ML2006a4 either with or without RTV.

### Mouse safety and efficacy

Having identified ML2006a4 as the lead compound with the most favorable PK, we assessed its toxicity in vitro and in vivo. In cytotoxicity tests, ML2006a4 demonstrated CC_50_ > 100 µM (table S2). We then compared the tolerability of ML2006a4 and NTV in mice. Protease inhibitor (40 mg/kg) and RTV (20 mg/kg) were orally administered twice daily (b.i.d.) as co-suspension in 0.5% methylcellulose and 2% Tween-80 (MCT-2). This dosing regimen was chosen to simulate the human equivalent dose of NTV, and maintained plasma concentrations at or above EC_50,p_ for ∼6 h (**Fig. 5A** and fig. S12). All groups gained weight similarly over the 4 days (fig. S12). While no clinical, biochemical, or hematological abnormalities were observed for the ML2006a4+RTV combination, NTV+RTV induced elevated liver enzymes in 3 of 6 mice, similar to published reports (supplementary text). Thus, 40+20 mg/kg of ML2006a4+RTV in MCT-2 appeared at least as safe as NTV+RTV at similar doses and was used for the subsequent efficacy study.

**Fig. 5.**
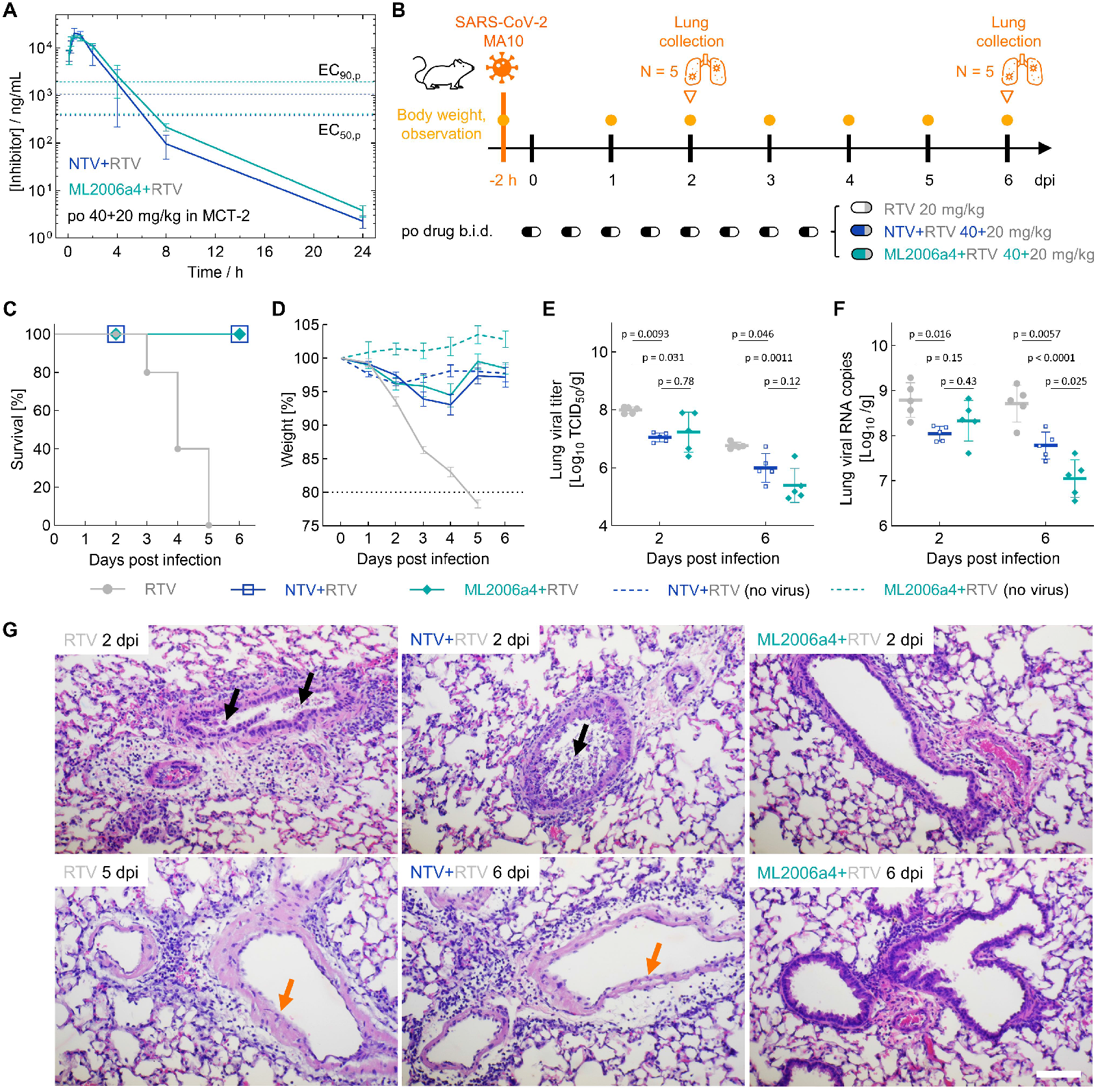
ML2006a4 is efficacious in mice. (**A**) Pharmacokinetics of the dosing regimen used for safety and efficacy testing in mice. Plasma concentrations were monitored for 24 h after administration of co-suspensions of 40+20 mg/kg ML2006a4+RTV or NTV+RTV in 0.5% methylcellulose and 2% Tween-80 (MCT-2). (**B**) Protocol for efficacy testing of ML2006a4 in 16-week-old female BALB/c mice. After intranasal infection with SARS-CoV-2 MA10 (10^5^ TCID_50_), the mice received oral b.i.d. dosing for 4 days. ML2006a4+RTV and NTV+RTV were dosed as in panel A, and the RTV-only control was dosed at 20 mg/kg. All infected groups contained 10 mice with 5 mice sacrificed at 2 dpi and 6 dpi, respectively. Uninfected control groups contained 5 mice that were sacrificed at 6 dpi. (**C**) Daily survival curves and (**D**) body weights for all five groups. No death or clinical decline was observed for any of the ML2006a4+RTV or NTV+RTV treated groups. (**E**) Infectious viral titers and (**F**) viral RNA levels were determined in lungs harvested at 2 and 6 dpi, respectively. For the RTV-only group, data plotted at 6 dpi are from lungs collected at death (actual time 3–5 dpi). (**G**) Representative images of H&E-stained lungs collected at 2 or 6 dpi. Scale bar is 80 µm. Histopathological analysis identified peribronchial and perivascular infiltrations with mononuclear cells in the three infected groups. However, infiltration was less pronounced in the ML2006a4+RTV group at both 2 and 6 dpi. RTV and NTV+RTV groups exhibited severe injury in respiratory epithelia (black arrows) at 2 dpi and did not show complete epithelial regeneration (orange arrows) at 6 dpi. Epithelial layers in the ML2006a4+RTV group showed minimal rupture at 2 dpi and more complete regeneration at 6 dpi.

To assess the in vivo therapeutic efficacy of ML2006a4, 16-week-old BALB/c mice were infected with mouse-adapted SARS-CoV-2 MA10, and drugs were administered orally b.i.d. for 4 days (**Fig. 5B**). Within 5 days post infection (dpi), SARS-CoV-2 MA10 was lethal to all mice in the RTV-only control group, while administration of NTV+RTV or ML2006a4+RTV fully rescued the infected mice and ensured minimal weight loss and no clinical abnormalities (**Fig. 5, C and D**). Viral titers and viral RNA levels in lung tissues were determined at 2 dpi and at the end of the study (i.e., terminal or 6 dpi). At both timepoints, ML2006a4+RTV or NTV+RTV showed significant reductions in infectious viral titers compared to RTV alone (**Fig. 5E**). At 6 dpi, viral titers and RNA levels trended lower in ML2006a4-treated mice than in NTV-treated mice, with the RNA difference being statistically significant (**Fig. 5, E and F**). Histopathological evaluation of lungs also revealed that, compared to NTV, ML2006a4 conferred better lung protection with less inflammation and respiratory epithelial injury at 2 dpi, as well as better epithelial regeneration at 6 dpi (**Fig. 5G**). In summary, ML2006a4 was as efficacious as NTV in promoting survival of mice infected with SARS-CoV-2 and provided superior viral suppression and lung protection.

### Comparison of ketoamide and nitrile adduct stability

To accurately assess binding affinity of the tighter inhibitors and to allow comparisons to recent data, we used the Morrison equation to calculate the equilibrium inhibitory constant, K_i_, as previously described for NTV (*10*). We find that the δ-lactam containing ketoamide compounds inhibit M^pro^ at ∼20 to 100-fold lower concentrations than NTV or ETV at 37 ºC. In particular, we measured K_i_ values of 0.26 nM for ML2006a4 vs. 4.5 nM for NTV and ETV (**Fig. 3C** and fig. S13). Our K_i_ value for NTV closely matched the published value, confirming the accuracy of our measurements (*10*). To our knowledge, ML2006a4 is the first orally dosable SARS-CoV-2 M^pro^ inhibitor reported with picomolar affinity.

To understand the basis for the higher affinity of ML2006a4 relative to NTV, we measured inhibition kinetics (fig. S14) and calculated dissociation kinetics (table S3) of the reversible hemithioketal and thioimidate adducts (*70*). Ketoamide-based inhibitors exhibit slower unbinding kinetics than NTV, with the ML2006a4 adduct showing a half-life of 5 h compared to <2 min for NTV. Our results align with other findings of slow dissociation by ketoamides, e.g., BPV and TPV from HCV protease (*71*) and leritrelvir from SARS-CoV-2 M^pro^ (*17*).

### Preorganization in affinity and specificity

To understand the role of preorganization at P2 in inhibitor affinity and specificity, we synthesized ML1201 and ML1206m as analogs of ML1001 and ML1006m, respectively, with a flexible P2 leucine (**Fig. 2A**). Both showed higher IC_50_ than the parental proline-containing compounds (**Fig. 2C**). While a P2 Leu allows a H-bond between its backbone NH and the side-chain of M^pro^ Gln189 (see e.g. PDB 6LU7, 6XHL, and 7K0H) (*51, 72, 73*), this additional H-bond apparently cannot compensate for the entropic penalty of binding the more flexible ML1201 and ML1206m. Thus, the preorganized proline backbone, also present in ML2006a4 and NTV, allows removal of a HBD to enhance permeability while also improving binding energy.

We assessed the specificity of ML2006a4 and other M^pro^ inhibitors vs. human cathepsins, which are common targets for protease inhibitors with electrophilic warheads (*50*). While BPV has an acceptable side effect profile despite sub-micromolar inhibition of Cat-K and Cat-S (*74*), broader inhibition of cathepsins by aldehyde-based drugs present a potential safety concern (*50, 69, 75*). We tested for inhibition of four human cathepsins: Cat-B/K/L/S (fig. S15). Cat-S appears to be the most promiscuous, as all ketoamides, except ML1006d, inhibit Cat-S with IC_50_ < 1.5 µM. However, at physiologically relevant concentrations, ML2006a4 shows minimal inhibition of Cat-B/K/L. Interestingly, ML1201 and ML1206m, which contain Leu at P2, show broader inhibition across the cathepsins than ML1001 and ML1006m with the Pro-derived P2 ring, revealing that preorganization at P2 also serves as a specificity determinant.

### Anticipating resistance

The history of HIV, HCV, and influenza antivirals cautions that resistance is likely to arise (*76*). While NTV mimics the substrate envelope in P1–P4 (*77*), it differs from natural substrates in lacking a P1’ analogue or the ability to form a tetrahedral adduct. Several potential NTV escape mutants have already been identified (*26*–*28*). One concerning M^pro^ mutation is S144A, as it maintains substantial proteolytic activity (*19, 25, 78*), is positioned near the reactive C145, was expanded during propagation of CoVs in the presence of NTV (*26*–*28*), and has been observed in natural SARS-CoV-2 genomes (*18, 19, 27*). Indeed, we observed a 12-fold K_i_ increase for NTV, from 4.5 nM against wild-type (WT) M^pro^ to 55 nM against M^pro^ S144A (**Fig. 6, A and B**). Similarly, S144A increased K_i_ 9.3-fold for nitrile-based PTV (**Fig. 6B**), 5.9-fold for aldehyde-based GC376, and 4.6-fold for hydroxymethyl ketone-based PF00835231 (*79*) (table S4). Notably, the non-covalent inhibitor ETV saw a 31-fold increase in K_i_, which is even more severe than for NTV (**Fig. 6B**). In contrast, ML2006a4 remained potent against M^pro^ S144A (**Fig. 6A**) with only a 2.4-fold increase in K_i_ (**Fig. 6B**).

**Fig. 6.**
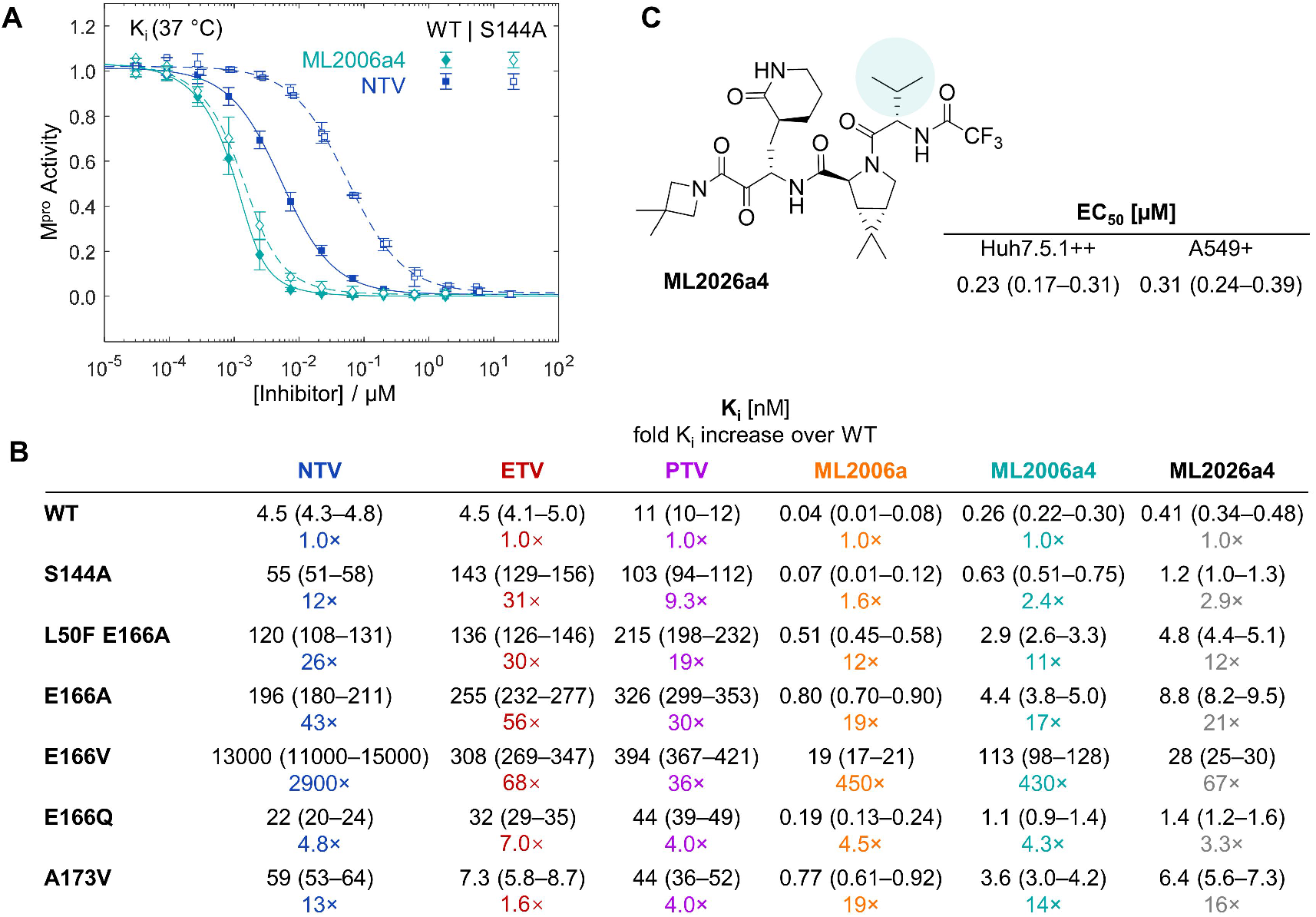
ML2006a4 has improved mutation tolerance. (**A**) Comparison of in vitro inhibition curves against M^pro^-coil WT and S144A shows a severe loss of potency for NTV but not ML2006a4 against M^pro^ S144A. Error bars are SD. (**B**) Inhibition constants, K_i_, at 37 °C for M^pro^-coil WT and six mutants show that ML2006a and ML2006a4 tolerate S144A and E166A better than NTV, ETV, and PTV. E166V most dramatically affects NTV binding. **(C)** ML2026a4 has a smaller P3 group relative to ML2006a4 and is less sensitive to the E166V (panel B). The antiviral EC_50_ of ML2026a4 is increased slightly compared to ML2006a4 (Fig. 4C). CI_95_ are reported in parentheses and data, fits, and statistical details are found in figs. S5, S6, S13, and S17.

To understand the structural basis for the differential susceptibility of NTV and ketoamide-based inhibitors to S144A (table S4), we obtained crystals of M^pro^ S144A bound to NTV or ML1006a. However, these did not reveal any clear differences from WT M^pro^ adducts (fig. S16). In the absence of conformational changes in the static crystals, we hypothesize that the ML1006a hemithioketal adduct forms a stronger interaction with the perturbed oxyanion loop of M^pro^ S144A compared to the thioimidate adduct of NTV. In addition, the P1’ extension may provide further binding redundancy to help maintain recognition of mutant M^pro^. However, a detailed mechanism for S144A-induced NTV resistance may require dynamical computer simulations of covalent bond formation and breaking.

Other reported NTV escape mutants include M^pro^ E166X (*26*–*28*) and A173V (*26, 27, 30*). Mutation of E166 is surprising since this residue is strictly conserved across all CoVs (*80*) and involved in M^pro^ dimerization (*81, 82*). However, even if fitness loss is seen in single point mutants, several mutations outside the active site (e.g. T21I, L50F, P252L, and T304I) have been observed to restore viral fitness of a range of active site mutants including S144A and E166X (*27, 28*). Based on these findings, we also tested the effect of E166Q/A/V and A173V mutations on the M^pro^ affinities of NTV and ML2006a4 (**Fig. 6B**). Among these, L50F E166A, E166A, and E166V conferred the largest reductions in NTV affinity, and in all cases impacted ML2006a4 binding less. Specifically, the K_i_ of NTV vs. ML2006a4 increased 26-vs. 11-fold for L50F E166A, 43-vs. 17-fold for E166A, and 2900-vs. 430-fold for E166V (**Fig. 6B** and fig. S17). The other mutants tested, E166Q and A173V, conferred smaller and equal K_i_ increases for NTV and ML2006a4. In summary, ML2006a4 exhibits reduced sensitivity to the M^pro^ mutants that cause the largest loss of NTV binding.

We also tested the effects of the above mutations on binding of ETV and PTV, which differ from NTV and ML2006a4 in lacking BPV-derived moieties. First, ETV does not interact with the S4 pocket (fig. S18), making ETV insensitive to A173V that affects binding of NTV, ML2006a4, and, to a lesser extent, PTV (**Fig. 6B**). Compared to ML2006a4 the K_i_ values of ETV and PTV were less affected by E166V but more sensitive to E166A (**Fig. 6B**). In fact, E166V and E166A equally increased the K_i_ of ETV (68- and 56-fold) or PTV (36- and 30-fold). Both E166V and E166A may destabilize the dimer interface and the flexible S1 pocket (*83*), thereby affecting interactions with the M^pro^ inhibitors.

Next, we investigated potential mechanisms for the distinct sensitivity of NTV and ML2006a4 to E166V. Structural modelling indicates that E166V, but not E166A, can cause a steric clash with the P3 t-butyl group of ML2006a4 and NTV (fig. S18), which was derived from BPV and absent in ETV and PTV. To test the role of the t-butyl, we synthesized ML2026a4 with a smaller isopropyl group in its place (**Fig. 6C**). As expected, ML2026a4 was less sensitive to E166V than ML2006a4, with a K_i_ increase of 67-fold vs. 430-fold. Interestingly, the 67-fold reduced affinity of ML2026a4 against M^pro^ E166V was similar in magnitude to the 68-fold loss of affinity for ETV, which lacks a P3 group altogether.

Against WT M^pro^ and other mutants, the K_i_ of ML2026a4 was within 2-fold of ML2006a4, indicating that the P3 alteration in ML2026a4 does not substantially impact recognition of M^pro^. In line with the K_i_ values, the antiviral EC_50_ values were only modestly higher for ML2026a4 than ML2006a4 (<3-fold) at 0.23 and 0.31 µM in Huh7.5.1++ and A549+, respectively (**Fig. 6C**). Thus, P3 branch reduction in ML2026a4 corrects the particular sensitivity of ML2006a4 to the E166V mutation, and may be a strategy for reducing the E166V-sensitivity of NTV and other BPV-derived M^pro^ inhibitors as well.

## Discussion

In this study, we develop ML2006a4 as a picomolar-affinity orally dosable SARS-CoV-2 M^pro^ inhibitor with similar or superior antiviral efficacy to NTV at identical concentrations in vitro and identical doses in vivo. Overall, ML2006a4 and ML2026a4 are less sensitive to M^pro^ mutations compared to NTV and have mutational sensitivity profiles distinct from ETV. Our results identify ML2006a4 and ML2026a4 as potential preclinical drug candidates that can be developed to prepare for future surges of NTV- or ETV-resistant coronaviruses.

Interestingly, antiviral efficacy of the combination of NTV and RTV, the two ingredients in the Paxlovid oral COVID-19 treatment, had until recently not been reported in animals with initial reports focusing on high-dose (300 mg/kg) NTV alone (*10, 84, 85*). Recent studies tested the NTV+RTV combination for protection against severe disease at 150+10 mg/kg in K18-hACE2 mice (*86, 87*) and at 250+83 mg/kg in hamsters (*88*). Our results confirm that moderate doses of NTV+RTV (40+20 mg/kg), equivalent to the clinical doses in humans, do also efficiently rescue BALB/c mice from SARS-CoV-2-induced death. With this model in hand, we were able to compare ML2006a4 and NTV at identical doses, in identical vehicles, and with identical amounts of RTV, in simultaneous parallel experiments. The comparison revealed that ML2006a4 has improved antiviral efficacy and confers better lung protection compared to NTV in mice. In addition, we found that therapeutic plasma levels of NTV or ML2006a4 could also be achieved without ritonavir by administering higher oral doses (**Fig. 4C** and **5A**).

In the course of developing ML2006a4, we demonstrated that a P2 proline ring is essential for its potency. Multiple clinically approved protease inhibitors incorporate cyclic structures that improve affinity by preorganizing the molecules for binding (*89, 90*). In energetic terms, this is considered pre-paying the entropic penalty of binding-induced rigidification (*91*–*93*). Additionally, preorganization can improve selectivity and stability by limiting alternative conformations and disfavoring non-specific interactions (*93*–*95*). Here, we directly compare inhibitors with flexible leucine or fixed proline P2 backbones, allowing us to identify the P2 proline as a major contributor to affinity, permeability, and specificity in ML2006a4 and NTV.

Another major outcome of this work is the development of a novel derivative of the ketoamide warhead to improve lipophilicity and oral bioavailability. Specifically, the azetidine substitution introduced here solves the permeability limitations of ketoamide-based SARS-CoV-2 M^pro^ inhibitors. While azetidines are common motifs in drugs (*96*), the electrophilic azetidinylated α-ketoamide is an underexplored motif. To our knowledge, only three limited literature reports on the plain azetidinyl exist (*38*–*40*). Our demonstration of enhanced permeability and pharmacokinetics by azetidinyl derivatives should greatly expand the medicinal utility of ketoamides in tight-binding protease inhibitors and other covalent drugs.

Our study also reveals notable differences between ketoamide and nitrile warheads. First, ketoamides exhibit much slower bidirectional inhibition kinetics with residence times of 5 h for ML2006a4 and 16 h for ML2006a2. It would be interesting to determine if adduct stability could effectively extend the pharmacodynamic profile and contribute to improved dosing protocols (*71, 97*). Second, the azetidinyl substitution provides a rigid platform for combined transition state or intermediate mimicry and interaction with S1’. In contrast, the nitrile warhead does not allow for extension or tuning.

Finally, ML2006a4 and ML2026a4 retain higher potency against several M^pro^ mutants containing resistance-associated mutations of NTV or ETV. We speculate that the more extensive interactions of the ketoamide adduct or the azetidinyl group may provide a degree of redundancy in binding absent in NTV. Future work will focus on cross-resistance in the context of the full viral replication cycle by testing the antiviral potency of ML2006a4 against SARS-CoV-2 replicons and recombinant viruses with NTV- or ETV-resistant mutations (*27, 28*). The possibility of NTV resistance arising is especially concerning given the routine occurrence of rebound cases with high viral levels in NTV-treated patients, as these >3-week infections provide favorable conditions for mutations to arise and amplify (*26, 98*).

## Conclusion

We report the rational development of ML2006a4 as the first oral picomolar-affinity SARS-CoV-2 M^pro^ inhibitor. Our design centers on a preorganized P2 bicyclic proline derivative, which we show improves affinity, permeability, and specificity. Azetidinyl derivatives of the ketoamide remove H-bond donors and improves membrane permeability and oral bioavailability, which is otherwise a key challenge for this potent but polar class of warheads. Similar to NTV, ritonavir-boosted ML2006a4 has high oral bioavailability in mice, and rescues mice from an otherwise lethal SARS-CoV-2 infection. Importantly, the activity of ML2006a4 against M^pro^ is less sensitive to several mutations conferring NTV or ETV resistance, serving as an example of anticipatory drug design to counteract resistance. Finally, the new ketoamide derivative reported here may find broad use in small-molecule inhibitors of other disease-relevant proteases.

## Acknowledgements

The authors thank Edward Anhoa Pham, Jeffrey S. Glenn, Mark Smith, Paul Humphries, Chaitan Khosla, and Nathanael S. Gray (Stanford University) for helpful discussions. The authors also thank Brett Hurst and Bart Tarbet, Utah State University, for initial service on antiviral replication assays. MW thanks Nikolaj L. Villadsen and Finn Ø. Sørensen for helpful discussions. The authors thank Shanghai Chempartner for excellent contract compound synthesis and characterization. A portion of this work was performed at the Stanford ChEM-H Macromolecular Structure Knowledge Center and the Stanford in vitro Biosafety Level 3 Service Center.

The pET His6 TEV LIC cloning vector (2B-T) was a gift from Scott Gradia (Addgene plasmid # 29666 ; http://n2t.net/addgene:29666 ; RRID:Addgene_29666). SARS-CoV-2-NLuc in the form of a passage 1 stock was a gift from Jacob Hou and Ralph Baric (UNC Chapel Hill). The A549-ACE2 cells were a gift from Ralf Bartenschlager (Heidelberg University). The Huh7.5.1-ACE2-TMPRSS2 cells were a Gift from Andreas Puschnik (UCSF).

The project is supported by a COVID-19 Response grant from Stanford ChEM-H and the Innovative Medicines Accelerator (MZL), a Harrington Scholar-Innovator award (MZL), Fast Grants 2082 and 2246 for COVID-19 from Emergent Ventures at the Mercatus Center at George Mason University (MZL and CB), a Stanford-Coulter Translational Research Grant (MZL and CB), an Antiviral Drug Discovery (AViDD) center funded by NIH/NIAID award U19AI171421 (MZL, CB, JG, and CTKT), a grant from the Denver Foundation (CB), grant NNF18OC0031816 from the Novo Nordisk Foundation and the Stanford Bio-X Program (MW), a Bio-X Stanford Interdisciplinary Graduate Student Fellowship (XZ), a E&M Foundation, Houston, postdoctoral fellowship (PH), a Postdoctoral Fellowship in Translational Medicine by the PhRMA Foundation (MK), and an anonymous donation to establish the Stanford BSL3 facility. SE and CB are Chan Zuckerberg Biohub investigators. Use of the Stanford Synchrotron Radiation Lightsource, SLAC National Accelerator Laboratory, is supported by the U.S. Department of Energy, Office of Science, Office of Basic Energy Sciences under Contract No. DE-AC02-76SF00515. The SSRL Structural Molecular Biology Program is supported by the DOE Office of Biological and Environmental Research, and by grant P30GM133894 from the National Institute of General Medical Sciences of the National Institutes of Health.

## Author contributions

Conceptualization: MW, MZL

Methodology: MW, YS, XZ, AR, PH, JG, DF, LN, AB, MK, B-HP, SE, C-TKT, CB, MZL.

Investigation: MW, YS, XZ, AR, PH, JG, PBP, DF, YW, LN, AB, MK, CH, PS-H, VT, AD, B-HP.

Visualization: MW, MZL

Funding acquisition: MW, SE, CB, MZL

Project administration: MW, CB, MZL

Supervision: JG, SE, C-TKT, CB, MZL

Writing – original draft: MW

Writing – review & editing: MW, MZL

## Data and materials availability

All data are available in the main text or the supplementary materials. Nine co-crystal structures of SARS-CoV-2 M^pro^ complexed with ML1000, ML1001, ML1006m, ML1006a, ML2006a, ML2006a2, ML,2006a4, ML3006a, or ML4006a have been deposited in the Protein Data Bank with IDs: 7SET, 7SF1, 7SF3, 7U92, 8EZV, 8EZZ, 8F02, 8F2C, and 8F2D. Crystal structures of SARS-CoV-2 M^pro^ S144A complexed with ML1006a or NTV have been deposited with IDs: 7UUG and 7UUP.

