## Supplementary Materials for "Design of SARS-CoV-2 protease inhibitors with improved affinity and reduced sensitivity to mutations"

### Methods and Materials

Reference compounds

The following reference compounds were purchased from commercial sources: BPV (Cayman Chemical, ≥ 98%), NPV (AdooQ, ≥ 98%), TPV (AdooQ Bioscience, ≥ 98%), GC376 (AOBIOUS, ≥ 98%), ebselen (Cayman Chemical, ≥ 99%), disulfiram (LKT Laboratories, ≥ 98%), ritonavir for $M^{pro}$ inhibition (Santa Cruz Biotechnology ≥ 98%) or ADME and PK (MedChemExpress and Selleckchem, ≥ 99.9%; or Macklin, 98.55%), Ensitrelvir fumarate (Medkoo, > 99%), PF-00835231 (Selleckchem, 99.08%), NTV (Jinan Honest Pharm CO., > 99%)

Synthesis of ML inhibitors

ML1000 was synthesized by ACME Bioscience (Palo Alto, CA, USA). All other ML-series compounds were synthesized by Chempartner (Shanghai, China). NTV was initially synthesized on small scale by Chempartner (Shanghai, China). The synthesis procedures are described in a separate section at the end of this document.

$M^{pro}$ expression and purification

Modified versions of previous protocols based on HRV protease or SUMO protease (*1*) processing of $M^{pro}$ fusion proteins were used to obtain purified $M^{pro}$ with either native or extended termini after expression in *E. coli.* Specifically, $M^{pro}$ variants were cloned as I) GST-$M^{pro}$-$His_6$, II) $His_6$-SUMO-$M^{pro}$, or III) $His_6$-SUMO-$M^{pro}$-coil fusions into a pET vector (Addgene plasmid #29666) using synthetic gene blocks for the $M^{pro}$ portion of the SARS-CoV-2 polyprotein (pp1ab residue 3264–3569). $M^{pro}$ point mutants were constructed in both constructs II and III. Constructs details are shown in **Fig. S19**. In our hands, the $His_6$-SUMO-$M^{pro}$ fusions produced higher yields of soluble protein compared to the GST-$M^{pro}$-$His_6$ constructs. This is likely due to toxicity and growth retardation associated with $M^{pro}$ activation upon autocatalytic removal of the GST-tag during expression of the GST-$M^{pro}$-$His_6$ fusion. In contrast, the $His_6$-SUMO-$M^{pro}$ fusion is produced in full-length and only becomes fully active after SUMO-tag removal during subsequent purification steps.

For SARS-CoV-2 $M^{pro}$ variants, expression plasmids were transformed into T7 Express lysY/$I^q$ *E. coli* cells (NEB). Small overnight cultures in 2×YT media with 1% glycose were used to inoculate larger cultures in 2×YT media that were then grown at 37 °C to $OD_{600}$ ~ 0.8 before induction with 0.5 mM IPTG. After induction and 4–5 h growth at room temperature (RT, ~23 °C) the cells were harvested, and the pellets were frozen. Chemical lysis of the resuspended pellets was performed in B-PER (Thermo Scientific) supplemented with 40 U/mL Pierce universal nuclease (Thermo Scientific), and the supernatant cleared by centrifugation at 15000 g for 30 min. The soluble fraction was batch-absorbed onto INDIGO-Ni resin (Cube Biotech) in a buffer with 10 mM imidazole, 200 mM NaCl, and 2 mM DTT. The resin was loaded onto gravity flow columns and washed with 20 column volumes of wash buffer containing 50 mM Tris (pH 8.0), 25 mM imidazole, 300 mM NaCl, and 2 mM DTT. High purity protein was eluted in a buffer of 50 mM Tris (pH 8.0), 250 mM imidazole, 300 mM NaCl, and 2 mM DTT. Eluted fractions with high protein content were pooled and buffer exchanged into HRV protease cleavage buffer (50 mM Tris pH 7.3, 150 mM NaCl, 1 mM DTT) or SUMO protease cleavage buffer (50 mM Tris pH 8.0, 150 mM NaCl, 1 mM DTT). The proteins were cleaved by incubation with 1.5% (w/w) His-tagged HRV-3C protease (Millipore Sigma SAE0045) or 10 U/mg His-tagged SUMO protease (Millipore Sigma SAE0067) by overnight incubation at 4 °C. The fully processed $M^{pro}$ variants were then purified using reverse affinity chromatography to remove the His-tagged HRV and SUMO proteases and the cleaved $His_6$-tagged fusion-domains/peptides. Purity of the samples was checked on SDS-PAGE (>95%), and protein concentrations were determined based on $A_{280}$ and predicted extinction coefficients.

Enzyme characterization

The proteolytic activity of purified $M^{pro}$ was measured using Covidyte™ substrate peptides (AATBioquest) that incorporate either iFluor™ 670 (IF5) or Tide Fluor™ 5 (TF5) as a far-red fluorophore and Tide Quencher™ 5 (TQ5) as a quencher. Upon cleavage of IF5-VNSTLQ|SGLRK(TQ5)M (Covidyte IF670, **Fig. S2 and S3**) or TF5-KTSAVLQ|SGFRKME(TQ5)M (Covidyte TF670, all other data) by $M^{pro}$, energy transfer to the quencher decreases and the fluorescence intensity increases. A Safire 2 microplate reader (TECAN) was used to monitor the fluorescence with excitation at 640/20 nm and emission at 680/20 nm. All measurements were performed as bottom-reads from lid- or film-covered 96-well plates (Greiner; µclear bottom medium binding or non-binding). The in vitro $M^{pro}$ activity was measured in our general assay buffer consisting of 50 mM Tris (pH 7.3), 50 mM NaCl, 1 mM EDTA, 3 mM DTT (fresh), 0.05% (v/v) TWEEN20 and 1% (v/v) DMSO.

Michaelis-Menten parameters were obtained at 37 °C in the assay buffer. These assay conditions are comparable to the conditions used to obtain the apparent equilibrium inhibition constants, $K_{i,app}$ (*vide supra*). After 1 h of preheating of a well plate containing all solutions as 2× concentrated stocks, 45 µL of Covidyte TF670 substrate solution was transferred to 45 µL of protease solution and the fluorescence measurement immediately initiated on the microplate reader. The final concentration of protease was 50 nM, and the final concentration of substrate was 3–55 µM. The reaction was followed for 3–4 h at 37 °C. At the end of the measurement, the product formation had plateaued in all wells, and thus, all substrate molecules were assumed to have been consumed. Thus, to correct for inner filter effects, the end-point fluorescence signals were normalized to the known concentration of total substrate added to each well. The time-resolved product formation curves where fitted (MathWorks MATLAB) to the full analytical solution to the differential equations of the Michaelis-Menten scheme (*2*):

$$[P(t)] = 1 - K_m \cdot W\left\{\frac{[S(0)]}{K_m} \cdot \exp\left(\frac{[S(0)] - k_{cat}[E]_T(t-t_0)}{K_m}\right)\right\} \qquad \text{(Eq. 1)}$$

Here P is the product, S is the substrate, W is the Lambert W function, $K_m$ is the substrate binding constant, $k_{cat}$ is the zero-order rate constant of catalysis, $[E]_T$ is the total enzyme concentration, and $t_0$ accounts for a small delay in the first measurement after mixing the substrate and protease. During non-linear fitting, $[E]_T$ was fixed to 50 nM, [S(0)] was allowed to vary for the individual curves, and $K_m$, $k_{cat}$ were globally optimized across the substrate dilution series. The results are presented in **Fig. S20**.

Michaelis-Menten parameters could not be accurately determined for E166X mutants since the time-resolved product formation curves were not well-described by Eq. 1. Further, investigations are required to fully understand the kinetics of $M^{pro}$ E166X mediated proteolysis of Covidyte TF670.

In vitro $M^{pro}$ inhibition assays – $IC_{50}$

For initial $M^{pro}$ activity measurements, 150 nM $M^{pro}$ was preincubated with varying concentrations of inhibitor for 30 min at 30 °C (**Fig. S3**) before adding IF670 substrate to a final concentration of 1.8 µM. The procedure was subsequently improved during the development of the $M^{pro}$-coil assay. For the data reported in **Fig. 1, 2 and S4** the following procedure was used: Following 15 min activation in assay buffer at RT (~23 °C), 1.5 nM $M^{pro}$-coil was preincubated in 60 µL assay buffer with varying concentrations of inhibitor in 96-well plates at RT for 1 h. Next, 30 µL TF670 substrate in assay buffer was added to a final concentration of 3 µM, and the measurement immediately initiated. In later assays, reported in **Fig. 3, 6, S9, S13 and S17**, the temperature during both preincubation and read-out was increased to 37 °C and the preincubation time was increased to 3 h to allow the slow-binding ketoamides to reach equilibrium with $M^{pro}$-coil (**Fig. S21**). The other parameters were unchanged. When running the assay at 37 °C, the substrate solution was preheated before addition to the well plate.

The rate of substrate cleavage was extracted as the initial linear fluorescence increase and normalized against the rate of substrate cleavage in the absence of inhibitor, *I*, to obtain the relative activity, $v_{rel}$. From this, the $IC_{50}$ constants were obtained using a four-parameter logistic function (MathWorks MATLAB):

$$v_{\text{rel}} = v_{\text{max}} - \frac{(v_{max} - v_{min})}{1 + 10^{n_H\left(-pIC_{50} - \log([I]_T)\right)}} \qquad \text{(Eq. 2)}$$

The maximal and minimal rates of substrate cleavage, $v_{max}$ and $v_{min}$, $pIC_{50} = -\log(IC_{50})$, and the Hill coefficient, $n_H$ were all free fitting parameters in the non-linear fitting routine. The inhibitor concentration, $[I]_T$, is the total concentration of inhibitor during the preincubation.

Calculation of the thermodynamic inhibitory constant – $K_i$

The $IC_{50}$ data presented in **Figs. 3** and **S9** was further analyzed as many of the inhibition curves show the characteristics of tight binding (i.e., $IC_{50} \leq [M^{pro}\text{-coil}]$). $K_{i,app}$ values were extracted by fitting (MathWorks MATLAB) the relative protease activity, $v_{rel}$, as a function of total inhibitor concentration, $[I]_T$, to the Morrison equation (*3*):

$$v_{rel} = v_{max} - (v_{max} - v_{min}) \frac{([E]_T + [I]_T + K_{i,app}) - \sqrt{([E]_T + [I]_T + K_{i,app})^2 - 4[E]_T[I]_T}}{2[E]_T} \qquad \text{(Eq. 3)}$$

Here, $[E]_T$, is the total concentration of active enzyme (i.e., active $[M^{pro}\text{-coil}]_T$), and $K_{i,app}$ is the apparent inhibition constant. Since $M^{pro}$ is only active as a dimer and the exact fraction of active dimeric $M^{pro}$-coil is unknown, when working in the low nM range, $[E]_T$ was initially chosen to be a fitting parameter instead of fixed at 1.5 nM (i.e., the total concentration of $M^{pro}$-coil in solution). Specifically, $K_{i,app}$, $v_{max}$, and $v_{min}$ were allowed to take individual values for each dataset, while $[E]_T$ was treated as a global parameter for all fits (**Fig. S13**). The extracted concentrations of $M^{pro}$ active sites were 1.5 nM for both WT and S144A, respectively, in perfect agreement with the total concentration of 1.5 nM added $M^{pro}$-coil. This supports that the $M^{pro}$-coil proteins are present as active dimers even at low nM concentrations.

The datasets recorded for ML2006a titrated against both $M^{pro}$-coil WT and S144A were not well-described by Eq. 3 when using the $M^{pro}$-coil concentrations from the global fitting routine. The problem appears to arise from ML2006a being the tightest binding inhibitor, and even the high-quality data presented in **Fig. S13** does not have enough datapoints along the steep transition of the inhibition curve for this specific inhibitor. Thus, when fitting the data for ML2006a, $[E]_T$ was included as an individual fitting parameter without constraints from the global fit. From this procedure, $[E]_T$ was estimated to 1.1 nM and 1.0 nM for the WT and S144A $M^{pro}$-coil ML2006a datasets, respectively. These numbers are still in good agreement with the numbers reported above, but highlights the challenges in applying Eq. 3 when $K_{i,app} << [E]_T$.

For all mutants, other than S144A, $[E]_T$ was fixed at the preincubation concentration of added $M^{pro}$-coil since these datasets did not contain enough datapoints in the tight binding limit to obtain stable estimates of $[E]_T$ from the global fitting routine. This procedure also resulted in good fits (**Fig. S17**).

We assume that $K_i = K_{i,app}$ since 1) $[S] << K_m$ (e.g., $[S]$ = 3 μM vs. $K_m$ = 44 μM for WT $M^{pro}$-coil, **Fig. S20**) and 2) several of the tested inhibitors show slow dissociation kinetics which invalidates the Cheng-Prusoff correction. Specifically, the Cheng-Prusoff equation is derived based on the assumption that both the substrate and the inhibitor are in equilibrium with the enzyme during the assay (*4*). However, since the inhibitors are preincubated in the absence of substrate and only the initial rate of substrate proteolysis is used in our analysis, the inhibitors have not dissociated from active site and equilibrated with the substrate during fluorescence read-out. Effectively, we assume that the substrate does not compete with the $M^{pro}$ inhibitors during the short measurement time of this assay.

<u>Binding kinetics</u>

Inhibitor binding kinetics were investigated in a time-resolved association experiment at 37 °C. An inhibitor dilution series was prepared and premixed with Covidyte TF670 substrate in a 96-well plate. After preheating for 1 h at 37 °C, 60 μL of combined substrate and inhibitor premix was added to 30 μL of $M^{pro}$-coil WT solution, and recording of the progress curves of substrate proteolysis was immediately initiated on the microplate reader. The final concentrations in the assay buffer were 6 μM of substrate, 1 nM of $M^{pro}$-coil WT, and inhibitor concentration as annotated in **Fig. S14**. The kinetics were monitored for 1 h, and the progression curves for each inhibitor were globally fit (MathWorks MATLAB) to a simple two-state reversible binding model (*3*):

$$I_t = v_s(t - t_0) + \frac{(v_i - v_s)}{k_{off} + k_{on,app}[I]_T}\left(1 - \exp\{-(k_{off} + k_{on,app}[I]_T)(t - t_0)\}\right) \quad \text{(Eq. 4)}$$

Here, $v_i$ and $v_s$ are the relative initial and steady-state rates of product formation, $k_{on,app}$ and $k_{off}$ are the first and zero order rate constants of a 2-state binding model (i.e., E + I ↔ EI), and $t_0$ is included to account for the small time-delay of measurement initiation. During non-linear regression of each inhibitor dataset (**Fig. S14**), $v_i$, $k_{on,app}$, $k_{off}$, and $t_0$ were treated as global fitting parameters, while $v_s$ is a free fitting parameter at each concentration. The substrate independent association constant, $k_{on}$, was calculated as $k_{on,app} \cdot (1+[S]/K_m)$ to correct for the direct competition for $M^{pro}$ binding between the substrate and the inhibitor that are added simultaneously in this experiment (*3*) (**Table S3**). Using this method, the data was sufficiently described by the two-state binding model. More complicated binding models (e.g., preequilibrium induced fit) were also explored and cannot be ruled out based on current data. For example, an induced fit model with a very weak encounter complex (i.e., $K_{i,encounter} >> K_i$) is fully consistent with the data (*3*). However, the more advanced models did not improve the fit quality.

The kinetics of NTV were too fast for quantification at 37 °C. Thus, the experiment described above was also performed for NTV at 25 °C. The substrate concentration was 3 μM in this experiment, and all other conditions identical to the assay at 37 °C.

<u>Cathepsin inhibition assays</u>

Recombinant human Cat-B (R&D Systems, 953-CY), Cat-K (EMD Millipore, 219461), Cat-L (R&D Systems, 953-CY), or Cat-S (Sino Biological, 10487-H08H) were first activated for 15 min in 50 mM MES pH 5.5, 1 mM EDTA, 6 mM DTT. Next, inhibitors were added to a concentration of 15 μM and 1.5 μM and incubated with the protease (75/15/15/150 pg/mL Cat-B/K/L/S) in 50 mM MES pH 5.5, 1 mM EDTA, 3 mM DTT, 0.5% DMSO for 1 h at RT in 96-well plates (Greiner; µclear bottom medium binding or non-binding). After addition of the fluorogenic Z-LR-AMC substrate peptide (R&D Systems) to a final concentration of 20 µM, the reaction was immediately followed at RT using the Safire 2 microplate reader (TECAN) with excitation at 380/20 nm and emission detection at 460/20 nm. The initial substrate cleavage rate was determined by fitting a linear function to fluorescence signal. The remaining activity was obtained as the relative substrate cleavage rate relative to a 0.5% DMSO only control.

Crystallography

Mature SARS-CoV-2 $M^{pro}$ was buffer exchanged into a solution of 20 mM Tris pH 7.3, 2 mM DTT or 3 mM TCEP using Amicon Ultra-4 10K centrifugal filters (Millipore). Immediately after buffer exchange, $M^{pro}$ was incubated with inhibitor for a minimum of 1 h at RT before dispensing sitting drops on Intelli-plates 96-3 LVR (Hampton Research) using a Oryx8 crystallization robot (Douglas Instruments). The plates were incubated at 16 °C. Crystals formed under a range of conditions (**Table S5**) from either commercial screens or from custom optimization screens that were prepared with a Scorpion Screen Builder (Art Robbins Instruments). Crystals were harvested and added to precipitant solution mixed with cryoprotectant and cryocooled by plunging into liquid $N_2$.

In general, crystals from a single crystallization condition showed a wide variation in X-ray diffracting power, and therefore a large number of crystals were screened for initial data quality assessment. The best candidates were selected and stored for further data collection. In some cases, X-ray diffracting raster sampling was combined with microfocus (15 x 15 µm) to find the best diffracting region on the crystal, or alternatively microfocus was combined with helical data collection by sliding along the longest axis of the crystal. Data collections were performed at 100 K using Stanford Synchrotron Radiation Lightsource (SSRL) beamlines BL12-1 and BL12-2 (SLAC National Accelerator Laboratory, Menlo Park, USA) (*5*).

Data was reduced with XDS (*6*) or DIALS (*7*), scaled with SCALA (*8*) or AIMLESS (*9*) and analyzed with different computing modules within the CCP4 suite (*10*) or CCP4i2 suite (*11*). Crystals belonged to the orthorhombic $P2_12_12$ or monoclinic C2 space groups and contained one polypeptide chain per asymmetry unit (**Table S6**). Structures were solved by the molecular replacement method with Phaser (*12*) using the polypeptide chain of SARS-CoV-2 $M^{pro}$ of PDB 6LZE (*13*) as the search model. Refinement was performed using REFMAC (*14*) with manual refinement performed in COOT (*15*).

Cell-based $M^{pro}$ activity assays

*Cell culture and transfection.* HEK293A cells were cultured at 37 ºC in 5% $CO_2$ in Dulbecco's Modified Eagle's Medium (DMEM, Gibco 31053036) supplemented with 10% FBS (GeminiBio) and 100 U/mL penicillin + 100 µg/mL streptomycin (GeminiBio). Huh7 cells were cultured at 37 ºC in 5% $CO_2$ in Roswell Park Memorial Institute 1640 medium (RPMI 1640, Life Technologies) supplemented with 10% FBS and 100 U/mL penicillin and 100 µg/mL streptomycin.

*$M^{pro}$ activity assay in HEK293A using western blotting.* Cells were transfected with 100 ng of a pcDNA3.1/Puro-CAG plasmid containing the construct shown in **Fig. S2A** using Lipofectamine 3000 (Invitrogen) in Opti-MEM (Life Technologies) according to the manufacturer's protocol. TPV and BPV were added 2 h post-transfection. 24 h post-transfection, cells were washed twice with PBS, then lysed with 50 µL of 50% 4× LDS sample buffer (NuPAGE, Life Technologies) and 10% 2-mercaptoethanol, and DNA was sheared by sonication. After heating at 80–90 °C for 1–2 min, cell lysates were loaded onto 4–12% Bis-Tris gels (NuPAGE, Life Technologies) along with a Precision Plus protein dual-color standard (Bio-Rad). Protein bands were transferred to PVDF membranes using a Trans-Blot Turbo Transfer System (Bio-Rad) and blocked with SuperBlock T20 (TBS) blocking buffer (Thermo Scientific). Membranes were probed with primary antibodies in SuperBlock T20 (TBS) blocking buffer and fluorophore-conjugated secondary antibodies in SuperBlock (TBS) blocking buffer (Thermo Scientific), with 3 washes in SuperBlock T20 (TBS) blocking buffer after each step. Membranes were imaged using an Odyssey imaging system (LI-COR) and signals quantified using ImageJ. The following primary antibodies were used for immunoblotting at the indicated dilutions: mouse monoclonal anti-FLAG (Sigma-Aldrich, F1804), 1:2000; rabbit polyclonal anti-beta Actin (Abcam, ab8227), 1:5000. Secondary antibodies were LI-COR 680RD goat-anti-rabbit and 800CW goat-anti-mouse, used at 1:5000 dilution each.

*$M^{pro}$ activity assay in Huh7 with NanoBiT sensor.* A schematic of this assay is presented in **Fig. S22**. Huh7 cells were seeded as 10000 cells/well into solid white Lumitrac 96-well microplates (Greiner). 24 h later, cells were transfected with 10 ng reporter plasmid with 40 ng carrier DNA plasmid (pcDNA3.1 vector) using Lipofectamine 3000, following manufacturer's instructions. 24 h post-transfection, inhibitor dilutions in DMSO

(final concentration < 0.33%) were added to the cell culture. 48 h post-transfection, cells were lysed, and Nano-Glo Luciferase Assay System (Promega) added according to the manufacturer's instructions. After mixing the assay solution thoroughly, the luminescence signal was read using the Safire 2 microplate reader (TECAN) with an integration time of 1000 ms and a speed of 1 read/min. The maximum signal within the first 15 min was used for data analysis. The data was normalized against a DMSO control (low luminescence) and a full inhibition control (high luminescence) to obtain the relative luminescence intensity, $I_{rel}$, that correlates with intracellular $M^{pro}$ activity. The data was fit to Eq. 5 (MathWorks MATLAB) to obtain $IC_{50,cell}$:

$$I_{\text{rel}} = I_{\text{min}} + \frac{(I_{max} - I_{min})}{1 + 10^{n_H\left(-pIC_{50,cell} - \log([I]_T)\right)}} \qquad \text{(Eq. 5)}$$

Antiviral assays with SARS-CoV-2-NLuc

A549-ACE2 (A549+) and Huh7.5.1-ACE2-TMPRSS2 (Huh7.5.1++) cells (*16*, *17*) were maintained in D10 growth media (DMEM (Gibco 11885084) supplemented with 10% FBS (GeminiBio), 2 mM L-glutamine, 10 mM HEPES, 1 mM sodium puruvate, 1% non-essential amino acids, and 1% antibiotic-antimycotic (all supplements from Gibco)). In addition, the media was supplemented with 0.5 mg/mL geneticin (Gibco) and 0.2 mg/mL hygromycin (Gibco) for A549+ or Huh7.5.1++, respectively. SARS-CoV-2-NLuc (USA-WA1/2020 strain expressing NLuc) virus (*18*) was passaged twice in VeroE6 cells and titered by plaque assay on VeroE6 cells and confirmed by deep sequencing. The antiviral replication assay was performed in D2 assay media (D10 media adjusted to 2% FBS and without antibiotics).

Inhibitors were added in D2 media to A549+ or Huh7.5.1++ cells 24 h after plating the cells in solid white 96-well plates. Control wells were treated with equal concentrations of DMSO. Within 15 min of drug addition, cells were infected with virus at MOI = 0.1 for 2 h, washed, and incubated with drug for 48 h before addition of lytic Nano-Glo assay (Promega) and reading on a GloMax Discovery plate reader (Promega). Infections and plate reading occurred inside Class II biosafety cabinets under Biosafety Level 3 containment at Stanford. The plates always contained a positive inhibition control (e.g., GC376 or NTV), a DMSO control column as positive infection control, and a column without cells for background signal subtraction. The 10-base logarithm of the luminescent signal was background-corrected and normalized against the DMSO control to represent the relative viral replication rate. $EC_{50}$ values were obtained by fitting the relative viral replication rate as a function of the inhibitor concentration to Eq. 2 (MathWorks MATLAB) with $IC_{50}$ replaced by $EC_{50}$.

Antiviral plaque assay in Calu-3 cells

Calu-3 cells (human lung epithelium, ATCC) were grown in medium (DMEM; Gibco) supplemented with 10% fetal calf serum (FCS; Omega Scientific Inc.), 1% Pen-strep (Gibco) and 1% nonessential amino acids (NEAA, Gibco). Cells were maintained in a humidified incubator with 5% $CO_2$ at 37 °C.

Calu-3 were pretreated with $M^{pro}$ inhibitors and infected with SARS-CoV-2 (USA-WA1/2020) at MOI = 0.1 in DMEM containing 2% FCS. After 1 h incubation at 37 °C, the viral inoculum was removed, cells were washed and supplemented with new medium containing inhibitors. Culture supernatants were harvested at 24 h postinfection and the viral titers were measured via plaque assays.

For plaque assays, monolayers of Vero E6 cells were infected with supernatants from Calu-3 cultures for 1 h at 37 °C. Cells were then overlaid with MEM and carboxymethyl cellulose and incubated at 37 °C. At 72 h postinfection, the overlay was removed, and the cells were submerged in 70% ethanol for fixation and viral inactivation followed by crystal violet staining. Viral titers (PFU/mL inoculum) were quantified according to standard methods (*19*) and normalized against a DMSO only control group. $EC_{50}$ values were obtained by fitting (GraphPad Prism 9) the relative viral titers to Eq. 2 with $v_{min}$ and $v_{max}$ set to 0% and 100%, respectively and $IC_{50}$ replaced by $EC_{50}$.

Cytotoxicity assays in Huh7, A549-ACE2, and SARS-CoV-2 infected Calu-3 cells

24 h prior to the inhibitor treatment, 10000 Huh7 cells (RPMI with 10% FBS, 100 U/mL penicillin, and 100 µg/mL streptomycin) or A549+ cells (D10 media) were seeded in 100 µL culture medium in solid white 96-well plates. The next day, the culture medium was replaced with fresh media containing inhibitors at the desired concentration (1–100 µM). Staurosporine (0.1 µM or 1 µM), a non-selective protein kinase inhibitor known to induce apoptosis, was used as a positive control. After 72 h, cell viability was determined using the CellTiter-Glo 2.0 kit (Promega, USA) according to the instructions of the manufacturer. The bioluminescence signal was measured on the Safire 2 microplate reader (TECAN).

Cell viability was also assessed in Calu-3 cells infected with SARS-CoV-2 using alamarBlue (Invitrogen) according to the manufacturer's protocol. Fluorescence was detected at 560 nm on a GloMax Discover microplate reader (Promega).

ADME

All in vitro ADME experiments were performed at Shanghai Chempartner. All concentration measurements were performed by LC-MS/MS. All measurements were performed in technical duplicates (n = 2).

The apparent permeability, $P_{app}$, was measured as diffusion of 10 µM compound across Caco-2 cell layers in both the apical-to-basal (A-B) and the basal-to-apical (B-A) direction over a period of 90 min. The assay was performed in HBSS at 37 °C. Quality control was performed using the following reference compounds: erythromycin, metoprolol, atenolol, lucifer yellow. To improve comparison of $P_{app}$ measured in independent assays, $P_{app}$ was normalized to both erythromycin and atenolol in each assay, and the absolute mean $P_{app}$ calculated after multiplying the following mean reference values (in units of $10^{-6}$ cm/s ± SD): atenolol: $P_{app,A\text{-}B}$ 0.53 ± 0.34, $P_{app,B\text{-}A}$ = 0.73 ± 0.38; erythromycin: $P_{app,A\text{-}B}$ = 0.25 ± 0.20, $P_{app,B\text{-}A}$ = 18.6 ± 4.6.

Kinetic solubility was measured after adding compound to a final concentration of 100 µM in PBS (pH 7.4, 1% DMSO) and incubating with shaking for 1 h at RT. Precipitate was spun down before measuring the dissolved concentration. Propranolol, ketoconazole, and tamoxifen served as controls.

Plasma protein binding was measured in either 100% or 10% C57BL/6 mouse plasma or 100% human plasma after dialysis at 37 °C for 5 h. Plasma was diluted in in 0.05 mM sodium phosphate buffer (pH 7.4). The initial compound concentration was 1 µM (0.2% DMSO). Warfarin and Quinidine served as reference compounds. The samples were quenched with MeCN. The recovered fraction was > 70% in all cases, except for ML2006a4 (46% in 10% mouse plasma and 68% in 100% human plasma). The reduced recovery for ML2006a4 is not expected to affect the accuracy of the results (*20*). From measurements in 10% plasma, the fraction of unbound compound, $f_{u,p}$, at 100% plasma concentration was extrapolated using the method described by Di et al. (*21*).

Plasma stability at 37 °C was measured in a solution consisting of 90% CD-1 mouse or human plasma and 10% 0.05 mM sodium phosphate buffer (pH 7.4) containing 0.5% BSA. The starting concentration of the compounds were 2 µM (0.4% DMSO), and samples were collected and quenched with MeCN at 0, 5, 15, 30, 45, and 60 min. Linear regression of ln([compound]) vs. time was used to determine the half-life, $t_{1/2,p}$. Procaine was used to check plasma activity.

Microsomal metabolic stability of the compounds was measured in 0.5 mg/mL CD-1 mouse (Corning or Xenotech) or human liver microsomes (Corning) for 45 min at 37 °C in 0.1 M potassium phosphate buffer (pH 7.4) with 1.0 mM EDTA. The starting concentration of the compounds was 1 µM (0.1% MeCN). For a second set of measurements, 1 µM RTV was added as a CYP3A inhibitor (*22*). The reactions were initiated by the addition of 6 mM NADPH, and samples were quenched with MeCN at 0, 5, 14, 30, 45 min. Linear regression of ln([compound]) vs. time was used to determine the half-life, $t_{1/2}$, and the related apparent intrinsic clearance: $CL_{int,app} = \ln(2)/(t_{1/2} \cdot C_{microsome})$. Testosterone was used for quality control.

Pharmacokinetics using a saline PEG300/DMSO/Tween-80 (PDT) formulation

Single dose pharmacokinetics were performed at Shanghai Chempartner for ML1006m, ML1006a, ML2006a, ML2006a4, NTV, and PTV in male 6–8-week-old non-fasted C57BL/6 mice. Each measurement series consisted of 9 mice and 9 time points (pre-dose, 0.08, 0.25, 0.5, 1, 2, 4, 8, and 24 h). The mice were sampled in groups of 3, and each mouse was used for sparse sampling at three interspersed time points (*N = 3/timepoint*). At each timepoint ~110 µL blood was collected into $K_2$EDTA tubes via facial vein bleeding. Plasma samples were put on ice and centrifuged to obtain plasma sample (2000 g for 5 min at 4 °C) within 15 min of collection. The plasma samples were stored at -70 °C until analysis. At the terminal point of each series (i.e., 4, 8, or 24 h) both lungs were collected from all mice. No abnormal clinical signs were observed during the studies. The lung samples were homogenized with 3 volumes (v/w) of PBS. Compound concentrations were quantified at the end of the experiment using LC-MS/MS detection. The PK parameters were calculated from the mean plasma concentration of the pooled data of each group (*N =3*) versus time using a non-compartmental analysis (NCA) implemented in PKSolver software with linear trapezoidal integration (*23*). Parameter uncertainty estimates were obtained by two separate non-compartmental analyses performed on mean pooled data ± SD.

All compounds were formulated as colorless clear solutions in 45% saline, 40% PEG300, 10% DMSO, 5% Tween-80 (PDT). Four groups were tested. In group 1, oral doses (po, 20 mg/kg) were administered at 2.0 mg/mL, while in group 2, intravenous doses (iv, 2.0 mg/kg) were administered at 0.4 mg/mL. In groups 3 and 4, po dosing of 20 mg/kg RTV was performed 30 min before po or iv dosing of $M^{pro}$ inhibitors at the same doses as group 1 and 2, respectively. The RTV dose was set based on allometric scaling of a typical human PK enhancing RTV dose of 100 mg (*24*). Thus, for a human dose of 1.4 mg/kg (assuming mean body mass of 60 kg) the equivalent dose in mice is ~20 mg/kg based on a allometric scaling factor of 12.3 (*25*).

Pharmacokinetics and tolerability assessment using a methylcellulose/Tween-80 formulation

At Shanghai Chempartner, ML2006a4 and NTV were formulated together with RTV as co-suspensions in 0.5% methylcellulose and 2% Tween-80 in water (MCT-2). Pharmacokinetics associated with this formulation were monitored after oral administration of single doses of ML2006a4+RTV or NTV+RTV (po, 20+20 mg/kg at 5 mL/kg or 40+20 mg/kg at 10 mL/kg) in 6–8-week-old male C57BL/6 mice ($N = 3$). The study was performed as described for the PDT formulation, except lungs were not collected.

For the dose tolerability study, four groups of 8–12-weeks-old female C57BL/6 mice ($N = 6$) received oral doses at 10 mL/kg twice daily (b.i.d.) for 4 days. The four groups were 1) vehicle-only control, 2) RTV (20 mg/kg), 3) ML2006a4+RTV (40+20 mg/kg), and 4) NTV+RTV (40+20 mg/kg). The mice were observed twice a day and weighed once a day. At the end of the study, blood was collected 12 h after administration of the final dose and used to run standard clinical chemistry and hematology panels.

Pharmacokinetics of high dose single agent ML2006a4 and NTV

At Shanghai Chempartner, ML2006a4 and NTV were formulated as suspensions in 0.5% methylcellulose and 5% Tween-80 in water (MCT-5). Pharmacokinetics were monitored after oral administration of single doses of ML2006a4 or NTV (po, 300 mg/kg at 30 mL/kg) in 6–8-week-old male C57BL/6 mice ($N = 3$). Plasma samples were collected at 0.5, 1, 2, 4, 8, and 12 h. Otherwise, the study was performed as described above but without lung collection at terminal.

Antiviral efficacy in mice

The antiviral efficacy experiments were performed under BSL3 containment at Galveston National Laboratory at University of Texas Medical Branch at Galveston, Texas (UTMB). Animal studies were conducted based on a protocol approved by the Institutional Animal Care and Use Committee at UTMB. A total of 40 female BALB/c mice aged 15–16 weeks (Jackson Laboratory, 000651) were randomly separated into 5 groups for the study. Groups 1) RTV (20 mg/kg), 2) NTV+RTV (40+20 mg/kg), 3) ML2006a4+RTV (40+20 mg/kg) ($N = 10$) were anesthetized with isoflurane and challenged with $10^5$ $TCID_{50}$/60 µL of SARS-CoV-2 MA10 (*26*) via intranasal (IN) route. Mice of group 4) NTV+RTV (40+20 mg/kg) and 5) ML2006a4+RTV (40+20 mg/kg) ($N = 5$) were not challenged with virus and served as control groups to test the tolerability of the indicated test articles. Starting 2 h post infection, mice were treated with the indicated test articles twice daily (b.i.d.) for 4 days. Mice were weighed daily and, at least once daily, clinically observed and scored based on a 1–4 grading system (1: Healthy, 2: ruffled fur, lethargic, 3: hunched posture, orbital tightening, increased respiratory rate, and/or > 15% weight loss, and 4: dyspnea and/or cyanosis, reluctance to move when stimulated or > 20% weight loss). Mice with a clinical score of 4 were immediately euthanized and their lungs collected. Otherwise, five mice in group 1, 2 and 3 were euthanized as planned at 2 dpi, and their lungs were collected for viral load quantification and histopathology assessments. Observation of the remaining five mice per group continued for up to 6 dpi before euthanasia and immediate lung collection. First, lungs were inflated and rinsed with PBS. Then, the inferior and post-caval lobes of the right lung were collected and frozen for infectious viral load quantification. Next, the superior and middle lobes of the right lung were collected and stored in RNAlater for quantification of viral RNA levels. Finally, the left lung was fixed in 10% buffered formalin in preparation for histology assessments.

Mouse adapted SARS-CoV-2 MA10 was propagated and titered in Vero E6 cells. Vero E6 cells (ATCC, CRL:1586) were grown in minimum essential medium (EMEM, Gibco) supplemented with penicillin (100 units/mL), streptomycin (100 µg/mL) and 10% fetal bovine serum (FBS). The SARS-CoV-2 MA10 was stored in EMEM supplemented with 2% FBS at -80 °C until needed.

For quantitation of viral titers in the lung tissue, the frozen lung specimens were weighed before homogenization in PBS/2% FBS solution using a TissueLyser (Qiagen), as previously described (*27*). After centrifugation at 3000 g, supernatants were used to quantify infectious virus titers in a standard Vero E6 cell-based infectivity assays in 96-well plates (*28*). Viral titers were expressed as 50% tissue culture infectious dose per gram of tissue ($TCID_{50}$/g). Statistical analysis of virus titers was performed in Prism 9 (GraphPad) as an ordinary one-way analysis of variance (ANOVA) with post-hoc Tukey multi comparison between all groups ($N = 5$ *per timepoint*). The adjusted p-values are presented in **Fig. 4F** together with the relevant data.

For RNA extraction and quantitative RT-PCR, lung tissues were weighed and homogenized in 1 mL of Trizol reagent (Invitrogen) using TissueLyser (Qiagen). The RNA was then extracted using Direct-zol RNA miniprep kits (Zymo research) according to the manufacturer's instructions. To quantify the viral copies, RNA was reverse transcribed using iScript cDNA Synthesis kits (Bio-Rad, 1708891) according to the manufacturer's instructions. The cDNA was then amplified using iQ SYBR green supermix (Bio-Rad) and a CFX96 real-time PCR detection system (Bio-Rad). Primer sets for the SARS-CoV-2 E gene (5'-ACAGGTACGTTAATAGTTAATAGCGT; 5'- ATATTGCAGCAGTACGCACACA) were used. The samples were run in duplicate using the following conditions: 95 °C for 3 min, then 45 cycles of 95 °C for 15 s and 58 °C for 30 s. The plasmid, 2019-nCoV_E Positive Control (IDT, 10006896) was included as a standard to quantify the absolute copies of viral RNA. Statistical analysis of RNA copy levels was performed in Prism 9 (GraphPad) as an ordinary one-way analysis of variance (ANOVA) with post-hoc Tukey multi comparison between all groups (*N = 5 per timepoint*). The adjusted p-values are presented in **Fig. 4G** together with the relevant data.

For histopathological evaluation, the formalin fixed lung tissues were processed by the Histopathology Core (UTMB) to prepare 5 µm paraffin sections stained with Hematoxylin and Eosin (H&E). The stained lung sections (*N = 5 per timepoint*) were evaluated by an experienced pathologist who was blinded to the treatment type. The sections were evaluated for pathologic changes, respiratory epithelial degeneration, and inflammatory infiltrations. Representative images were taken using a Zeiss microscope equipped with an EOS Rebel T3i Canon camera.

Hazards

No unexpected or unusually high safety hazards were encountered.

### Supplementary Text

#### *In vitro ADME*

In vitro absorption, distribution, metabolism and excretion (ADME) properties were tested for multiple compounds (**Table S1**). A main challenge identified in the in vitro ADME studies was low permeability with $P_{app,A\text{-}B} < 10^{-6}$ cm/s for all initial ketoamide-based compounds, which limits oral bioavailability and $EC_{50}$. Thus, increasing permeability was one of the primary objectives of the study.

Inspecting the data of the inhibitors with the highest structural similarity to NTV, the efflux ratios ($P_{app,B\text{-}A}$ /$P_{app,A\text{-}B}$) also indicate that several of the compounds are targets for efflux. ML1004m and ML1005m are interesting outliers with $P_{app}$ ratios < 2. ML1006m had a slightly increased $P_{app}$ ratio of 4.3, while efflux appears to be a concern for ML1006a and NTV with high $P_{app}$ ratios of 105 and 25, respectively. Thus, the trifluoromethyl P4 cap as well as the azetidine substituted ketoamide and nitrile warheads appear to increase efflux. Metabolic stability was tested in human and mouse liver microsomes. In human microsomes, ML1004m through ML1006m showed high stability with intrinsic clearance rates of < 5 µL $min^{-1}$ $mg^{-1}$, while ML1006a and NTV underwent faster degradation at 13 and 18 µL $min^{-1}$ $mg^{-1}$, respectively. Since peptidomimetics including BPV are known targets of CYP3A degradation (*29*, *30*), the pharmacokinetic enhancer ritonavir (RTV) was added to inhibit CYP3A (*22*, *31*). After addition of RTV, the clearance rate of ML1006a and NTV dropped to < 5 µL $min^{-1}$ $mg^{-1}$, confirming these compounds as targets of CYP3A degradation.

While the intracellular $IC_{50,cell}$ and $EC_{50}$ assays had shown improved permeability of ML1006a (**Fig. 2C**) relative to ML1006m, this trend is surprisingly reversed in the $P_{app}$ data. In contrast, ML2006a, ML2006a2, ML2006a4, ML3006a, and ML4006a all showed clear improvements in $P_{app,A\text{-}B}$ relative to ML1006a consistent with the trends seen in the $IC_{50}/IC_{50,cell}$ ratios (**Fig. 3C**). Particularly, the $P_{app,A\text{-}B}$ values of the ML2006a2 and ML2006a4 compounds are now comparable to BPV. Similar to NTV, ML2006a4 is also cleared by a CYP3A mechanism since the intrinsic clearance rate of 77 µL $min^{-1}$ $mg^{-1}$ in human microsomes is efficiently reduced to <5 µL $min^{-1}$ $mg^{-1}$ by the addition of 1 µM RTV. In contrast to ML2006a4 and NTV, the microsome-mediated metabolism of PTV is not fully inhibited by the addition of RTV.

#### *Repeated dose tolerability assessment*

To test and compare the in vivo tolerability of ML2006a4 and NTV, mice were dosed twice daily (b.i.d.) with ML2006a4+RTV or NTV+RTV for 4 days, and clinical, hematological, and biochemical evaluations were performed by comparing four groups: 1) vehicle-only, 2) RTV (20 mg/kg), 3) ML2006a4+RTV (40+20 mg/kg), and 4) NTV+RTV (40+20 mg/kg). In all groups, similar body weight gains were observed (**Fig. S12C**) and no clinical signs were registered. Full hematology and clinical chemistry reports are presented in **Table S7 and S8**. When comparing group 2–4 against group 1, no patterns of adverse effects were identified in the hematology dataset. Single observations in group 3 and 4 showed reduced platelets but within the lower limit of reference values (*32*). However, the clinical chemistry dataset showed substantial variation in select groups and parameters. Thus, this dataset was further analyzed by annotating large deviations defined as values outside the interval of $mean_{vehicle} \pm 3SD_{vehicle}$. Specifically, 3 mice in the NTV+RTV treated group 4 showed clear deviations in multiple biochemical markers of liver and renal function, and particularly increased levels of ALT (3.9-7.6× baseline), AST (2.6-9.6× baseline), LDH (2.6-4.5× baseline), CK (3.2-3.8× baseline), and CREA (1.4-1.5× baseline). Reversible liver changes and increased levels of AST and ALT have previously been reported when dosing NTV alone in rats or monkeys, but have not been observed in human clinical studies (*33*). Group 2, with RTV only dosing, also had slightly increased values of in AST and LDH in 3 mice, however, the magnitude was not as pronounced as for the NTV+RTV in group 4. Depending on the dose, RTV has in rare cases been associated with increased levels of aminotransferase levels and hepatotoxicity in humans (*34*, *35*). In contrast, ML2006a4+RTV dosing did not cause any systematic deviations in clinical chemistry relative to vehicle dosing.

### Supplementary Figures

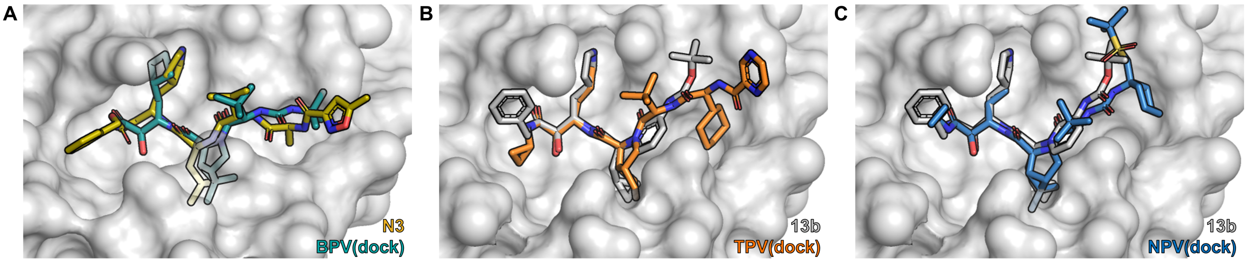


**Fig. S1. HCV protease inhibitors manually docked into SARS-CoV-2 M$^{pro}$.**
(**A**) Overlay of boceprevir (BPV, cyan) dock from **Fig. 1A** onto the co-crystal of SARS-CoV-2 M$^{pro}$ and the inhibitor N3 (yellow) (PDB 6LU7). (**B**) Telaprevir (TPV, orange) could also be manually docked with good complementarity in its P1 and P2 groups, with a different P2 proline analog than in BPV (Fig. 2A), whereas its P4 group appeared slightly too large for the S4 pocket. (**C**) Narlaprevir (NPV, blue) also demonstrated complementary in its P1 and P2 groups, which are identical to BPV, but its P4 group was clearly too large for the S4 pocket. Co-crystal structures confirming the overall poses have since also been published for TPV (e.g. PDB 7K6D and 6XQS) and NPV (e.g. PDB 7D1O, 6XQT, and 7JYC) (*36–38*). The manual docking was in all instances performed against 13b (grey) bound in M$^{pro}$ (PDB 6Y2G, protomer B).

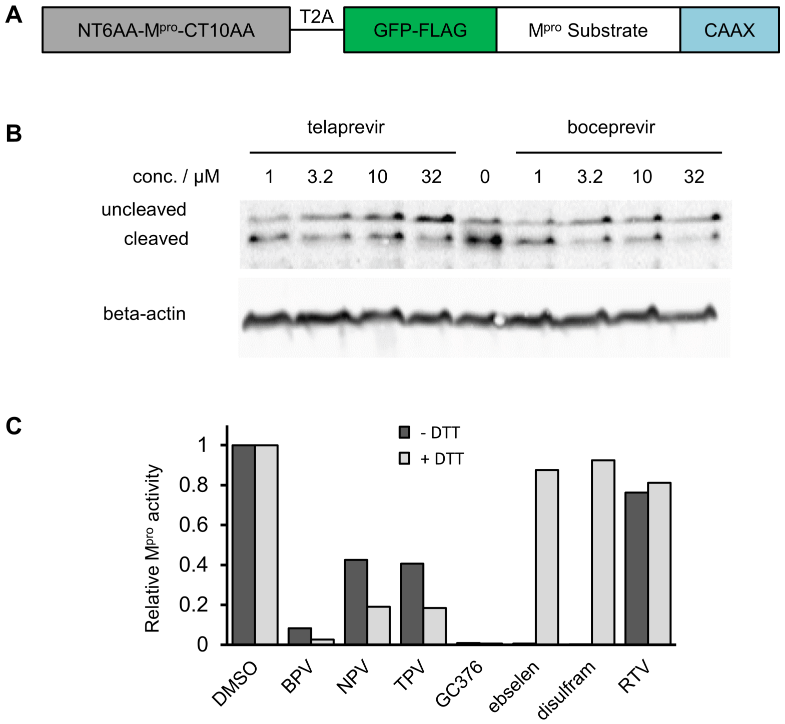


**Fig. S2. Pilot experiments assessing SARS-CoV-2 M$^{pro}$ activity in cells.**

(**A**) Schematic of the construct for co-expressing a SARS-CoV-2 M$^{pro}$ precursor and a substrate protein. The M$^{pro}$ precursor is designed to autocatalytically cleave at the natural N- and C-terminal substrate sequences to produce mature M$^{pro}$, recapitulating its natural maturation process. The substrate comprises FLAG-tagged green fluorescent protein (GFP), a substrate site, and the membrane-targeting CAAX sequence, so that cleavage at the substrate site liberates GFP from the membrane. (**B**) After transfection in HEK293A, addition of BPV or TPV can inhibit cleavage at micromolar concentrations, as assessed by anti-FLAG immunoblotting. (**C**) Initial screening of inhibitory capacity of a focused set of approved and investigational drugs with preincubation of 225 µM inhibitor and 150 nM SARS-CoV-2 M$^{pro}$-His$_6$. His-tagged M$^{pro}$ was initially used due to ease of production and purification. BPV inhibits M$^{pro}$ under reducing conditions, while ebselen and disulfram do not inhibit in the presence of 2 mM DTT. These results are consistent with proposals of disulfiram and ebselen forming a reducible bond with M$^{pro}$, making them non-optimal inhibitors of M$^{pro}$ in the reducing intracellular environment (*39*, *40*).

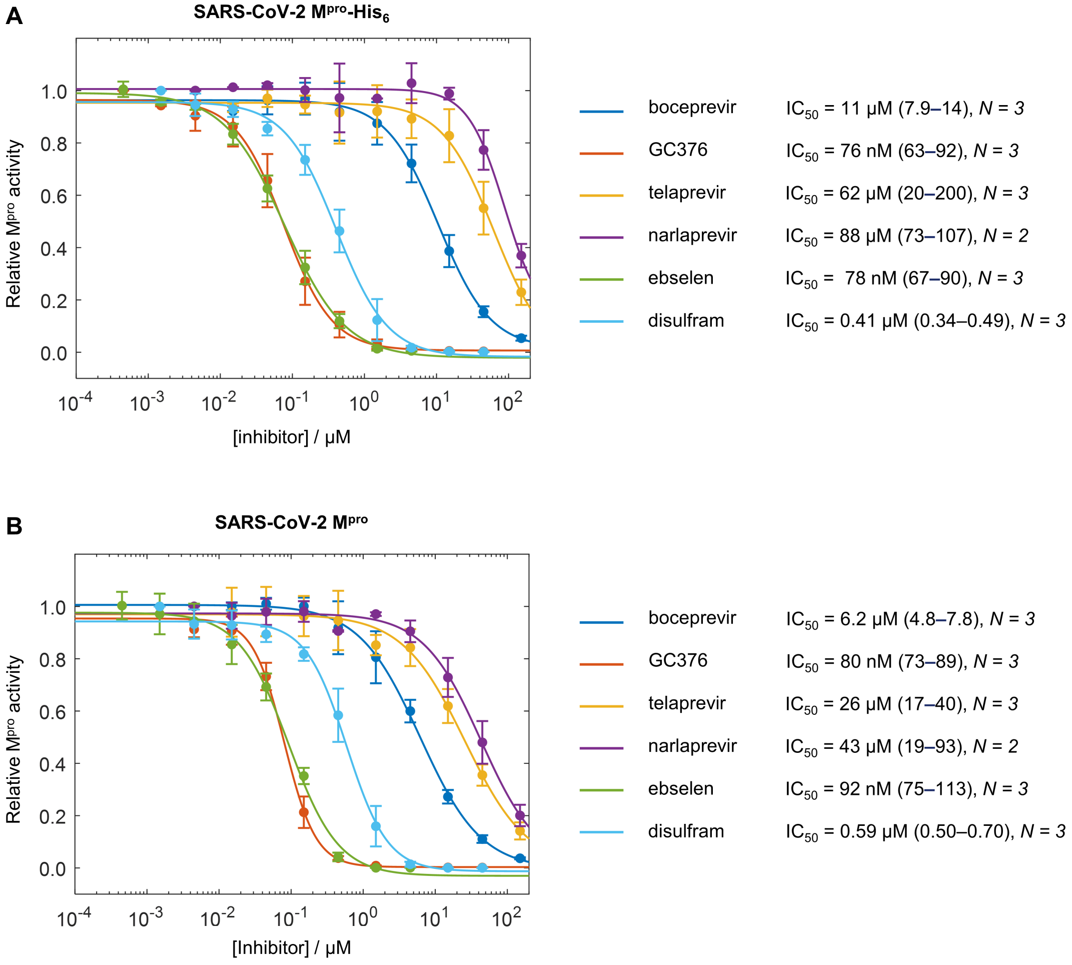


**Fig. S3. Initial M$^{pro}$ inhibition curves.**
IC$_{50}$ measurements by inhibitor titrations on SARS-CoV-2 (**A**) M$^{pro}$-His$_6$ or (**B**) M$^{pro}$, without the His tag. M$^{pro}$ activity was measured after preincubation of 150 nM SARS-CoV-2 M$^{pro}$-His$_6$ or M$^{pro}$ with inhibitors for 30 min at 30 °C followed by reaction with 1.8 µM of Covidyte IF670 substrate at 30 °C. We tested drugs in the presence of 2 mM DTT, except for ebselen and disulfiram, for which we omitted DTT. GC376 was included as a known potent M$^{pro}$ inhibitor. BPV is the most potent of the HCV protease inhibitors. IC$_{50}$ values were extracted by aggregating all *N* datasets and performing non-linear regression. CI$_{95}$ of the fit is reported in parentheses.

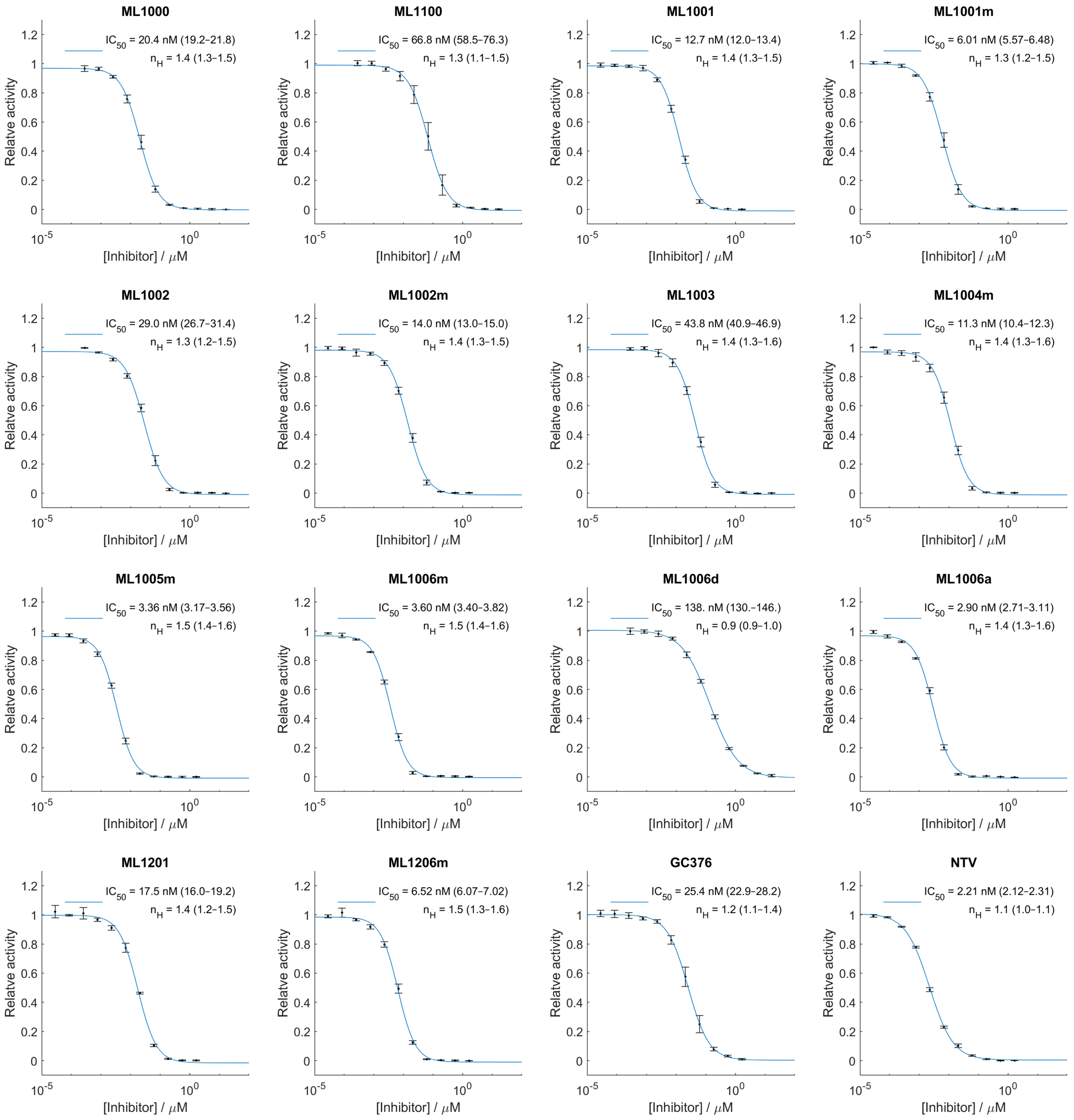


**Fig. S4. M$^{pro}$-coil inhibition curves at 25 °C.**
M$^{pro}$ activity was measured after preincubation of 1.5 nM SARS-CoV-2 M$^{pro}$-coil with inhibitors for 1 h at RT (~23 °C) followed by reaction with 3 μM of Covidyte TF670 substrate at 25 °C ($N = 3$). As discussed in Fig. S21, equilibrium has not been fully established for the ketoamide-based inhibitors under these conditions, so the measured $IC_{50}$ values are expected to be higher than their values at equilibrium. $IC_{50}$ values and Hill coefficients, $n_H$, were extracted by aggregating all $N$ datasets and performing non-linear regression. $CI_{95}$ of the fit is reported in parentheses. Error bars are SD.

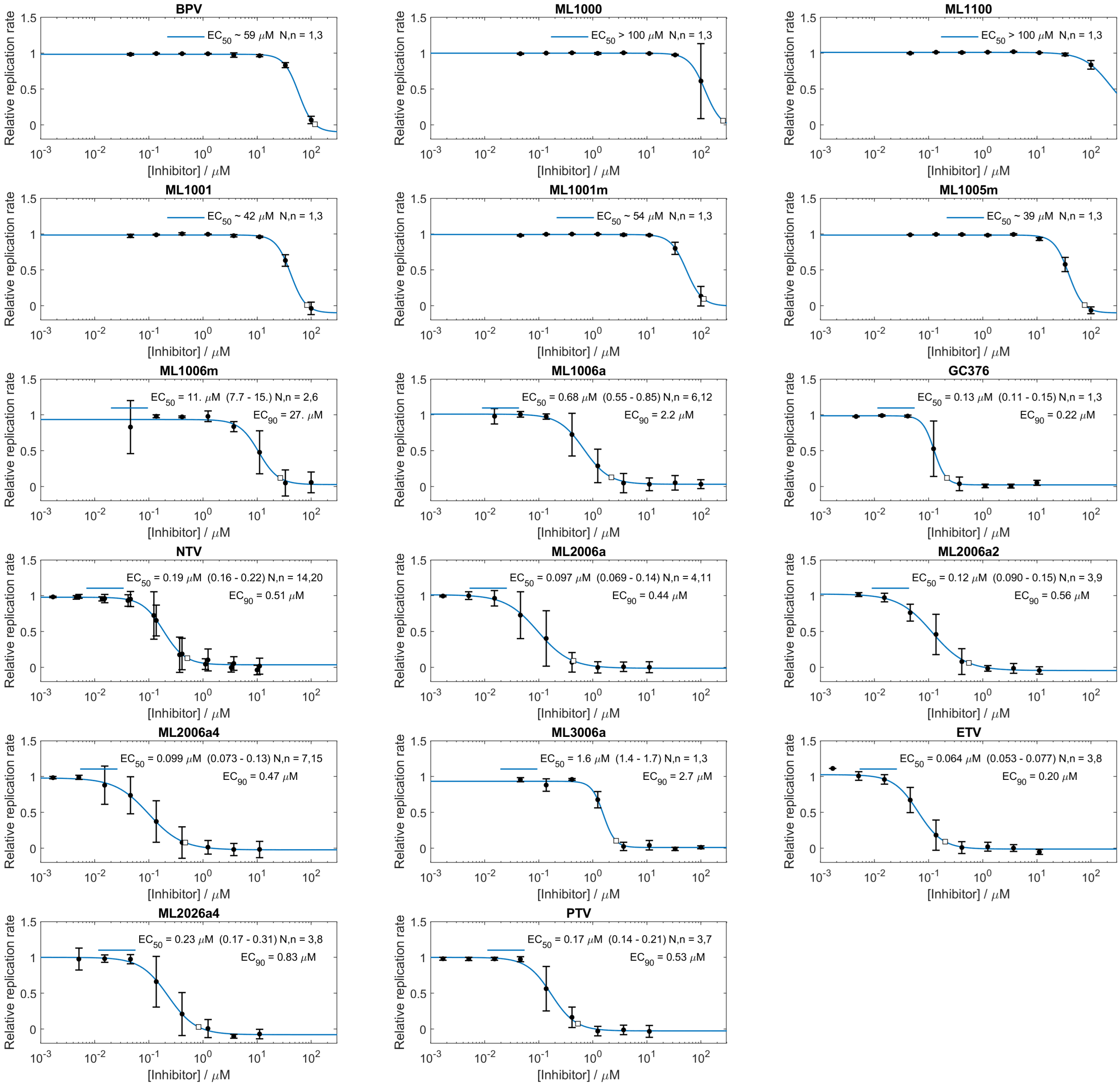


**Fig. S5. Inhibition of SARS-CoV-2-NLuc replication in Huh7.5.1-ACE2-TMPRSS2 cells.**
The luminescent virus signal was transformed into a relative replication rate as described in the methods section. Replicates are denoted as *N* for biological replicates performed on separates days, and *n* for total number of infection series. $EC_{50}$ values were extracted by aggregating all *n* datasets and performing non-linear regression. $CI_{95}$ of the fit is reported in parentheses. Error bars are SD. The $EC_{90}$ extracted from the fit is marked with a white square. For NTV, the concentrations of the dilution series were changed slightly between the independent experiments, giving rise to additional non-overlapping datapoints compared to the other datasets.

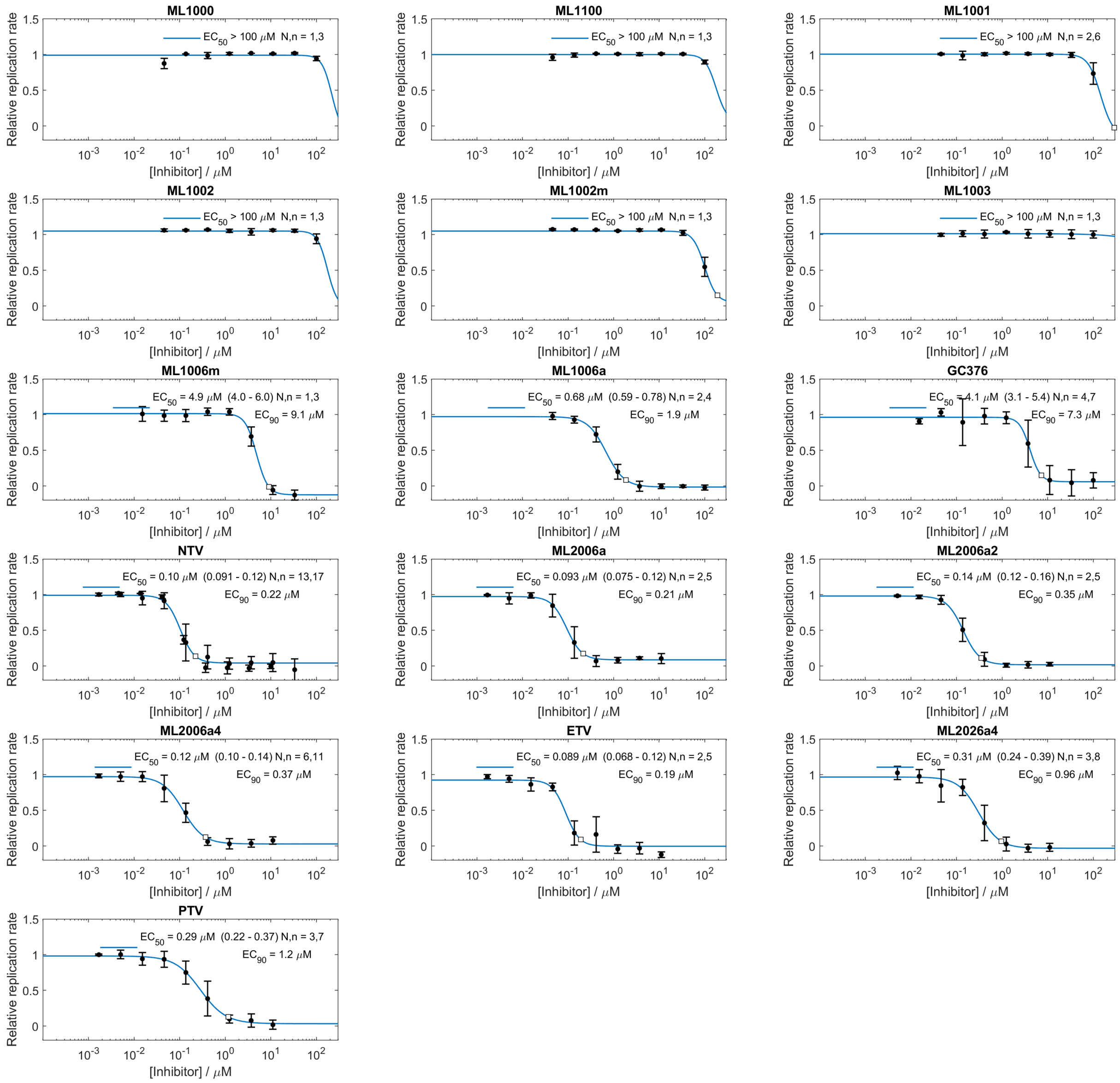


**Fig. S6. Inhibition of SARS-CoV-2-NLuc replication in A549-ACE2 cells.**
The luminescent virus signal was transformed into a relative replication rate as described in the methods section. The replicates are denoted as *N* for biological replicates performed on separates days, and *n* for total infection series. $EC_{50}$ values were extracted by aggregating all *n* datasets and performing non-linear regression. $CI_{95}$ of the fit is reported in parentheses. Error bars are SD. The $EC_{90}$ extracted from the fit is marked with a white square. For NTV, the concentrations of the dilution series were changed between the independent experiments, giving rise to additional non-overlapping datapoints compared to the other datasets.

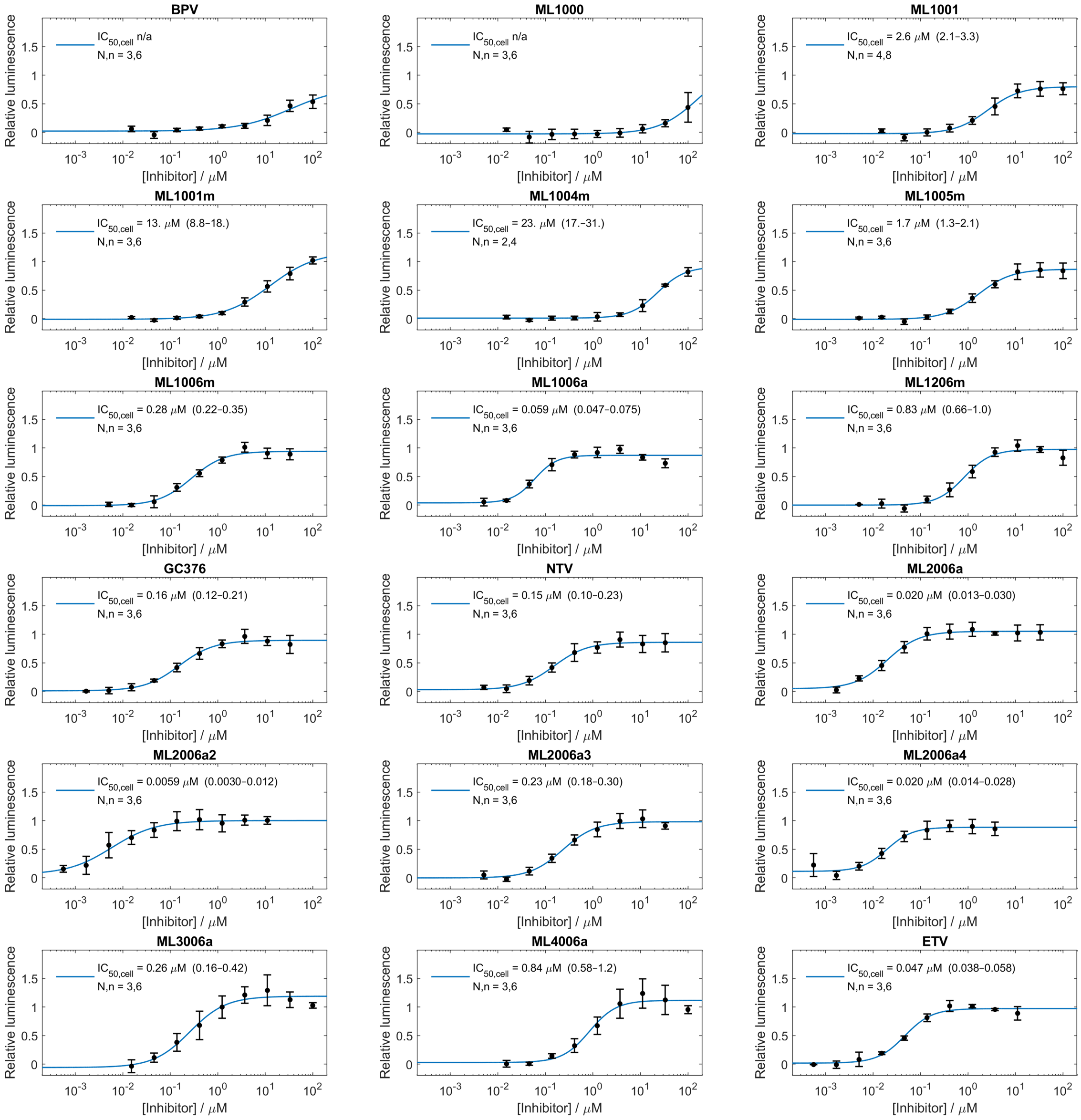


**Fig. S7. Intracellular SARS-CoV-2 M$^{pro}$ inhibition assay.**

Accumulation of the luminescent sensor is suppressed when M$^{pro}$ is active and builds up when M$^{pro}$ is inhibited (see schematic in Fig. S22). The replicates are denoted as *N* for biological replicates performed on separates days, and *n* for total number of replicates including replicates performed on the same plate. $IC_{50,cell}$ values were extracted by aggregating all *n* datasets and performing non-linear regression. $CI_{95}$ of the fit is reported in parentheses. Error bars are SD. In instances where the fitting routine could not accurately estimate the mid-point of the logistic function, $IC_{50}$ values are denoted as n/a.

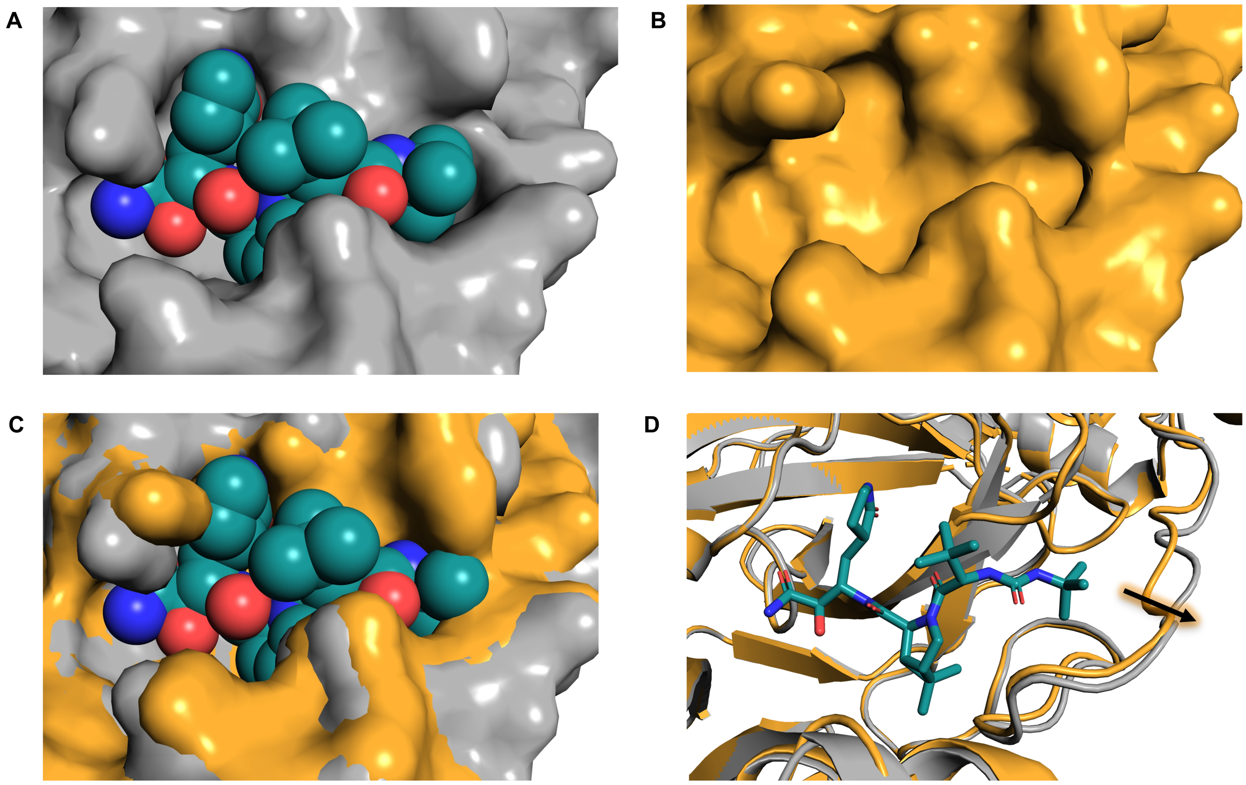

**Fig. S8. ML1000 binding widens the M$^{pro}$ binding pocket.**
(**A**) Surface and spheres representation of the SARS-CoV-2 M$^{pro}$ ML1000 co-crystal structure that is also presented in Fig. 1 and 2 of the main text. The P1-P4 groups appear to fit snuggly in the active site binding pocket. (**B**) SARS-CoV-2 M$^{pro}$ apo structure (PDB 6YB7). (**C and D**) Two representations of overlays of the structures from **A** and **B** indicate that the tertbutyl capped urea group of ML1000 stretches the S4 pocket relative to apo M$^{pro}$. The linker between domain II and III of M$^{pro}$ that defines this edge of the S4 pocket is known to be flexible (*41–44*).

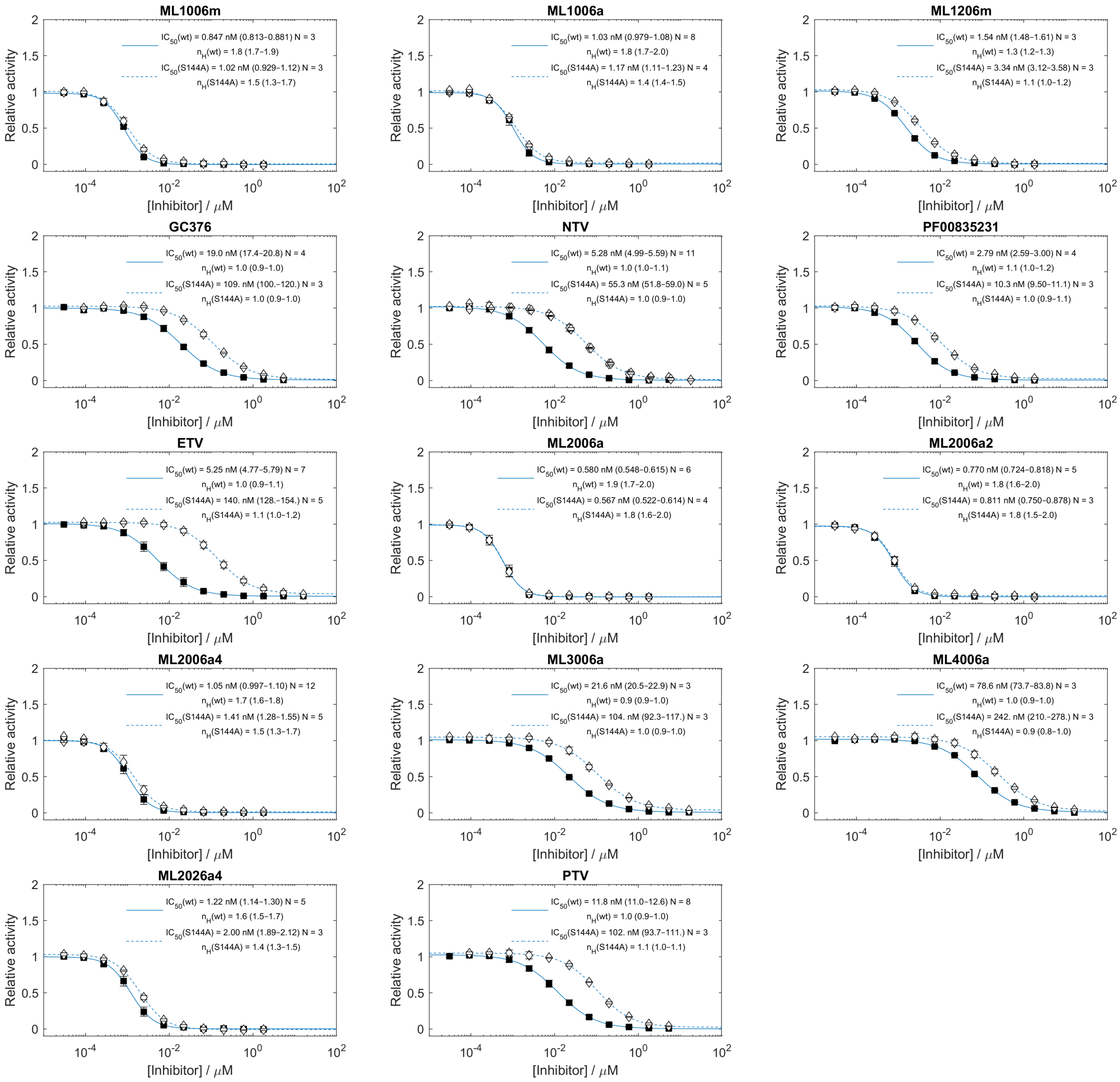


**Fig. S9**. **$M^{pro}$-coil WT and S144A inhibition curves at 37 °C.**
$M^{pro}$ inhibition curves obtained after preincubation of 1.5 nM SARS-CoV-2 $M^{pro}$-coil WT or S144A with inhibitors for 3 h at 37 °C followed by reaction with 3 μM of Covidyte TF670 substrate at 37 °C. Other than extended preincubation and increased assay temperature the assay was performed as described in the methods section and Fig. S4. $IC_{50}$ values and Hill coefficients, $n_H$, were extracted by aggregating all $N$ datasets and performing non-linear regression. $CI_{95}$ of the fit is reported in parentheses. Error bars are SD.

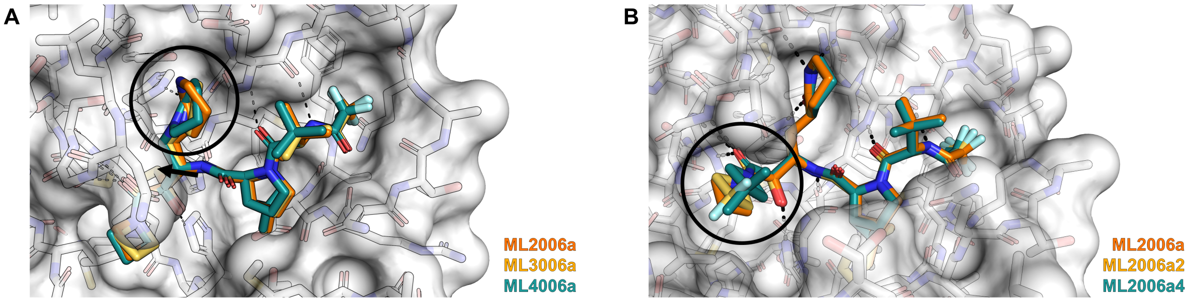


**Fig. S10**. **M$^{pro}$ Co-crystal structures of inhibitors with optimized P1 and P1' groups.**
(**A**) The overlay of ML2006a, ML3006a, and ML4006a with H-bonds shown for ML4006a (black) show that the inverted lactams of ML3006a and ML4006a fit in the S1 pocket and form a H-bond with H163. However, due to the planar nature of the inverted lactams, the rings are angled differently in the pocket compared to the δ-lactam of ML2006a. Furthermore, no HBD is available for interaction with the side chain of E166 and the backbone carbonyl of F140. Overall, ML3006a and particularly ML4006a appears to shift (arrow) slightly to minimize a potential steric clash with E166. **(B)** The overlay of ML2006a, ML2006a2, and ML2006a4 only shows a minor change in the angle of the azetidinyl in S1'. From the available structures it appears that the larger 3,3-dimethyl is slightly angled out of the S1' pocket to avoid steric clashes. For both figures only the surface contour of M$^{pro}$ from the ML2006a co-crystal is shown.

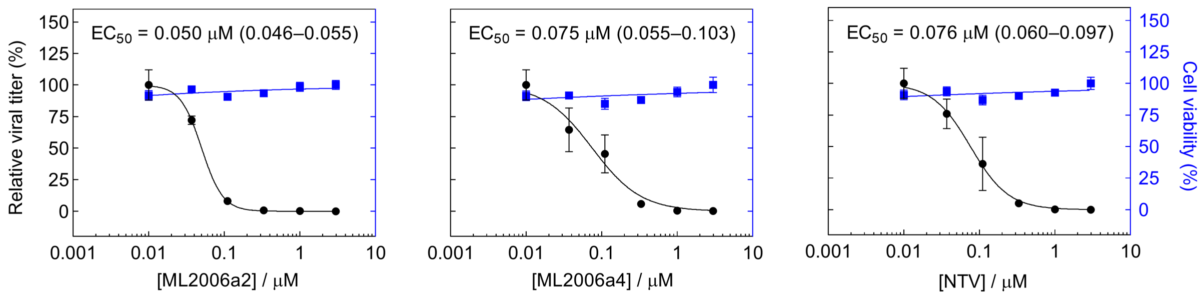


**Fig. S11**. **Inhibition of infectious SARS-CoV-2 production in Calu-3 cells.**
Dose response of SARS-CoV-2 infection (black, USA-WA1/2020 strain, MOI = 0.1, $N = 3$) and cell viability (blue, $N = 4$) in Calu-3 cells measured via plaque and alamarBlue assays at 24 h postinfection, respectively. $CI_{95}$ of the fit is reported in parentheses. Error bars are SD.

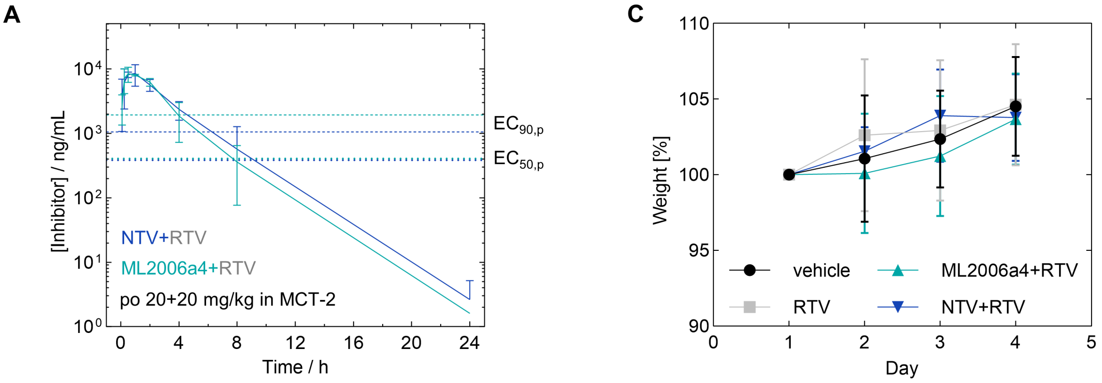


**B** **Oral pharmacokinetics in male non-fasted C57BL/6 mice using either PDT or MCT-2 formulations**

| | Formu-lation | Dose | $C_{max}$ | $C_{max,u}$[a] | $t_{max}$ | $AUC_\infty$ | $AUC_{\infty,u}$[a] | $t_{1/2}$ | $C_{max}$(RTV) |
|---|---|---|---|---|---|---|---|---|---|
| mouse | | [mg/kg] | [µM] | [µM] | [h] | [µg·h /mL] | [µg·h /mL] | [h] | [µg/mL] |
| **RTV[b] + ML2006a4** | PDT | 20 + 20 | 8.5 ± 0.5 | 1.30 ± 0.08 | 2.0 | 39 (32–46) | 6.0 (5.0–7.0) | 1.8 | 20 ± 14 |
| **RTV[c] + ML2006a4** | MCT-2 | 20 + 20 | 13 ± 4 | 2.05 ± 0.54 | 0.5 | 30 (20–39) | 4.5 (3.1–5.9) | 2.0 | 1.0 ± 0.5 |
| **RTV[c] + ML2006a4** | MCT-2 | 40 + 20 | 28 ± 2 | 4.27 ± 0.27 | 0.5 | 49 (38–60) | 7.5 (5.9–9.1) | 2.3 | 0.7 ± 0.2 |
| **RTV[b] + NTV** | PDT | 20 + 20 | 25 ± 3 | 6.12 ± 0.65 | 0.5 | 61 (51–72) | 15 (12–17) | 2.1 | 16 ± 2 |
| **RTV[c] + NTV** | MCT-2 | 20 + 20 | 17 ± 6 | 4.11 ± 1.52 | 1.0 | 32 (19–47) | 7.9 (4.6–11.4) | 2.0 | 1.0 ± 0.8 |
| **RTV[c] + NTV** | MCT-2 | 40 + 20 | 41 ± 10 | 9.96 ± 2.47 | 0.5 | 44 (28–59) | 10.5 (6.8–14.2) | 2.3 | 0.4 ± 0.4 |
| human[d] | | mg | [µM] | [µM] | [min] | [µg·h /mL] | [µg·h /mL] | [h] | [µg/mL] |
| **RTV + NTV** (fed) | tablet | 100 + 250 | 6.7 | 1.9 | 4.0 | 28 | 8.3 | 6.0 | – |
| **RTV + NTV** (fasted) | tablet | 100 + 250 | 5.8 | 1.7 | 2.75 | 29 | 8.2 | 6.9 | – |

**Fig. S12**. **Human-equivalent dosing and 4-day repeated dose tolerability study.**
(**A**) Plasma concentrations were monitored for 24 h after administration of MCT-2 formulated co-suspensions of 20+20 mg/kg ML2006a4+RTV or NTV+RTV. The equivalent data at the 40+20 mg/kg dosing level is presented in Fig. 5A. $EC_{50,p}$ and $EC_{90,p}$ were measured in Huh7.5.1++ cells. (**B**) Oral pharmacokinetic parameters were extracted by non-compartmental analysis of the data presented in panel A and Fig. 4B and 5A. Dosing with 40+20 mg/kg co-suspensions of ML2006a4+RTV and NTV+RTV in MCT-2 was chosen to simulate the human equivalent dosing regimen of Paxlovid for the mouse safety and efficacy experiments. While the human equivalent dose was primarily chosen by comparing $AUC_{\infty,u}$ this simple comparison does not take into account the temporal characteristics. Specifically, $t_{1/2}$ was substantially shorter for NTV in our mouse experiments compared to the $t_{1/2}$ observed in humans. Thus, in contrast to the situation in humans (*45*), the plasma concentrations periodically drop below $EC_{90,p}$ during the b.i.d. treatment protocol used in the mouse efficacy experiments (Fig. 5A-B). In parentheses, parameter intervals calculated using non-compartmental analysis of the plasma concentration mean ± SD (N = 3). For direct experimental observations, ± SD are reported. [a]Calculated using $f_u$ reported in Table S1. [b]RTV dosed po 30 min prior to test compound. [c]RTV dosed in a co-suspension with the $M^{pro}$ inhibitor. [d]The PK data for a Paxlovid comparable treatment regimen was extracted from the human phase I study of NTV (*45*), although, the approved Paxlovid dosing regimen uses 300 mg NTV instead of 250 mg (*46*). (**C**) The daily body weight of 8–12-weeks-old female C57BL/6 mice (*N* = *6*) were comparable for all groups upon twice daily dosing for 4 consecutive days with MCT-2 formulated co-suspensions of 40+20 mg/kg ML2006a4+RTV or NTV+RTV.

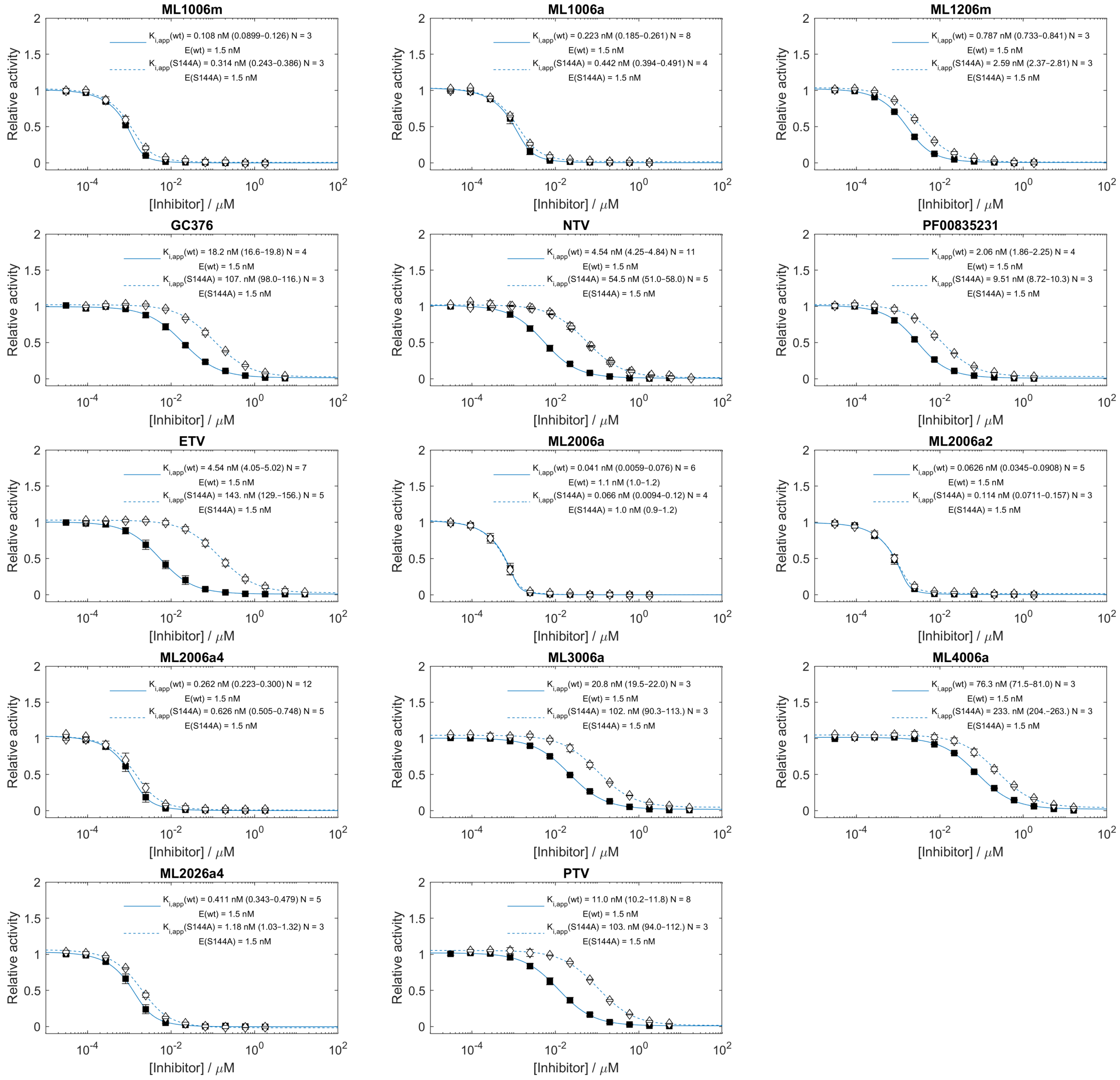


**Fig. S13. Equilibrium inhibitory constant ($K_i$) fits for $M^{pro}$-coil WT and S144A at 37 °C.**
The data of Fig. S9 was reanalyzed to obtain $K_{i,app}$ values by non-linear fitting of the data to the Morrison equation (Eq. 3). $CI_{95}$ of the fit is reported in parentheses. Error bars are SD.

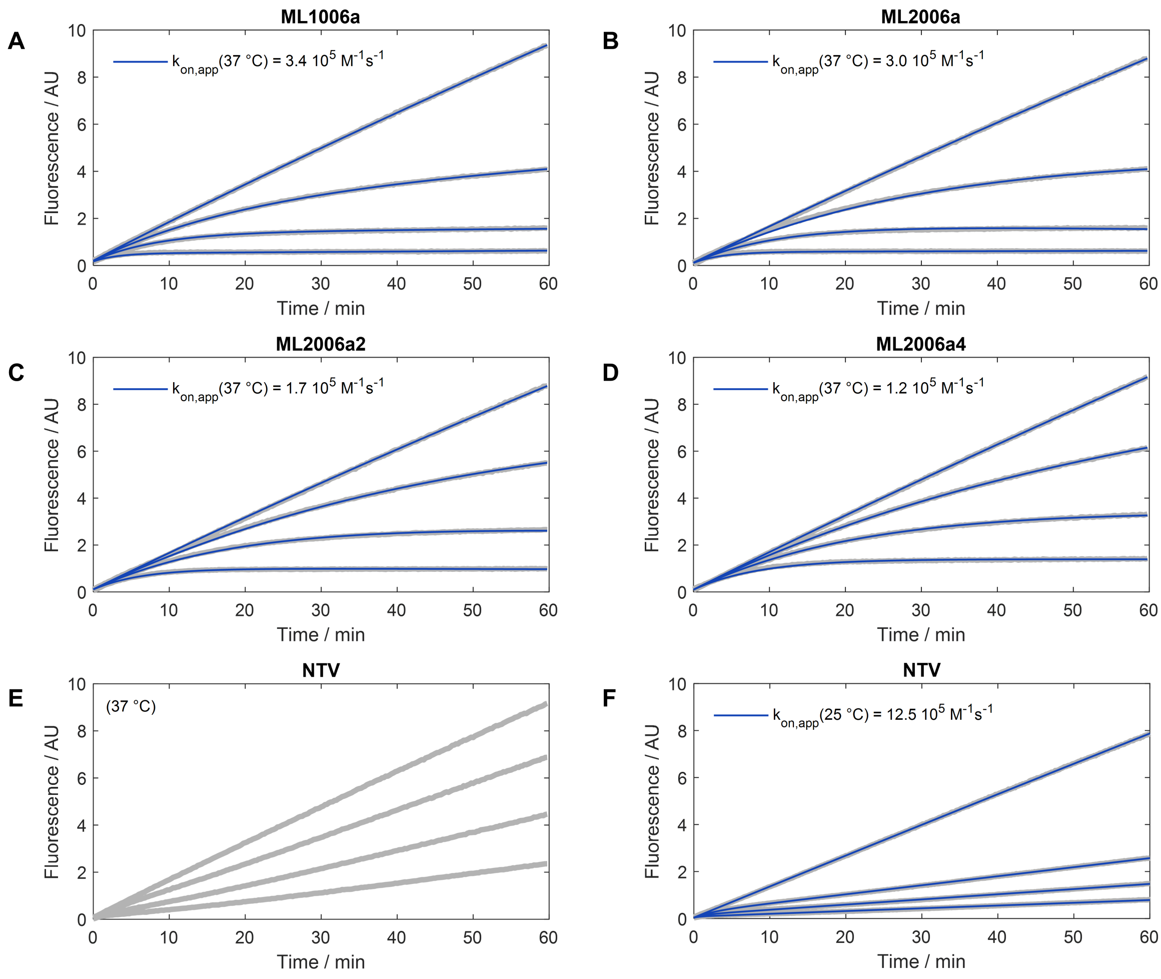


**Fig. S14. Inhibitor binding kinetics.**
Examples of time-resolved inhibition of $M^{pro}$-coil WT measured at 37 °C after the addition of (**A**) ML1006a, (**B**) ML2006a, (**C**) ML2006a2, (**D**) ML2006a4, and (**E**) NTV. From top to bottom the inhibitor concentrations are: 0, 1.67, 5.0, and 15 nM. Data are shown in grey and fits to Eq. 4 in blue. The NTV data does not show clear curvature during the measurement window. Thus, equilibrium is assumed to have settled prior to the measurement, and $k_{on}$ cannot be resolved at 37 °C. Therefore, (**F**) an experiment was performed at 25 °C for NTV with inhibitor concentrations of 0, 5, 10, and 20 nM from top to bottom. The lower temperature slowed the binding kinetics and allowed for fitting of Eq. 4 to the data. The $k_{on}$ measured at 25 °C was used as a lower limit for $k_{on}$ at 37 °C.

**A** Remaining Activity

| [Inhibitor] | Cat-B 15 µM | Cat-K 15 µM | Cat-L 15 µM | Cat-S 15 µM |
|---|---|---|---|---|
| **ML1001** | 0.28 | 0.07 | 0.60 | 0.00 |
| **ML1201** | 0.03 | 0.03 | 0.00 | 0.00 |
| **ML1001m** | 0.79 | 0.49 | 0.94 | 0.03 |
| **ML1004m** | 0.89 | 0.40 | 0.92 | 0.04 |
| **ML1005m** | 0.87 | 0.29 | 0.87 | 0.06 |
| **ML1006m** | 0.88 | 0.27 | 0.90 | 0.00 |
| **ML1206m** | 0.24 | 0.33 | 0.02 | -0.01 |
| **ML1006d** | 1.00 | 0.87 | 0.87 | 0.81 |
| **ML1006a** | 0.70 | 0.38 | 0.82 | 0.02 |
| **ML2006a1** | 0.89 | 0.52 | 0.88 | 0.01 |
| **ML2006a2** | 0.73 | 0.72 | 0.93 | 0.01 |
| **ML2006a3** | 0.95 | 0.87 | 0.98 | 0.00 |
| **ML2006a4** | 0.93 | 0.62 | 0.93 | 0.00 |
| **ML3006a1** | 0.93 | 0.94 | 0.74 | 0.05 |
| **ML4006a1** | 0.92 | — | 0.90 | 0.12 |
| **NTV** | 0.98 | 0.64 | 0.91 | 0.31 |
| **GC376** | 0.00 | -0.02 | 0.02 | 0.00 |

**B** Remaining Activity

| [Inhibitor] | Cat-B 1.5 µM | Cat-K 1.5 µM | Cat-L 1.5 µM | Cat-S 1.5 µM |
|---|---|---|---|---|
| **ML1001** | 0.82 | 0.28 | 1.00 | 0.00 |
| **ML1201** | 0.26 | 0.16 | 0.03 | 0.00 |
| **ML1001m** | 0.99 | 0.89 | 1.03 | 0.11 |
| **ML1004m** | 0.99 | 0.90 | 1.02 | 0.12 |
| **ML1005m** | 0.99 | 0.74 | 1.00 | 0.06 |
| **ML1006m** | 0.98 | 0.66 | 0.97 | 0.00 |
| **ML1206m** | 0.75 | 0.73 | 0.21 | 0.02 |
| **ML1006d** | 0.96 | 0.90 | 0.94 | 1.06 |
| **ML1006a** | 0.91 | 0.80 | 0.94 | 0.06 |
| **ML2006a1** | 0.94 | 0.83 | 0.98 | 0.16 |
| **ML2006a2** | 0.88 | 0.92 | 0.90 | 0.12 |
| **ML2006a3** | 0.94 | 0.95 | 0.96 | 0.19 |
| **ML2006a4** | 0.92 | 0.92 | 0.91 | 0.10 |
| **ML3006a1** | 0.92 | 0.96 | 0.86 | 0.32 |
| **ML4006a1** | 0.90 | 0.87 | 0.88 | 0.49 |
| **NTV** | 0.94 | 0.95 | 0.92 | 0.84 |
| **GC376** | 0.05 | 0.00 | 0.00 | 0.00 |

Cathepsin Activity: 1 – 0

**C** Standard Deviation Remaining Activity

| [Inhibitor] | Cat-B 15 µM | Cat-K 15 µM | Cat-L 15 µM | Cat-S 15 µM |
|---|---|---|---|---|
| **ML1001** | 0.00 | 0.00 | 0.01 | 0.01 |
| **ML1201** | 0.00 | 0.00 | 0.00 | 0.00 |
| **ML1001m** | 0.01 | 0.02 | 0.05 | 0.01 |
| **ML1004m** | 0.07 | 0.03 | 0.08 | 0.03 |
| **ML1005m** | 0.02 | 0.01 | 0.07 | 0.07 |
| **ML1006m** | 0.01 | 0.05 | 0.02 | * |
| **ML1206m** | 0.00 | 0.02 | 0.00 | 0.01 |
| **ML1006d** | 0.05 | 0.01 | 0.02 | 0.19 |
| **ML1006a** | 0.00 | 0.01 | 0.01 | 0.04 |
| **ML2006a1** | 0.01 | 0.03 | 0.12 | 0.01 |
| **ML2006a2** | 0.01 | 0.04 | 0.03 | 0.00 |
| **ML2006a3** | 0.00 | 0.05 | 0.03 | * |
| **ML2006a4** | 0.01 | 0.01 | 0.01 | 0.00 |
| **ML3006a1** | 0.02 | * | 0.23 | 0.01 |
| **ML4006a1** | 0.00 | ** | 0.02 | 0.00 |
| **NTV** | 0.01 | 0.03 | 0.17 | 0.04 |
| **GC376** | * | 0.02 | 0.03 | 0.00 |

**D** Standard Deviation Remaining Activity

| [Inhibitor] | Cat-B 1.5 µM | Cat-K 1.5 µM | Cat-L 1.5 µM | Cat-S 1.5 µM |
|---|---|---|---|---|
| **ML1001** | 0.03 | 0.01 | 0.03 | 0.02 |
| **ML1201** | 0.01 | 0.00 | 0.00 | 0.00 |
| **ML1001m** | 0.05 | 0.05 | 0.01 | 0.00 |
| **ML1004m** | 0.05 | 0.05 | 0.03 | 0.00 |
| **ML1005m** | 0.05 | 0.03 | 0.02 | 0.00 |
| **ML1006m** | 0.05 | 0.03 | 0.00 | * |
| **ML1206m** | 0.03 | 0.03 | 0.00 | 0.01 |
| **ML1006d** | 0.04 | 0.03 | 0.03 | 0.05 |
| **ML1006a** | 0.04 | 0.17 | 0.02 | 0.00 |
| **ML2006a1** | 0.04 | * | 0.00 | 0.05 |
| **ML2006a2** | 0.07 | 0.00 | 0.05 | * |
| **ML2006a3** | 0.03 | 0.03 | 0.01 | 0.01 |
| **ML2006a4** | 0.05 | 0.02 | 0.04 | 0.06 |
| **ML3006a1** | 0.04 | 0.01 | 0.01 | 0.03 |
| **ML4006a1** | 0.04 | 0.04 | 0.02 | 0.07 |
| **NTV** | 0.14 | 0.17 | 0.19 | 0.09 |
| **GC376** | 0.00 | 0.00 | 0.00 | 0.00 |

SD: 1 – 0

**Fig. S15. Cathepsin inhibition by M$^{pro}$ inhibitors.**
Remaining activity of human cathepsins B, K, L, and S when preincubated for 1 h with (**A**) 15 and (**B**) 1.5 µM of inhibitor, and (**C to D**) the standard deviation (*N = 2, ** $N$ = 1, ** $N$ = 0) related to A and B, respectively.

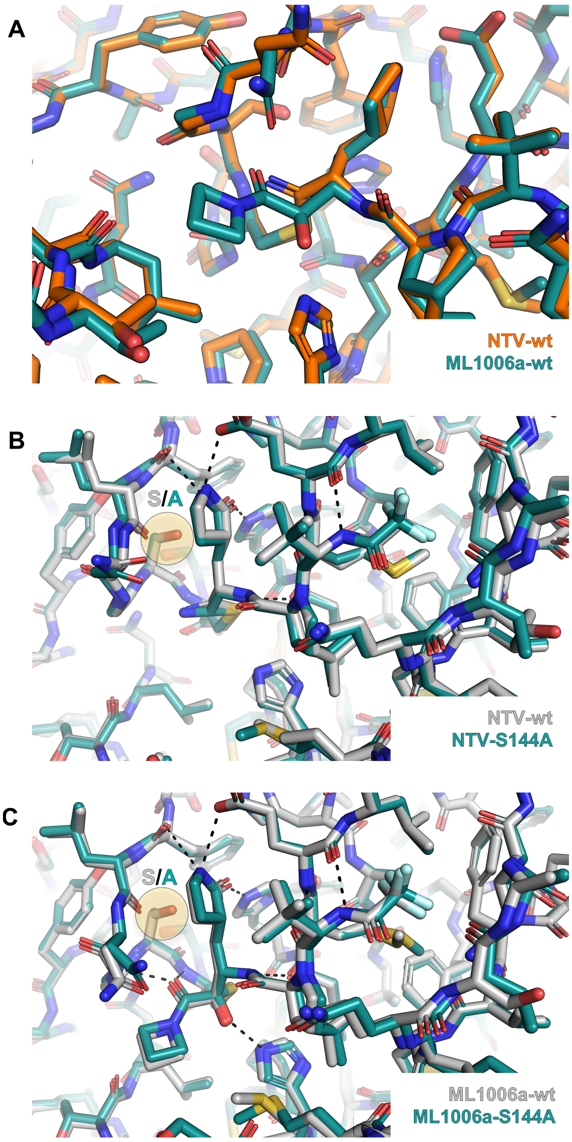


**Fig. S16. Structure comparison of M$^{pro}$ WT and S144A inhibited by ML1006a or NTV.**
(**A**) In the adduct state, the protein conformation of M$^{pro}$ WT is similar for the thioimidate (NTV) and the hemithioketal (here ML1006a) products. However, the thioimidate and hemithioketal interact differently with the catalytic dyad and oxyanion hole. Specifically, the ketoamide forms a sp$^3$ hybridized adduct with C145 while also forming H-bonds to H41 and the oxyanion hole, while the thioimidate adduct of the nitrile is sp$^2$ hybridized and not strongly H-bonded to H41 or the oxyanion hole. Overlays of SARS-CoV-2 M$^{pro}$ WT and S144A co-crystals complexed with (**B**) NTV and (**C**) ML1006a do not show any clear structural alterations due to the mutation. Since no clear conformational changes are seen in the static crystals, the effects of the S144A mutation may instead result from changes in protein dynamics. Specifically, the S144A mutation removes a H-bond donor that stabilizes the oxyanion loop conformation thereby potentially affecting the dynamics of C145 and the oxyanion hole and reactions and interactions with substrates or inhibitors. The coordinates of PDB 7VH8 (*47*) were used for representing the structure NTV bound in SARS-CoV-2 M$^{pro}$ WT in panel B.

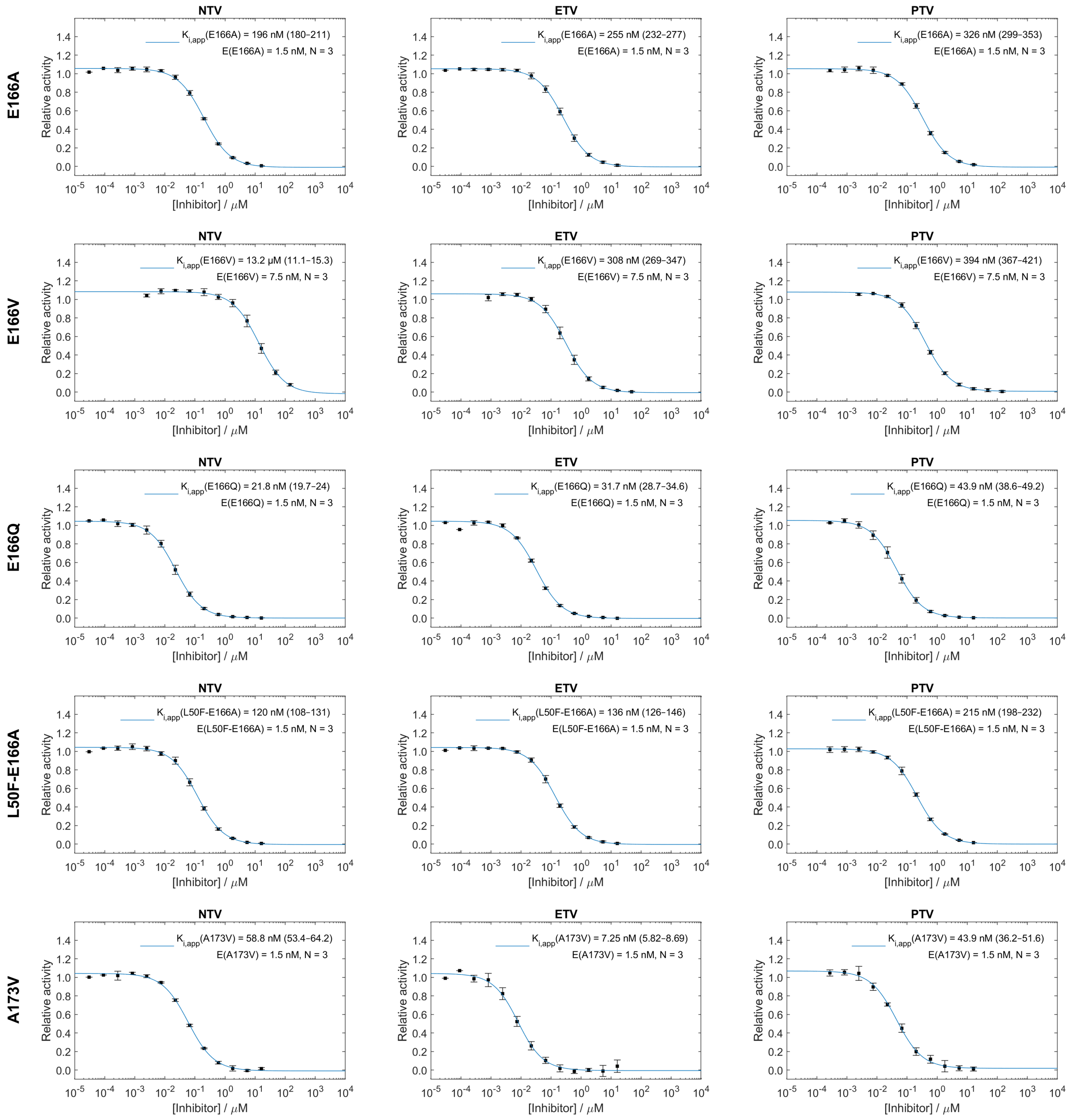


**Fig. S17. Equilibrium inhibitory constant ($K_i$) fits for $M^{pro}$-coil mutants at 37 °C.**
The Morrison equation (Eq. 3) was used to fit the *N* aggregated datasets and obtain $K_{i,app}$ values. $CI_{95}$ of the fit is reported in parentheses. Error bars are SD.

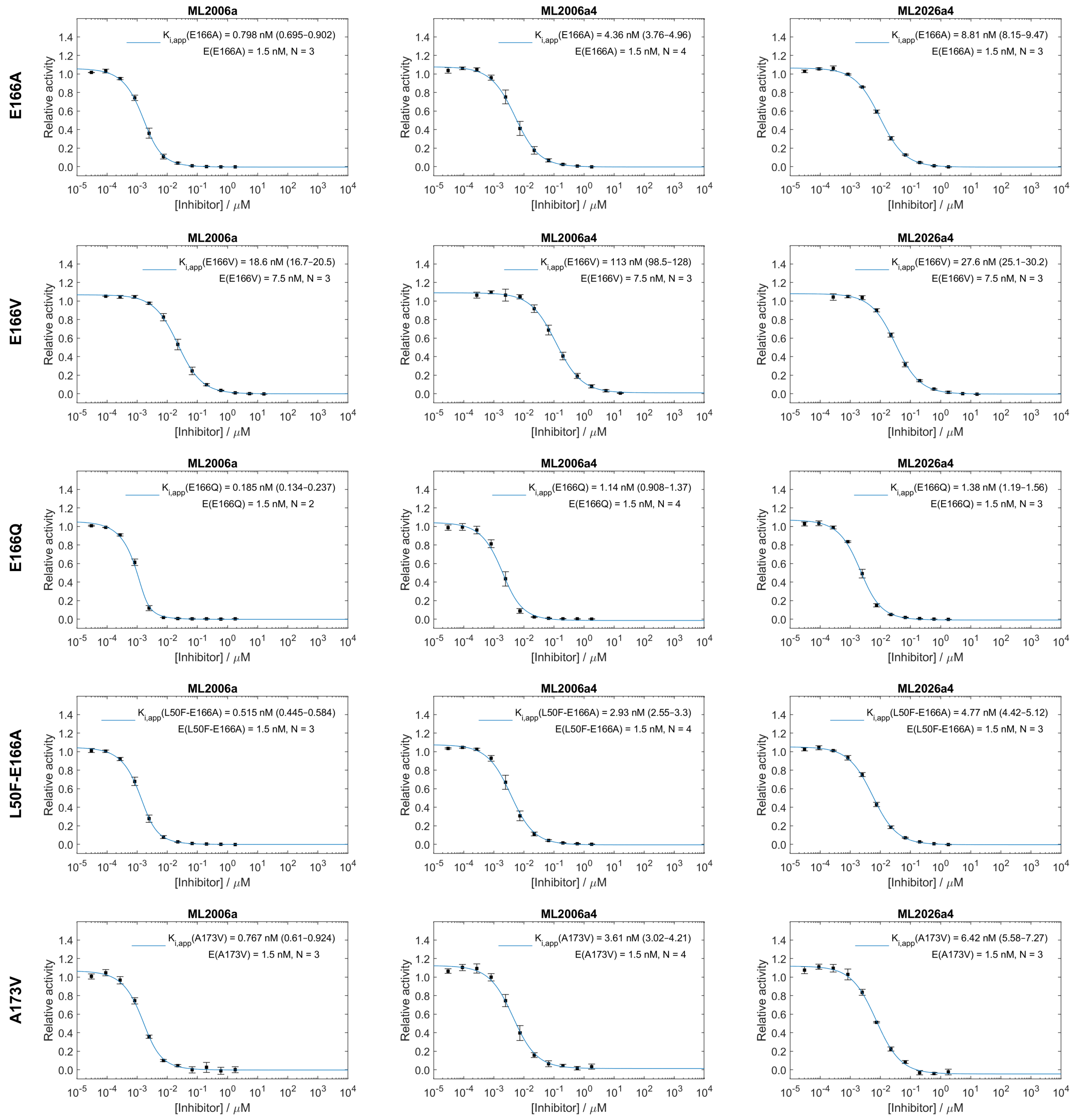


**Fig. S17 continued. Equilibrium inhibitory constant ($K_i$) fits for $M^{pro}$-coil mutants at 37 °C.**
The Morrison equation (Eq. 3) was used to fit the *N* aggregated datasets and obtain $K_{i,app}$ values. $CI_{95}$ of the fit is reported in parentheses. Error bars are SD.

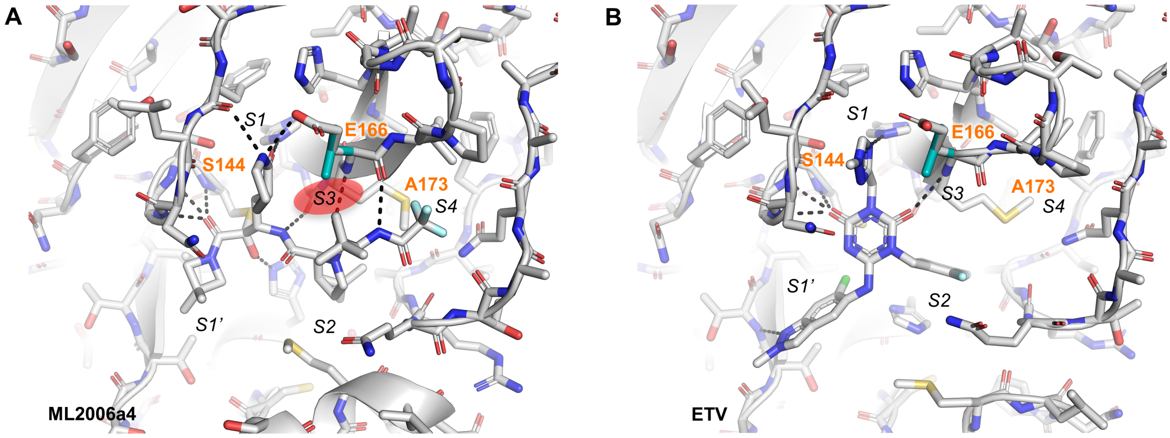


**Fig. S18. Models of M$^{pro}$ E166V in complex with ML2006a4 or ETV.**
Static models of the likely conformation of E166V (cyan) prepared by introducing the mutation in co-crystals of WT M$^{pro}$ (**A**) ML2006a4 or (**B**) ETV (PDB 8DZ0) (*48*). In ML2006a4-complexed M$^{pro}$ E166V model the predicted valine rotamer clashes (red oval) with the P3 tert-butyl of ML2006a4. NTV experiences a similar clash (not shown), while in contrast, ETV does not occupy the S3 site. The S144 and A173 residues are difficult to spot in these figures that focus on representing the E166V mutation, however, labels have been added to point to the position of S144 and A173.

**A**

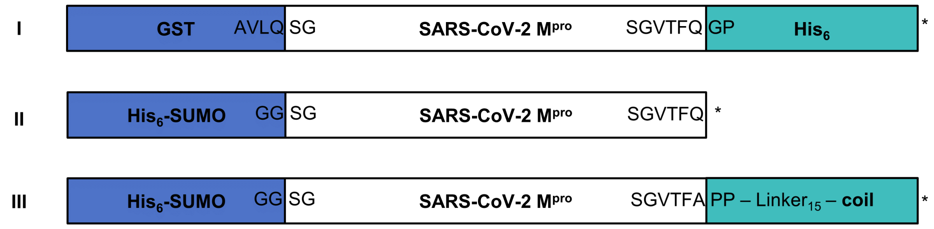


**B**

**Construct I: GST-M$^{pro}$-His$_6$ → M$^{pro}$-His$_6$**

MSPILGYWKIKGLVQPTRLLLEYLEEKYEEHLYERDEGDKWRNKKFELGLEFPNLPYYIDGDVKLTQSMAIIRYIADKHNMLGGCPK
ERAEISMLEGAVLDIRYGVSRIAYSKDFETLKVDFLSKLPEMLKMFEDRLCHKTYLNGDHVTHPDFMLYDALDVVLYMDPMCLDAFP
KLVCFKKRIEAIPQIDKYLKSSKYIAWPLQGWQATFGGGDHPPKGSITSAVLQSGFRKMAFPSGKVEGCMVQVTCGTTTLNGLWLDD
VVYCPRHVICTSEDMLNPNYEDLLIRKSNHNFLVQAGNVQLRVIGHSMQNCVLKLKVDTANPKTPKYKFVRIQPGQTFSVLACYNGS
PSGVYQCAMRPNFTIKGSFLNGSCGSVGFNIDYDCVSFCYMHHMELPTGVHAGTDLEGNFYGPFVDRQTAQAAGTDTTITVNVLAWL
YAAVINGDRWFLNRFTTTLNDFNLVAMKYNYEPLTQDHVDILGPLSAQTGIAVLDMCASLKELLQNGMNGRTILGSALLEDEFTPFD
VVRQCSGVTFQGPLGSHHHHHH*

**Construct II: His$_6$-SUMO-M$^{pro}$ → M$^{pro}$**

MGHHHHHHGSDSEVNQEAKPEVKPEVKPETHINLKVSDGSSEIFFKIKKTTPLRRLMEAFAKRQGKEMDSLRFLYDGIRIQADQTPE
DLDMEDNDIIEAHREQIGGSGFRKMAFPSGKVEGCMVQVTCGTTTLNGLWLDDVVYCPRHVICTSEDMLNPNYEDLLIRKSNHNFLV
QAGNVQLRVIGHSMQNCVLKLKVDTANPKTPKYKFVRIQPGQTFSVLACYNGSPSGVYQCAMRPNFTIKGSFLNGSCGSVGFNIDYD
CVSFCYMHHMELPTGVHAGTDLEGNFYGPFVDRQTAQAAGTDTTITVNVLAWLYAAVINGDRWFLNRFTTTLNDFNLVAMKYNYEPL
TQDHVDILGPLSAQTGIAVLDMCASLKELLQNGMNGRTILGSALLEDEFTPFDVVRQCSGVTFQ*

**Construct III: His$_6$-SUMO-M$^{pro}$-Linker$_{15}$-Coil → M$^{pro}$-Linker$_{15}$-Coil**

MGHHHHHHGSDSEVNQEAKPEVKPEVKPETHINLKVSDGSSEIFFKIKKTTPLRRLMEAFAKRQGKEMDSLRFLYDGIRIQADQTPE
DLDMEDNDIIEAHREQIGGSGFRKMAFPSGKVEGCMVQVTCGTTTLNGLWLDDVVYCPRHVICTSEDMLNPNYEDLLIRKSNHNFLV
QAGNVQLRVIGHSMQNCVLKLKVDTANPKTPKYKFVRIQPGQTFSVLACYNGSPSGVYQCAMRPNFTIKGSFLNGSCGSVGFNIDYD
CVSFCYMHHMELPTGVHAGTDLEGNFYGPFVDRQTAQAAGTDTTITVNVLAWLYAAVINGDRWFLNRFTTTLNDFNLVAMKYNYEPL
TQDHVDILGPLSAQTGIAVLDMCASLKELLQNGMNGRTILGSALLEDEFTPFDVVRQCSGVTF*APP*GGSGGSGGGSGGSGGEIAALK
QEIAALKKENAALKWEIAALKQG*

**Fig. S19. Constructs used for M$^{pro}$ expression and purification.**
(**A**) Schematic drawing of the M$^{pro}$ constructs expressed in *E. Coli*. (**B**) Amino acid sequences of the expressed proteins (here in WT form) with domains annotated by color. The final construct obtained after the full purification procedure and used in the biochemical assays are underlined. Construct III contains the mutated amino acids APP (italics) at the C-terminus of M$^{pro}$ to prevent autocatalytic processing.

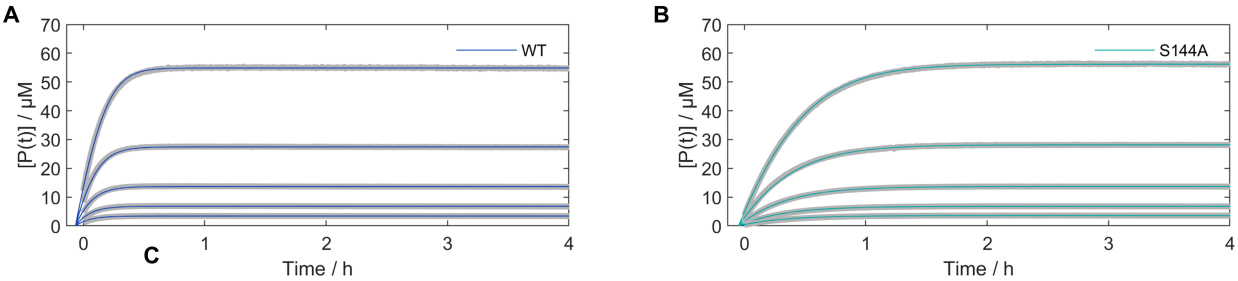


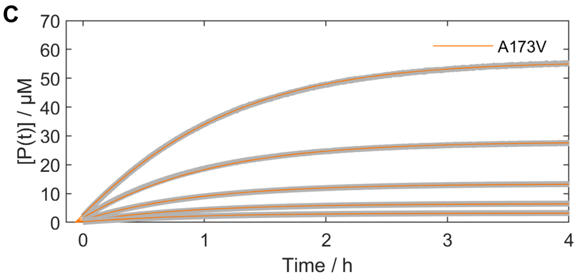


**D**

| | *N* | $K_m$ [μM] | $k_{cat}$ [s$^{-1}$] | $k_{cat}/K_m$ [10$^4$ s$^{-1}$M$^{-1}$] | Relative catalytic efficiency |
|---|---|---|---|---|---|
| **WT** | 3 | 44 ± 7 | 2.0 ± 0.5 | 4.5 ± 0.6 | 1 |
| **S144A** | 3 | 101 ± 25 | 1.4 ± 0.5 | 1.3 ± 0.2 | 0.30 |
| **A173V** | 2 | 139 ± 31 | 0.9 ± 0.2 | 0.65 ± 0.01 | 0.14 |

**Fig. S20. M$^{pro}$ enzyme characterization.**
Michaelis-Menten parameters of M$^{pro}$ proteolysis of Covidyte TF670 were determined for M$^{pro}$-coil (**A**) WT, (**B**) S144A, and (**C**) A173V via global non-linear fitting of Eq. 1 to the production formation curves. The grey curves are representative data at increasing substrate concentrations between ~3 and 55 µM, and the colored lines are the associated fits. (**D**) Average data of independent experiments. The E166X mutants gave rise to complicated kinetics of product formation that could not be described by Eq. 1. Further, biochemical investigations of these mutants are needed to better understand the nature of this non-traditional enzyme behavior and build a suitable model. Thus, no Michaelis-Menten parameters are reported for the E166X mutants.

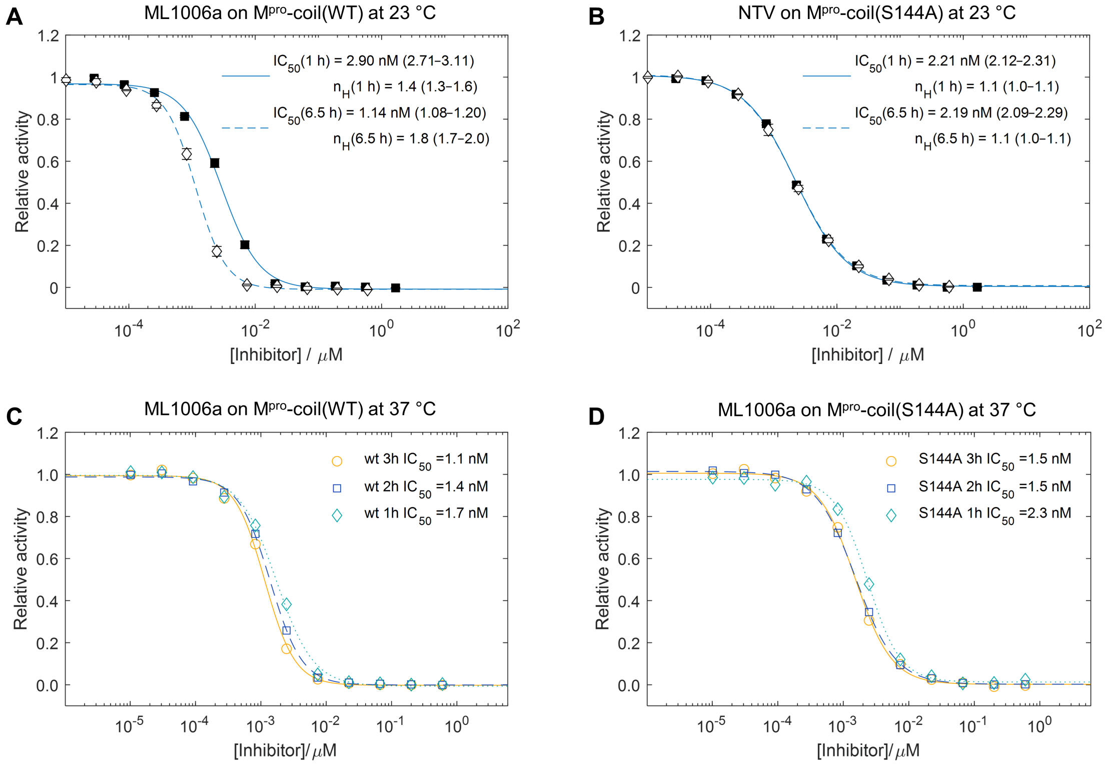


**Fig. S21. Extended equilibration times are required for the slow binding ketoamide-based inhibitors.** Ketoamides showed slow reversible binding kinetics leading to incomplete equilibration with $M^{pro}$-coil at RT (~ 23 °C) after 1 h. In contrast, NTV reaches equilibrium under these conditions. $M^{pro}$-coil WT inhibition curves after preincubation with (**A**) ML1006a and (**B**) NTV at RT for 1 and 6.5 h preincubation, respectively. A clear decrease in the $IC_{50}$ value is seen for ML1006a but not NTV. Thus, $IC_{50}$ values reported for ML1006a and related slow binding ketoamides in Fig. 2 underestimate the equilibrium $M^{pro}$ binding affinity. $IC_{50}$ values and Hill coefficients, $n_H$, were extracted by aggregating all datasets (*N = 3*) and performing non-linear regression. $CI_{95}$ of the fit is reported in parentheses. Error bars are SD. Equilibration was also assayed at elevated temperature by determining $M^{pro}$-coil (**C**) WT and (**D**) S144A inhibition curves after 1, 2, or 3 h preincubation with ML1006a at 37 °C. Within the precision of the experiment, no large change is seen in the $IC_{50}$ value after 2 h of incubation. Thus, a preincubation step of 3 h was used to determine the equilibrium $K_i$ values reported in this study. No decrease in $M^{pro}$ activity was observed during the extended incubation times at either 25 or 37 °C.

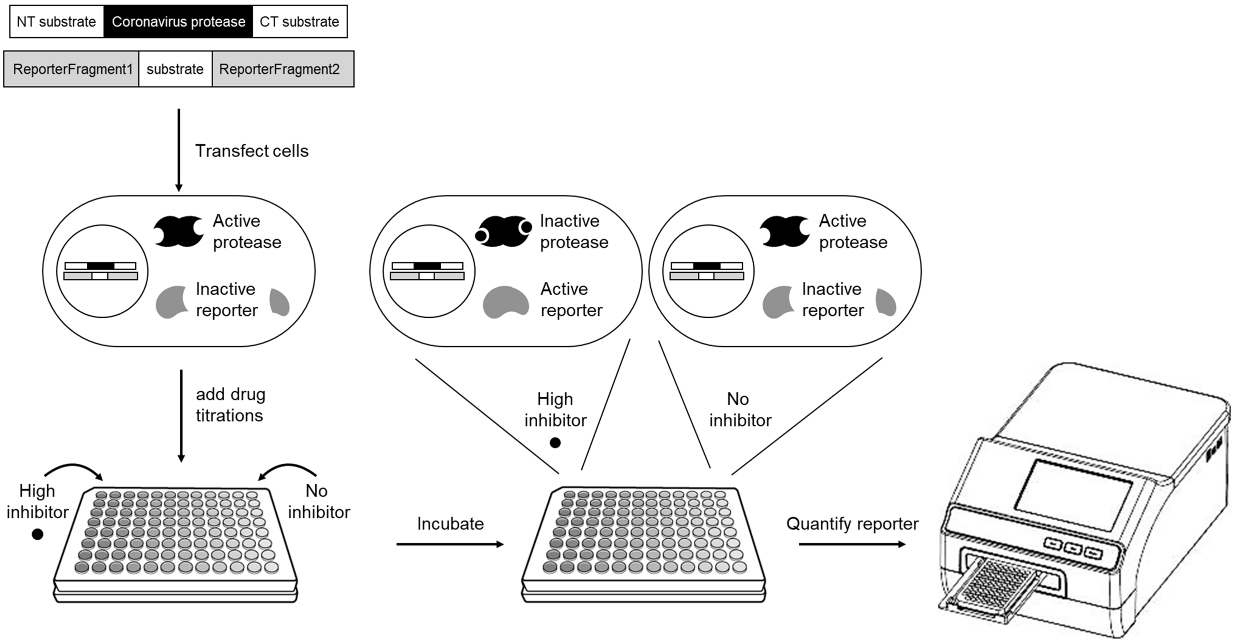


**Fig. S22. Schematic of bioluminescence-based assay for intracellular detection of M$^{pro}$ inhibition.** Huh7 cells are transiently transfected with a plasmid coexpressing a M$^{pro}$ precursor and a NanoLuc bioluminescent reporter with an internal M$^{pro}$ substrate site. If expressed in the absence of drug inhibitors, M$^{pro}$ autocatalytically matures and then cleaves the NanoLuc reporter to inactive it, suppressing bioluminescence. In the presence of drug inhibitors, active NanoLuc accumulates, producing high bioluminescence.

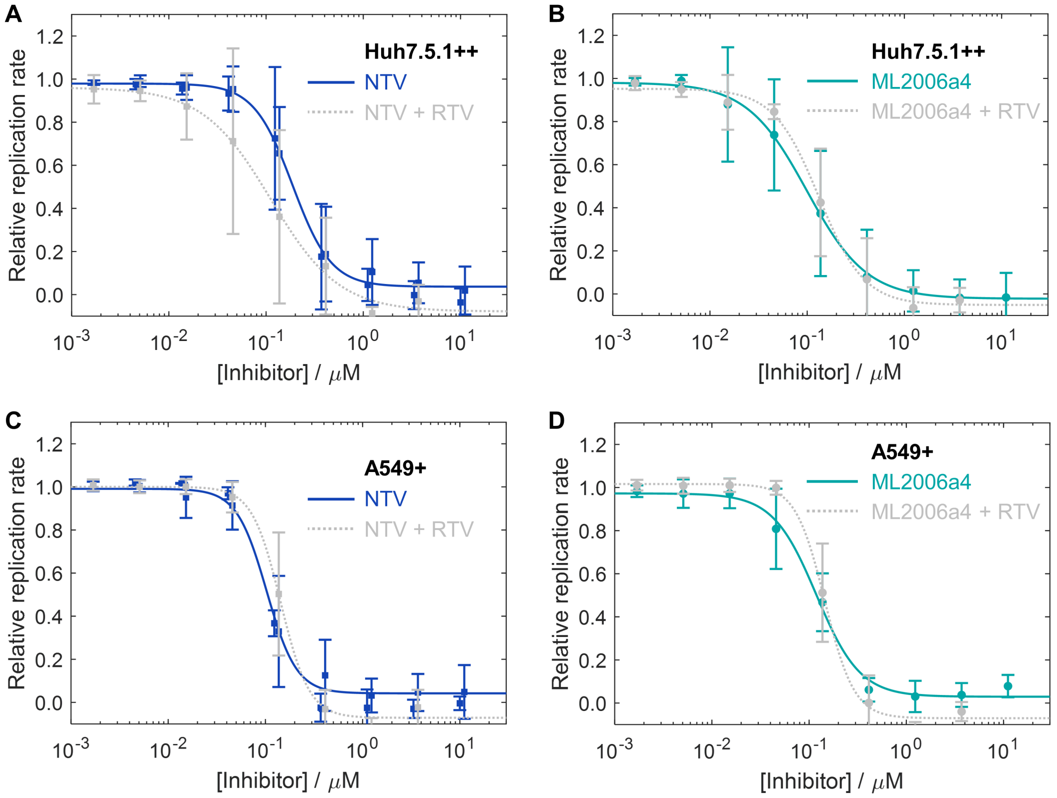


**Fig. S23. The antiviral potencies of ML2006a4 and NTV are RTV independent.**
Inhibition of viral replication in Huh7.5.1++ cells by (**A**) NTV or (**B**) ML2006a4 in the presence or absence of 1 µM RTV. Inhibition of viral replication in A549+ cells by (**C**) NTV or (**D**) ML2006a4 in the presence or absence of 1 µM RTV. The data recorded without RTV is replotted from Fig. S5 and S6. *N,n* = 2,6 for all datasets recorded in the presence of RTV. Error bars are SD. Within the margin of error, we do not observe a systematic change in the antiviral potency of ML2006a4 or NTV in the presence of 1 µM RTV. This is consistent with the expectation that RTV does not have an antiviral effect (Fig. S2), and A549+, Huh7.5.1++, or Calu-3 cells are not known to overexpress P-gp efflux transporters (*49–51*), that can otherwise be inhibited by RTV (*52*). Thus, throughout this paper we have consistently referred to $EC_{50}$ and $EC_{90}$ values determined in the absence of RTV since these have more independent data series and thus presumably better accuracy. Similarly, when plotting the $EC_{50,p}$ and $EC_{90,p}$ values together with the pharmacokinetic data, the $EC_{50}$ values obtained in Huh7.5.1++ in the absence of RTV were used.

### Supplementary Tables

**Table S1. In vitro ADME.**

| Compound | $P_{app}$ A-B[a] [$10^{-6}$ cm/s] | $P_{app}$ ratio[a] B-A/A-B | Solubility[b] [µM] | $f_{u,p}$[c] % | | $t_{1/2,p}$[d] [min] | | $CL_{int,app}$[e] [µL min$^{-1}$ mg$^{-1}$] | | | |
|---|---|---|---|---|---|---|---|---|---|---|---|
| | | | *PBS* | *mouse* | *human* | *mouse* | *human* | *mouse* | | *human* | |
| | | | | | | | | - | + RTV | - | + RTV |
| **ML1004m** | 0.27 | 1.8 | 83 | - | - | 180 | - | - | - | <5 | <5 |
| **ML1005m** | 0.34 | 1.6 | 76 | - | - | 217 | - | <5 | <5 | <5 | <5 |
| **ML1006m** | 0.22 | 4.3 | 75 | 53 | - | 209 | >450 | <5 | <5 | <5 | <5 |
| **ML1006a** | 0.08 | 105 | 74 | 33 | 46 | 275 | >450 | 27 | <5 | 13 | <5 |
| **ML2006a** | 0.22 | 93 | 63 | 24[e] | 39 | 81 | 188 | 63 | <5 | 32 | <5 |
| **ML2006a2** | 0.90 | 39 | 61 | - | 35 | <5 | 135 | 97 | <5 | 60 | <5 |
| **ML2006a4** | 1.0 | 55 | 85 | 15 | 19 | 45 | 189 | 110[e] | <5 | 77[e] | <5 |
| **ML3006a** | 0.65 | 38 | 53 | - | 32 | 104 | >450 | 48 | <5 | 22 | <5 |
| **ML4006a** | 1.7 | 19 | 47 | - | 29 | 45 | >450 | 100 | <5 | 48 | <5 |
| **NTV** | 0.51 | 25 | 50 | 24 | 29 | 150 | >450 | 108[e] | <5 | 18[e] | <5 |
| **PTV** | 2.9 | 27 | 23 | 1.3 | 1.4 | 330 | 148 | 100 | 5.4 | 152 | 41 |
| **GC376** | 0.37 | 35 | 71 | - | - | 232 | 95 | - | - | - | - |
| **BPV** | 0.92 | 18 | 73 | - | - | 28 | >450 | - | - | - | - |

[a]Apparent permeability, $P_{app}$, in Caco-2 cell layers in apical-to-basal (A-B) and basal-to-apical (B-A) directions. [b]Kinetic solubility at 100 µM in PBS with 1% DMSO. [c]Fraction of unbound compound in either 100% C57BL/6 mouse plasma or 100% human plasma at 37 °C. [d]Stability at 37 °C in 90% CD-1 mouse or human plasma. [e]Apparent intrinsic clearance calculated from the half-life in 0.5 mg/mL CD-1 mouse or human liver microsomes at 37 °C without or with 1 µM RTV added as a CYP3A inhibitor. [e]N = 2, experiments performed on different dates.

**Table S2. Inhibitor cytotoxicity.**[a]

| | **% Cell Viability** | | | |
|---|---|---|---|---|
| | A549-ACE2 | | Huh7 | |
| Compound Conc [μM] | 10 | 100 | 10 | 100 |
| **ML1000** | 99 | 100 | 105 | 104 |
| **ML1100** | 118 | 99 | 111 | 104 |
| **ML1001** | 98 | 96 | 101 | 102 |
| **ML1001m** | 103 | 101 | 102 | 102 |
| **ML1002** | 97 | 89 | 109 | 99 |
| **ML1002m** | 98 | 98 | 107 | 108 |
| **ML1003** | 100 | 98 | 104 | 89 |
| **ML1004m** | 104 | 99 | 108 | 100 |
| **ML1005m** | 102 | 100 | 106 | 96 |
| **ML1006m** | 100 | 106 ± 10 (*N* = 2) | 106 | 85 ± 17 (*N* = 3) |
| **ML1006d** | 113 | 121 | 119 | 110 |
| **ML1006a** | 104 | 107 ± 12 (*N* = 2) | 107 ± 11 (*N* = 2) | 83 ± 9 (*N* = 3) |
| **ML2006a** | 94 ± 2 (*N* = 2) | 87 ± 0 (*N* = 2) | 96 ± 1 (*N* = 2) | 93 ± 0 (*N* = 2) |
| **ML2006a2** | 96 ± 0 (*N* = 2) | 91 ± 1 (*N* = 2) | 97 ± 3 (*N* = 2) | 92 ± 1 (*N* = 2) |
| **ML2006a4** | 93 ± 3 (*N* = 2) | 87 ± 1 (*N* = 2) | 89 ± 12 (*N* = 2) | 87 ± 5 (*N* = 2) |
| **ML3006a** | 93 | 87 | 89 | 86 |
| **ML4006a** | 99 | 91 | 90 | 78 |
| **ML1201** | 99 | 103 | 108 | 107 |
| **ML1206m** | - | 114 | 118 | 107 |
| **GC376** | - | 98 | 107 | 97 |
| **NTV** | 103 | 108 ± 12 (*N* = 2) | 109 | 103 ± 12 (*N* = 3) |
| **BPV** | - | 79 | 106 | 88 |

[a]All viability measurements were performed in technical triplicates ($n = 3$) and the mean reported. For some compounds, biological replicates (*N* as noted in table) were performed, and the standard deviation reported.

**Table S3. Bidirectional binding rates at 37 °C.**

| | $K_i$ $M^{pro}$-coil WT [nM][a] | $k_{on}$[b] $M^{pro}$-coil WT [$10^5$ $M^{-1}$ $s^{-1}$] | $k_{off,calc}$[c] $M^{pro}$-coil WT [$10^{-5}$ $s^{-1}$] | $t_{1/2,calc}$[d] $M^{pro}$-coil WT [h] |
|---|---|---|---|---|
| **ML1006a** | 0.22 (0.19–0.26) | 3.9 | 8.7 | 2.2 |
| **ML2006a** | 0.04 (0.01–0.08) | 3.4 | 1.4 | 14 |
| **ML2006a2** | 0.06 (0.03–0.09) | 2.0 | 1.2 | 16 |
| **ML2006a4** | 0.26 (0.22–0.30) | 1.4 | 3.5 | 5.4 |
| **NTV** | 4.5 (4.3–4.8) | >13[e] | >610[e] | <0.032[e] |

[a]Equilibrium inhibition constants obtained from data in Fig. S13. $CI_{95}$ is reported in parentheses.
[b]Association rate constants, $k_{on}$, obtained from the fits in Fig. S14 and corrected for substrate competition.
[c]While $k_{on}$ can be accurately determined from the inhibitor binding progression curves, quantification of $k_{off}$ in Eq. 4 is associated with large uncertainty. Thus, we calculated $k_{off,calc} = K_i \cdot k_{on}$ for a two-state reversible binding model.
[d]Calculated dissociation half-life, $t_{1/2,calc} = \ln(2)/k_{off,calc}$.
[e]The binding kinetics of NTV were too rapid for quantification at 37 °C with the available instrumentation. Thus, $k_{on,fit}$ measured at 25 °C was used as a lower limit estimate. This value was propagated in the calculation of $k_{off,calc}$ and $t_{1/2,calc}$.

**Table S4. Sensitivity of inhibitor affinity to the $M^{pro}$ S144A mutation**

| | $K_i$ ($M^{pro}$-coil, 37 °C) | | $K_i$ ratio |
|---|---|---|---|
| | WT [nM][a] | S144A [nM][a] | S144A / WT |
| **ML1006m** | 0.11 (0.09–0.13) | 0.31 (0.24–0.39) | 2.9 |
| **ML1206m** | 0.79 (0.73–0.84) | 2.6 (2.4–2.8) | 3.3 |
| **ML1006a** | 0.22 (0.19–0.26) | 0.44 (0.39–0.49) | 2.0 |
| **ML2006a** | 0.04 (0.01–0.08) | 0.07 (0.01–0.12) | 1.6 |
| **ML2006a2** | 0.06 (0.03–0.09) | 0.11 (0.07–0.16) | 1.8 |
| **ML2006a4** | 0.26 (0.22–0.30) | 0.63 (0.51–0.75) | 2.4 |
| **ML2026a4** | 0.41 (0.34–0.48) | 1.2 (1.0–1.3) | 2.9 |
| **ML3006a** | 21 (19–22) | 102 (90–113) | 4.9 |
| **ML4006a** | 76 (72–81) | 233 (204–263) | 3.1 |
| **GC376** | 18 (17–20) | 107 (98–116) | 5.9 |
| **PF00835231** | 2.1 (1.9–2.3) | 9.5 (8.7–10.3) | 4.6 |
| **NTV** | 4.5 (4.3–4.8) | 55 (51–58) | 12 |
| **ETV** | 4.5 (4.1–5.0) | 143 (129–156) | 31 |
| **PTV** | 11 (10–12) | 103 (94–112) | 9.3 |

[a]Equilibrium inhibition constants obtained from data in Fig. S13. $CI_{95}$ is reported in parentheses.

### Table S5. Crystallization conditions.

| PDB ID | Inhibitor | Protein | Precipitant | (Protein:Precipitant) [uL : uL] | Cryoprotectant |
|---|---|---|---|---|---|
| **7SET** | ML1000 0.5 mM | 9 mg/mL $M^{pro}$ WT in 50 mM Tris pH 7.3, 2 mM DTT, 5% DMSO | Molecular Dimensions Proplex HT-96 well D9: 0.1 M MES pH 6.5, 15% w/v PEG 6000, 5% v/v 2-methyl-2,4-petanediol | 0.25:0.25 | 25% glycerol |
| **7SF1** | ML1001 1.5 mM | 5 mg/mL $M^{pro}$ WT in 20 mM Tris pH 7.3, 2 mM DTT, 5% DMSO | Hampton Research PEGRx HT well F5: 0.1 M HEPES pH 7.5, 0.2 M L-Proline, 24% w/v PEG 1500 | 0.16:0.16 | 30% glycerol |
| **7SF3** | ML1006m 1 mM | 7 mg/mL $M^{pro}$ WT in 20 mM Tris pH 7.3, 2 mM DTT, 3% DMSO | Molecular Dimension JCSG-*plus* HT-96 well B4: 0.1 M HEPES pH 7.5, 10% w/v PEG8000 | 0.18:0.18 | 20% glycerol |
| **7U9A** | ML1006a 0.9 mM | 5.2 mg/mL $M^{pro}$ WT in 20 mM Tris pH 7.3, 3 mM TCEP, 3% DMSO | 0.1 M MES pH 6.5, 8% w/v PEG20000 | 0.20:0.20 | 25% PEG 200 |
| **7UUG** | ML1006a 0.9 mM | 5.1 mg/mL $M^{pro}$ S144A in 20 mM Tris pH 7.3, 3 mM TCEP, 3% DMSO | 0.1 M MES pH 6.5, 10% w/v PEG20000 | 0.19:0.19 | 25% PEG 200 |
| **7UUP** | ML1006N 0.9 mM | 5.1 mg/mL $M^{pro}$ S144A in 20 mM Tris pH 7.3, 3 mM TCEP, 3% DMSO | 0.1 M MES pH 6.5, 10% w/v PEG20000 | 0.19:0.19 | 25% PEG 200 |
| **8EZV** | ML2006a 0.9 mM | 5.3 mg/mL $M^{pro}$ WT in 20 mM Tris pH 7.3, 3 mM TCEP, 3% DMSO | Hampton Research PEGRx HT well H2: 20% v/v 2-Propanol, 0.1 M Tris pH 8.0, 5% w/v PEG 8000 | 0.23:0.23 | 25% PEG 200 |
| **8EZZ** | ML2006a2 0.9 mM | 5.3 mg/mL $M^{pro}$ WT in 20 mM Tris pH 7.3, 3 mM TCEP, 3% DMSO | Hampton Research PEGRx HT well D6: 0.1 M HEPES pH 7.5 4% w/v PEG 8000 | 0.23:0.23 | 25% PEG 200 |
| **8F02** | ML2006a4 0.9 mM | 5.3 mg/mL $M^{pro}$ WT in 20 mM Tris pH 7.3, 3 mM TCEP, 3% DMSO | Hampton Research PEGRx HT well C12: 0.1 M BICINE pH 8.5, 8% w/v mPEG 5000 | 0.23:0.23 | 25% PEG 200 |
| **8F2C** | ML3006a 0.9 mM | 5.3 mg/mL $M^{pro}$ WT in 20 mM Tris pH 7.3, 3 mM TCEP, 3% DMSO | 0.1 M MES pH 6.5, 4% PEG 35000 | 0.23:0.23 | 25% PEG 200 |
| **8F2D** | ML4006a 0.9 mM | 5.3 mg/mL $M^{pro}$ WT in 20 mM Tris pH 7.3, 3 mM TCEP, 3% DMSO | 0.1 M HEPES pH 7.5, 6% PEG 20000 | 0.23:0.23 | 25% PEG 200 |

### Table S6. Diffraction data and refinement statistics (1/2)

| $M^{pro}$ variant | WT | WT | WT | WT | S144A | S144A |
|---|---|---|---|---|---|---|
| **Inhibitor** | ML1000 | ML1001 | ML1006m | ML1006a | ML1006a | NTV |
| **PDB ID** | **7SET** | **7SF1** | **7SF3** | **7U92** | **7UUG** | **7UUP** |
| **Data Collection** | | | | | | |
| Beamline | SSRL BL12-2 | SSRL BL12-1 | SSRL BL12-2 | SSRL BL12-2 | SSRL BL12-2 | SSRL BL12-2 |
| Wavelength (Å) | 0.97946 | 0.97946 | 0.97946 | 0.97946 | 0.97946 | 0.97946 |
| Space group | $P2_12_12$ | C121 | $P2_12_12$ | $P2_12_12$ | $P2_12_12$ | $P2_12_12$ |
| Cell dimensions | | | | | | |
| *a, b, c* (Å) | 45.40, 63.26, 104.86 | 114.55, 52.92, 45.19 | 45.68, 64.44, 105.45 | 45.53, 64.26, 104.82 | 45.46, 63.61, 104.75 | 45.45, 63.96, 105,16 |
| *α, β, γ* (°) | 90, 90, 90 | 90, 103.32, 90 | 90, 90, 90 | 90, 90, 90 | 90, 90, 90 | 90, 90, 90 |
| Wilson B factor[a] | 23.8 | 32.5 | 23.6 | 24.5 | 44.5 | 33.5 |
| Matthews coef. ($Å^3$/Da)[b] | 2.23 | 1.97 | 2.25 | 2.27 | 2.24 | 2.26 |
| Solvent content (%) | 44.74 | 37.57 | 45.47 | 45.79 | 45.09 | 45.59 |
| Resolution (Å)[c] | 36.88 (1.70) | 39.19 (1.85) | 37.27 (1.75) | 37.15 (1.80) | 36.99 (2.00) | 37.05 (2.00) |
| No. of reflections/unique | 98934 / 32102 | 86585 / 22318 | 117206 / 31676 | 127506 / 29103 | 142206 / 21242 | 77254 / 21186 |
| $R_{merge}$[d] | 0.064 (0.532) | 0.083 (0.611) | 0.074 (0.836) | 0.086 (0.866) | 0.061 (1.269) | 0.095 (1.223) |
| $I/\sigma(I)$[e] | 9.2 (1.9) | 6.9 (1.8) | 9.6 (1.7) | 9.6 (1.9) | 13.2 (1.5) | 22.7 (1.2) |
| $CC_{1/2}$ | 0.997 (0.778) | 0.995 (0.768) | 0.998 (0.655) | 0.997 (0.738) | 0.999 (0.585) | 0.996 (0.419) |
| Completeness (%)[f] | 95.0 (97.0) | 98.8 (99.2) | 98.6 (98.9) | 99.5 (99.2) | 99.9 (100) | 99.2 (99.3) |
| Redundancy[g] | 3.1 (3.0) | 3.9 (3.9) | 3.7 (3.7) | 4.4 (4.1) | 6.7 (6.9) | 3.6 (3.7) |
| **Refinement** | | | | | | |
| Resolution (Å) | 36.91–1.70 | 39.22–1.85 | 37.30–1.75 | 35.04–1.80 | 34.94–2.00 | 35.08–2.00 |
| No. reflections/test set | 30479 / 1612 | 21183 / 1122 | 30068 / 1579 | 27596 / 1484 | 20152 / 1066 | 20110 / 1056 |
| $R_{work}$ / $R_{free}$[h] | 0.167 / 0.206 | 0.165 / 0.208 | 0.169 / 0.195 | 0.172 / 0.207 | 0.182 / 0.240 | 0.191 / 0.237 |
| $F_{obs}$-$F_{calc}$correlation[i] | 0.974 | 0.976 | 0.969 | 0.968 | 0.974 | 0.966 |
| No. atoms | | | | | | |
| Protein | 2407 | 2403 | 2403 | 2411 | 2393 | 2370 |
| Inhibitor | 39 | 39 | 39 | 41 | 41 | 35 |
| Inhibitor RSCC[j] | 0.96 | 0.92 | 0.93 | 0.92 | 0.95 | 0.94 |
| Other ligand/ion | 1 (Cl) | - | 1 (Cl) | 1 (Cl) | 1 (Cl) | 1 (Cl) |
| Water | 278 | 165 | 238 | 202 | 68 | 87 |
| Median *B*-factors | | | | | | |
| Protein | 26 | 37 | 24 | 27 | 49 | 39 |
| Inhibitor | 23 | 42 | 22 | 25 | 48 | 39 |
| Ligand/Ion | 42 | - | 54 | 46 | 60 | 52 |
| R.m.s. deviations | | | | | | |
| Bond lengths (Å) | 0.010 | 0.009 | 0.012 | 0.010 | 0.008 | 0.007 |
| Bond angles (°) | 1.604 | 1.543 | 1.691 | 1.690 | 1.591 | 1.488 |
| Ramachandran stats.[k] | | | | | | |
| Favored+Allowed (%) | 99.3 | 99.7 | 99.7 | 100 | 99.7 | 99.7 |
| Outliers (%) | 0.7 | 0.3 | 0.3 | 0 | 0.3 | 0.3 |

[a]Overall B-factor value, an approximation to the fall-off of atomic scattering with resolution.
[b]Ratio of the volume of the asymmetric unit to the molecular weight of all protein molecules in the asymmetric unit.
[c]Value in parenthesis is for the highest-resolution shell: 1.70-1.79 Å in 7SET, 1.85-1.95 Å in 7SF1, 1.75-1.84 Å in 7SF3, 1.80-1.90 Å in 7U9A, 2.00-2.11 Å in 7UUG, and 2.00-2.11 Å in 7UUP. Values in parentheses in subsequent rows relates to this highest resolution shell.
[d]Reliability factor for symmetry-related reflections calculated as: $R_{merge} = \Sigma_{hkl}\Sigma j=1 \text{ to } N \mid I_{hkl} - I_{hkl}(j) \mid / \Sigma_{hkl}\Sigma j=1 \text{ to } N\ I_{hkl}(j)$, where N is the redundancy of the data.
[e]Ratio of mean intensity to the mean standard deviation of the intensity over the entire resolution range.
[f]Fraction of measured reflections to possible observations at the resolution range.
[g]Number of measurements of individual, symmetry unique reflections.
[h]Average deviation between the observed and calculated structure factors calculated as: $R_{work} = \Sigma_{hkl} ||F_{obs}| - |F_{calc}|| / \Sigma_{hkl} |F_{obs}|$, where the $F_{obs}$ and $F_{calc}$ are the observed and calculated structure factor amplitudes of reflection hkl. $R_{free}$ is equal to $R_{factor}$ but for a randomly selected 5.0% subset of reflections that were held aside throughout refinement for cross-validation.
[i]Correlation coefficient between observed and calculated structure factor amplitudes.
[j]Real space correlation coefficient for the inhibitor and experimental density.
[k]According to MolProbity.

### Table S6. Diffraction data and refinement statistics — continued (2/2)

| **$M^{pro}$ variant** | WT | WT | WT | WT | S144A |
|---|---|---|---|---|---|
| **Inhibitor** | ML2006a | ML2006a2 | ML2006a4 | ML3006a | ML4006a |
| **PDB ID** | **8EZV** | **8EZZ** | **8F02** | **8F2C** | **8F2D** |
| **Data Collection** | | | | | |
| Beamline | SSRL BL12-2 | SSRL BL12-2 | SSRL BL12-2 | SSRL BL12-2 | SSRL BL12-2 |
| Wavelength (Å) | 1.03317 | 0.97946 | 0.97946 | 0.97946 | 0.97946 |
| Space group | $P2_12_12$ | $P2_12_12$ | $P2_12_12$ | $P2_12_12$ | $P2_12_12$ |
| Cell dimensions | | | | | |
| *a*, *b*, *c* (Å) | 45.82, 64.07, 104.42 | 45.49, 63.18, 103.65 | 45.65, 63.23, 103.80 | 45.58, 63.85, 104.38 | 45.40, 64.29 ,105.64 |
| *α*, *β*, *γ* (°) | 90, 90, 90 | 90, 90, 90 | 90, 90, 90 | 90, 90, 90 | 90, 90, 90 |
| Wilson B factor[a] | 26.1 | 30.1 | 44.3 | 34.3 | 22.7 |
| Matthews coef. ($Å^3$/Da)[b] | 2.27 | 2.20 | 2.21 | 2.24 | 2.28 |
| Solvent content (%) | 45.92 | 44.14 | 44.45 | 45.21 | 46.02 |
| Resolution (Å)[c] | 64.07 (1.80) | 40.07 (1.85) | 37.01 (2.00) | 37.09 (1.95) | 37.08 (1.95) |
| No. of reflections/unique | 210204 / 29174 | 113602 / 25876 | 87816 / 20441 | 79313 / 22645 | 89628 / 22625 |
| $R_{merge}$[d] | 0.145 (1.889) | 0.090 (1.334) | 0.108 (1.921) | 0.114 (1.614) | 0.150 (1.276) |
| $I/\sigma(I)$ [e] | 8.0 (1.3) | 7.2 (1.3) | 5.5 (0.7) | 7.1 (0.9) | 5.5 (1.5) |
| $CC_{1/2}$ | 0.995 (0.645) | 0.994 (0.444) | 0.997 (0.255) | 0.996 (0.312) | 0.961 (0.152) |
| Completeness (%)[f] | 99.6 (99.6) | 98.8 (98.6) | 97.9 (99.6) | 99.1 (99.3) | 97.6 (98.5) |
| Redundancy[g] | 7.2 (7.2) | 4.4 (4.5) | 4.3 (4.3) | 3.5 (3.5) | 4.0 (4.0) |
| **Refinement** | | | | | |
| Resolution (Å) | 54.67–1.80 | 40.10–1.85 | 37.04–2.00 | 34.98–1.95 | 37.11–1.95 |
| No. reflections/test set | 27728 / 1401 | 24499 / 1353 | 19391 / 1032 | 21542 / 1086 | 21545 / 1059 |
| $R_{work}$ / $R_{free}$[h] | 0.179 / 0.220 | 0.190 / 0.254 | 0.193 / 0.255 | 0.196 / 0.267 | 0.196 / 0.252 |
| $F_{obs}$-$F_{calc}$correlation[i] | 0.967 | 0.972 | 0.976 | 0.971 | 0.953 |
| No. atoms | | | | | |
| Protein | 2395 | 2393 | 2384 | 2382 | 2387 |
| Inhibitor | 42 | 44 | 44 | 41 | 42 |
| Inhibitor RSCC[j] | 0.95 | 0.96 | 0.95 | 0.95 | 0.94 |
| Other ligand/ion | - | 1 (Cl) | - | 1 (Cl) | 1 (Cl) |
| Water | 267 | 125 | 49 | 96 | 161 |
| Median *B*-factors | | | | | |
| Protein | 28 | 38 | 51 | 40 | 26 |
| Inhibitor | 23 | 35 | 48 | 42 | 28 |
| Ligand/Ion | - | 50 | - | 49 | 39 |
| R.m.s. deviations | | | | | |
| Bond lengths (Å) | 0.007 | 0.008 | 0.007 | 0.007 | 0.008 |
| Bond angles (°) | 1.479 | 1.569 | 1.520 | 1.515 | 1.500 |
| Ramachandran stats.[k] | | | | | |
| Favored+Allowed (%) | 100 | 100 | 100 | 100 | 100 |
| Outliers (%) | 0 | 0 | 0 | 0 | 0 |

[a]Overall B-factor value, an approximation to the fall-off of atomic scattering with resolution.
[b]Ratio of the volume of the asymmetric unit to the molecular weight of all protein molecules in the asymmetric unit.
[c]Value in parenthesis is for the highest-resolution shell: 1.80-1.84 Å in 8EZV, 1.85-1.89 Å in 8EZZ, 2.00-2.11 Å in 8F02, 1.95-2.06 Å in 8F2C, 1.95-2.00 Å in 8F2D. Values in parentheses in subsequent rows relates to this highest resolution shell.
[d]Reliability factor for symmetry-related reflections calculated as: $R_{merge} = \Sigma_{hkl}\Sigma j{=}1$ to N | $I_{hkl} - I_{hkl}$ (j) | / $\Sigma_{hkl}\Sigma j{=}1$ to N $I_{hkl}$ (j), where N is the redundancy of the data.
[e]Ratio of mean intensity to the mean standard deviation of the intensity over the entire resolution range.
[f]Fraction of measured reflections to possible observations at the resolution range.
[g]Number of measurements of individual, symmetry unique reflections.
[h]Average deviation between the observed and calculated structure factors calculated as: $R_{work} = \Sigma_{hkl} ||F_{obs}| - |F_{calc}|| / \Sigma_{hkl} |F_{obs}|$, where the $F_{obs}$ and $F_{calc}$ are the observed and calculated structure factor amplitudes of reflection hkl. $R_{free}$ is equal to $R_{factor}$ but for a randomly selected 5.0% subset of reflections that were held aside throughout refinement for cross-validation.
[i]Correlation coefficient between observed and calculated structure factor amplitudes.
[j]Real space correlation coefficient for the inhibitor and experimental density.
[k]According to MolProbity.

### Table S7 Hematology after 4-day repeated b.i.d. dosing

| Group ID | Exp ID | age (weeks) | Animal ID | WBC ($x10^9$ /L) | RBC ($x10^{12}$ /L) | HGB (g/L) | HCT (%) | MCV (fL) | MCH (pg) | MCHC (g/L) | PLT ($x10^9$ /L) | RDW-SD (fL) | RDW (%) | PDW (fL) | MPV (fL) |
|---|---|---|---|---|---|---|---|---|---|---|---|---|---|---|---|
| Group 1 Vehicle po b.i.d. | 1 | 8 - 10 | 2101* | - | - | - | - | - | - | - | - | - | - | - | - |
| | 1 | 8 - 10 | 2102 | 5.23 | 10.5 | 154 | 47.1 | 44.9 | 14.7 | 327 | 1272 | 20.9 | 17.5 | 6.80 | 6.50 |
| | 1 | 8 - 10 | 2103 | 3.13 | 10.4 | 158 | 48.3 | 46.4 | 15.2 | 327 | 1138 | 21.3 | 17.3 | 6.90 | 6.60 |
| | 2 | 10 - 12 | 2104 | 8.97 | 9.86 | 148 | 45.4 | 46.0 | 15.0 | 326 | 1087 | 20.5 | 16.2 | 6.70 | 6.60 |
| | 2 | 10 - 12 | 2105 | 7.79 | 9.74 | 146 | 45.2 | 46.4 | 15.0 | 323 | 1130 | 21.6 | 16.4 | 6.70 | 6.80 |
| | 2 | 10 - 12 | 2106 | 4.21 | 10.2 | 149 | 46.6 | 45.8 | 14.7 | 320 | 1155 | 20.8 | 16.8 | 6.80 | 6.70 |
| **Mean** | | | | **5.87** | **10.13** | **151.00** | **46.52** | **45.90** | **14.92** | **324.60** | **1156.40** | **21.02** | **16.84** | **6.78** | **6.64** |
| **SD** | | | | **2.45** | **0.32** | **4.90** | **1.28** | **0.62** | **0.22** | **3.05** | **69.31** | **0.43** | **0.56** | **0.08** | **0.11** |
| Group 2 RTV 20 mg/kg po b.i.d. | 1 | 8 - 10 | 2201 | 4.18 | 10.4 | 155 | 48.4 | 46.5 | 14.9 | 320 | 1017 | 21.0 | 17.0 | 6.70 | 6.80 |
| | 1 | 8 - 10 | 2202 | 6.02 | 11.0 | 162 | 49.6 | 45.1 | 14.7 | 327 | 1272 | 20.9 | 17.8 | 6.70 | 6.50 |
| | 1 | 8 - 10 | 2203 | 7.01 | 10.5 | 154 | 47.7 | 45.6 | 14.7 | 323 | 1004 | 20.4 | 17.2 | 6.80 | 6.80 |
| | 2 | 10 - 12 | 2204 | 2.68 | 9.99 | 145 | 45.5 | 45.5 | 14.5 | 319 | 1000 | 20.3 | 16.7 | 6.90 | 6.90 |
| | 2 | 10 - 12 | 2205 | 4.43 | 9.53 | 137 | 43.8 | 46.0 | 14.4 | 313 | 977 | 20.4 | 15.2 | 6.80 | 6.70 |
| | 2 | 10 - 12 | 2206 | 4.77 | 10.3 | 151 | 47.1 | 46.0 | 14.7 | 321 | 1060 | 20.9 | 16.8 | 6.60 | 6.60 |
| **Mean** | | | | **4.85** | **10.27** | **150.67** | **47.02** | **45.78** | **14.65** | **320.50** | **1055.00** | **20.65** | **16.78** | **6.75** | **6.72** |
| **SD** | | | | **1.51** | **0.49** | **8.69** | **2.08** | **0.49** | **0.18** | **4.64** | **109.79** | **0.31** | **0.87** | **0.10** | **0.15** |
| Group 3 ML2006a4/RTV 40/20 mg/kg po b.i.d. | 1 | 8 - 10 | 2301 | 6.46 | 10.5 | 156 | 47.9 | 45.7 | 14.9 | 326 | 1270 | 21.0 | 17.3 | 6.80 | 6.60 |
| | 1 | 8 - 10 | 2302 | 6.08 | 9.40 | 143 | 43.5 | 46.3 | 15.2 | 329 | 335 | 21.9 | 16.6 | 6.70 | 6.90 |
| | 1 | 8 - 10 | 2303 | 4.53 | 10.5 | 157 | 48.0 | 45.5 | 14.9 | 327 | 1166 | 19.6 | 17.0 | 6.80 | 6.50 |
| | 2 | 10 - 12 | 2304 | 2.68 | 9.95 | 146 | 45.0 | 45.2 | 14.7 | 324 | 1121 | 20.0 | 16.2 | 6.70 | 6.70 |
| | 2 | 10 - 12 | 2305 | 2.88 | 9.90 | 145 | 44.8 | 45.3 | 14.6 | 324 | 1126 | 20.4 | 16.5 | 6.70 | 6.80 |
| | 2 | 10 - 12 | 2306 | 4.12 | 8.97 | 137 | 42.1 | 46.9 | 15.3 | 325 | 872 | 23.5 | 16.6 | 6.80 | 7.00 |
| **Mean** | | | | **4.46** | **9.87** | **147.33** | **45.22** | **45.82** | **14.93** | **325.83** | **981.67** | **21.07** | **16.70** | **6.75** | **6.75** |
| **SD** | | | | **1.58** | **0.61** | **7.76** | **2.36** | **0.66** | **0.27** | **1.94** | **342.80** | **1.44** | **0.39** | **0.05** | **0.19** |
| Group 4 NTV/RTV 40/20 mg/kg po b.i.d. | 1 | 8 - 10 | 2401 | 4.20 | 10.9 | 164 | 50.4 | 46.1 | 15.0 | 325 | 172 | 21.6 | 17.9 | 7.80 | 6.80 |
| | 1 | 8 - 10 | 2402 | 4.54 | 10.3 | 155 | 46.8 | 45.7 | 15.1 | 331 | 1162 | 20.3 | 16.9 | 6.70 | 6.70 |
| | 1 | 8 - 10 | 2403 | 3.81 | 9.66 | 143 | 43.6 | 45.1 | 14.8 | 328 | 1110 | 20.7 | 16.7 | 6.80 | 6.70 |
| | 2 | 10 - 12 | 2404 | 5.59 | 9.48 | 142 | 44.1 | 46.5 | 15.0 | 322 | 1093 | 21.3 | 16.2 | 6.60 | 6.70 |
| | 2 | 10 - 12 | 2405 | 3.86 | 9.45 | 139 | 42.8 | 45.3 | 14.7 | 325 | 1065 | 20.8 | 16.5 | 6.80 | 6.80 |
| | 2 | 10 - 12 | 2406 | 2.69 | 8.04 | 122 | 38.9 | 48.4 | 15.2 | 314 | 919 | 22.6 | 13.5 | 7.20 | 7.20 |
| **Mean** | | | | **4.12** | **9.64** | **144.17** | **44.43** | **46.18** | **14.97** | **324.17** | **920.17** | **21.22** | **16.28** | **6.98** | **6.82** |
| **SD** | | | | **0.95** | **0.97** | **14.39** | **3.88** | **1.20** | **0.19** | **5.85** | **375.53** | **0.82** | **1.48** | **0.45** | **0.19** |

| Group ID | Animal ID | P-LCR (%) | PCT (%) | NRBC# (10^9/L) | NRBC% (%) | RET% (%) | RET# (10^12/L) | RET-He (pg) | IRF (%) | LFR (%) | MFR (%) | HFR (%) | WBC Differential (absolute) Neut ($x10^9$ /L) | Lymph ($x10^9$ /L) | Mono ($x10^9$/L) | Eos ($x10^9$/L) | Baso ($x10^9$/L) |
|---|---|---|---|---|---|---|---|---|---|---|---|---|---|---|---|---|---|
| Group 1 Vehicle po b.i.d. | 2101* | - | - | - | - | - | - | - | - | - | - | - | - | - | - | - | - |
| | 2102 | 3.10 | 1.12 | 0.00 | 0.00 | 4.70 | 0.493 | 17.3 | 64.5 | 35.5 | 14.7 | 49.8 | 0.560 | 4.48 | 0.150 | 0.0400 | 0.00 |
| | 2103 | 3.60 | 0.950 | 0.0100 | 0.300 | 4.50 | 0.468 | 17.5 | 64.2 | 35.8 | 13.8 | 50.4 | 0.230 | 2.80 | 0.0800 | 0.0200 | 0.00 |
| | 2104 | 3.10 | 0.970 | 0.0600 | 0.700 | 3.95 | 0.390 | 17.1 | 66.6 | 33.4 | 15.4 | 51.2 | 1.16 | 7.53 | 0.230 | 0.0400 | 0.0100 |
| | 2105 | 4.50 | 1.08 | 0.0400 | 0.500 | 4.23 | 0.412 | 17.2 | 64.8 | 35.2 | 14.7 | 50.1 | 1.13 | 6.44 | 0.180 | 0.0300 | 0.0100 |
| | 2106 | 4.10 | 1.06 | 0.0100 | 0.200 | 4.47 | 0.455 | 17.6 | 64.6 | 35.4 | 14.1 | 50.5 | 0.540 | 3.58 | 0.0800 | 0.0100 | 0.00 |
| **Mean** | | **3.68** | **1.04** | **0.02** | **0.34** | **4.37** | **0.44** | **17.34** | **64.94** | **35.06** | **14.54** | **50.40** | **0.72** | **4.97** | **0.14** | **0.03** | **0.00** |
| **SD** | | **0.62** | **0.07** | **0.03** | **0.27** | **0.29** | **0.04** | **0.21** | **0.95** | **0.95** | **0.62** | **0.52** | **0.41** | **1.97** | **0.07** | **0.01** | **0.01** |
| Group 2 RTV 20 mg/kg po b.i.d. | 2201 | 5.30 | 0.970 | 0.0100 | 0.200 | 4.24 | 0.441 | 17.1 | 63.6 | 36.4 | 14.2 | 49.4 | 0.390 | 3.57 | 0.120 | 0.0900 | 0.0100 |
| | 2202 | 3.30 | 0.910 | 0.0400 | 0.700 | 3.88 | 0.426 | 17.7 | 60.7 | 39.3 | 15.8 | 44.9 | 0.320 | 5.52 | 0.130 | 0.0500 | 0.00 |
| | 2203 | 4.30 | 0.750 | 0.00 | 0.00 | 4.35 | 0.455 | 17.4 | 63.4 | 36.6 | 14.3 | 49.1 | 0.590 | 6.01 | 0.150 | 0.260 | 0.00 |
| | 2204 | 4.40 | 0.900 | 0.0200 | 0.700 | 4.23 | 0.423 | 17.5 | 60.6 | 39.4 | 15.3 | 45.3 | 0.460 | 2.17 | 0.0400 | 0.0100 | 0.00 |
| | 2205 | 3.50 | 0.890 | 0.0100 | 0.200 | 3.53 | 0.336 | 17.7 | 59.0 | 41.0 | 14.6 | 44.4 | 0.870 | 3.48 | 0.0800 | 0.00 | 0.00 |
| | 2206 | 3.40 | 0.990 | 0.0200 | 0.400 | 3.47 | 0.356 | 17.3 | 59.7 | 40.3 | 13.3 | 46.4 | 0.520 | 4.12 | 0.110 | 0.0200 | 0.00 |
| **Mean** | | **4.03** | **0.90** | **0.02** | **0.37** | **3.95** | **0.41** | **17.45** | **61.17** | **38.83** | **14.58** | **46.58** | **0.53** | **4.15** | **0.11** | **0.07** | **0.00** |
| **SD** | | **0.78** | **0.08** | **0.01** | **0.29** | **0.38** | **0.05** | **0.23** | **1.91** | **1.91** | **0.88** | **2.17** | **0.19** | **1.42** | **0.04** | **0.10** | **0.00** |
| Group 3 ML2006a4/RTV 40/20 mg/kg po b.i.d. | 2301 | 4.00 | 1.00 | 0.0500 | 0.800 | 3.76 | 0.394 | 17.8 | 66.8 | 33.2 | 15.0 | 51.8 | 0.400 | 5.88 | 0.140 | 0.0400 | 0.00 |
| | 2302 | 4.90 | 0.540 | 0.0100 | 0.200 | 5.60 | 0.526 | 18.0 | 69.1 | 30.9 | 14.6 | 54.5 | 0.330 | 5.19 | 0.160 | 0.390 | 0.0100 |
| | 2303 | 3.00 | 0.910 | 0.0400 | 0.900 | 3.09 | 0.326 | 17.5 | 58.5 | 41.5 | 15.6 | 42.9 | 0.440 | 3.94 | 0.120 | 0.0300 | 0.00 |
| | 2304 | 3.70 | 1.08 | 0.0400 | 1.50 | 3.58 | 0.356 | 17.4 | 62.1 | 37.9 | 14.6 | 47.5 | 0.350 | 2.29 | 0.0400 | 0.00 | 0.00 |
| | 2305 | 4.10 | 1.01 | 0.0200 | 0.700 | 3.39 | 0.336 | 17.6 | 58.9 | 41.1 | 14.9 | 44.0 | 0.380 | 2.44 | 0.0600 | 0.00 | 0.00 |
| | 2306 | 4.90 | 0.800 | 0.0200 | 0.500 | 5.08 | 0.456 | 18.3 | 62.4 | 37.6 | 14.5 | 47.9 | 0.720 | 3.20 | 0.170 | 0.0300 | 0.00 |
| **Mean** | | **4.10** | **0.89** | **0.03** | **0.77** | **4.08** | **0.40** | **17.77** | **62.97** | **37.03** | **14.87** | **48.10** | **0.44** | **3.82** | **0.12** | **0.08** | **0.00** |
| **SD** | | **0.73** | **0.20** | **0.02** | **0.44** | **1.01** | **0.08** | **0.34** | **4.24** | **4.24** | **0.41** | **4.45** | **0.14** | **1.47** | **0.05** | **0.15** | **0.00** |
| Group 4 NTV/RTV 40/20 mg/kg po b.i.d. | 2401 | 2.10 | 0.180 | 0.0100 | 0.200 | 5.11 | 0.559 | 17.4 | 66.5 | 33.5 | 15.2 | 51.3 | 0.300 | 3.77 | 0.0600 | 0.0600 | 0.0100 |
| | 2402 | 4.10 | 1.06 | 0.0100 | 0.200 | 3.43 | 0.352 | 17.3 | 60.6 | 39.4 | 15.4 | 45.2 | 0.760 | 3.68 | 0.0800 | 0.0100 | 0.0100 |
| | 2403 | 4.10 | 1.01 | 0.0100 | 0.300 | 4.08 | 0.394 | 18.0 | 62.2 | 37.8 | 14.7 | 47.5 | 0.470 | 3.23 | 0.100 | 0.0100 | 0.00 |
| | 2404 | 3.70 | 0.960 | 0.0400 | 0.700 | 4.43 | 0.420 | 18.4 | 67.9 | 32.1 | 14.2 | 53.7 | 0.650 | 4.77 | 0.160 | 0.0100 | 0.00 |
| | 2405 | 3.90 | 0.970 | 0.0200 | 0.500 | 3.97 | 0.375 | 17.6 | 64.5 | 35.5 | 14.8 | 49.7 | 0.610 | 3.13 | 0.110 | 0.0100 | 0.00 |
| | 2406 | 5.50 | 0.970 | 0.0400 | 1.50 | 4.44 | 0.357 | 18.2 | 66.9 | 33.1 | 14.7 | 52.2 | 0.350 | 2.23 | 0.0900 | 0.0100 | 0.0100 |
| **Mean** | | **3.90** | **0.86** | **0.02** | **0.57** | **4.24** | **0.41** | **17.82** | **64.77** | **35.23** | **14.83** | **49.93** | **0.52** | **3.47** | **0.10** | **0.02** | **0.01** |
| **SD** | | **1.09** | **0.33** | **0.01** | **0.50** | **0.56** | **0.08** | **0.45** | **2.88** | **2.88** | **0.42** | **3.15** | **0.18** | **0.84** | **0.03** | **0.02** | **0.01** |

### Table S8 Clinical biochemistry after 4-day repeated b.i.d. dosing

| | Exp ID | age (weeks) | Animal ID | ALB (g/L) | ALP (U/L) | ALT (U/L) | AST (U/L) | Ca (mmol/L) | TC (mmol/L) | CK (U/L) | D-Bil (μmol/L) | γ-GT (U/L) | GLU (mmol/L) | P (mmol/L) | LDH (U/L) | T-Bil (μmol/L) | Mg (mmol/L) | TP (g/L) | TG (mmol/L) | UREA (mmol/L) | UA (μmol/L) | CREA (μmol/L) | I-Bil (μmol/L) | GLB (g/L) | A/G (ALB/GLB) |
|---|---|---|---|---|---|---|---|---|---|---|---|---|---|---|---|---|---|---|---|---|---|---|---|---|---|
| **Group 1** Vehicle po b.i.d. | 1 | 8 - 10 | 2101 | 38.2 | 224 | 36.0 | 104 | 2.40 | 2.25 | 1222 | BQL | BQL | 5.00 | 2.70 | 448 | BQL | 1.41 | 57.4 | 1.25 | 11.0 | BQL | 15.5 | NA | 19.2 | 1.99 |
| | 1 | 8 - 10 | 2102 | 39.2 | 196 | 33.0 | 83.0 | 2.40 | 2.30 | 128 | BQL | BQL | 5.70 | 2.00 | 317 | BQL | 1.41 | 59.8 | 1.15 | 12.7 | BQL | 14.0 | NA | 20.6 | 1.91 |
| | 1 | 8 - 10 | 2103 | 40.2 | 217 | 29.5 | 86.0 | 2.50 | 2.30 | 206 | BQL | BQL | 7.40 | 2.00 | 335 | BQL | 1.45 | 61.0 | 0.950 | 12.4 | BQL | 14.5 | NA | 20.8 | 1.93 |
| | 2 | 10 - 12 | 2104 | 33.66 | 141.9 | 45.0 | 161.4 | 2.43 | 1.95 | 521.85 | BQL | BQL | 6.12 | 3.33 | 426.93 | 2.58 | 1.482 | 57.18 | 1.17 | 12.66 | BQL | 15.9 | NA | 23.52 | 1.43 |
| | 2 | 10 - 12 | 2105 | 30.78 | 148.2 | 88.8 | 171.9 | 2.34 | 1.41 | 1318.68 | BQL | BQL | 9.69 | 2.88 | 538.59 | 2.25 | 1.35 | 51.81 | 0.87 | 14.61 | BQL | 18.3 | NA | 21.03 | 1.46 |
| | 2 | 10 - 12 | 2106 | 34.74 | 171.3 | 39.3 | 181.8 | 2.37 | 1.86 | 1118.61 | 1.92 | BQL | 10.98 | 2.43 | 433.47 | 3.66 | 1.272 | 57.42 | 0.93 | 15.66 | BQL | 16.5 | 1.74 | 22.68 | 1.53 |
| **Mean** | | | | **36.13** | **182.90** | **45.27** | **131.27** | **2.41** | **2.01** | **752.46** | **1.92** | **-** | **7.48** | **2.56** | **416.47** | **2.83** | **1.39** | **57.43** | **1.05** | **13.16** | **-** | **15.78** | **1.74** | **21.30** | **1.71** |
| **SD** | | | | **3.66** | **34.63** | **21.98** | **45.30** | **0.06** | **0.35** | **532.48** | **-** | **-** | **2.38** | **0.52** | **81.10** | **0.74** | **0.07** | **3.16** | **0.16** | **1.68** | **-** | **1.53** | **-** | **1.56** | **0.26** |
| *Mean - 3SD* | | | | *25.15* | *79.01* | *-20.68* | *-4.65* | *2.24* | *0.96* | *-844.98* | - | - | *0.34* | *0.99* | *173.18* | *0.62* | *1.17* | *47.96* | *0.59* | *8.12* | - | *11.18* | - | *16.62* | *0.93* |
| *Mean + 3SD* | | | | *47.11* | *286.79* | *111.21* | *267.18* | *2.57* | *3.06* | *2349.89* | - | - | *14.62* | *4.12* | *659.75* | *5.04* | *1.62* | *66.89* | *1.52* | *18.21* | - | *20.39* | | *25.97* | *2.49* |
| **Group 2** RTV 20 mg/kg po b.i.d. | 1 | 8 - 10 | 2201 | 35.3 | 152 | 41.0 | 92.0 | 2.30 | 2.40 | 112 | BQL | BQL | 3.85 | 2.20 | 368 | BQL | 1.29 | 54.6 | 1.45 | 8.75 | BQL | 14.5 | NA | 19.3 | 1.83 |
| | 1 | 8 - 10 | 2202 | 39.5 | 179 | 41.5 | 64.0 | 2.45 | 2.40 | 121 | BQL | BQL | 5.45 | 2.25 | 266 | BQL | 1.31 | 58.1 | 1.30 | 10.8 | BQL | 19.0 | NA | 18.6 | 2.12 |
| | 1 | 8 - 10 | 2203 | 38.2 | 189 | 43.0 | 325 | 2.55 | 2.65 | 1426 | BQL | BQL | 4.90 | 3.00 | 727 | BQL | 1.39 | 58.0 | 1.20 | 10.8 | BQL | 17.5 | NA | 19.8 | 1.93 |
| | 2 | 10 - 12 | 2204 | 32.40 | 146.7 | 50.4 | 262.2 | 2.37 | 1.68 | 1068.15 | BQL | BQL | 6.18 | 3.09 | 706.98 | BQL | 1.311 | 54.45 | 0.96 | 10.98 | BQL | 18.0 | NA | 22.05 | 1.47 |
| | 2 | 10 - 12 | 2205 | 32.40 | 163.8 | 40.8 | 112.8 | 2.46 | 1.50 | 444.24 | 1.68 | BQL | 6.96 | 3.12 | 395.43 | 3.48 | 1.221 | 53.70 | 1.32 | 14.85 | BQL | 19.5 | 1.80 | 21.30 | 1.52 |
| | 2 | 10 - 12 | 2206 | 36.36 | 168.0 | 46.5 | 281.4 | 2.61 | 2.16 | 1286.58 | BQL | BQL | 7.08 | 3.84 | 675.51 | 3.12 | 1.482 | 62.19 | 1.29 | 12.54 | BQL | 20.4 | NA | 25.83 | 1.41 |
| **Mean** | | | | **35.69** | **166.25** | **43.87** | **189.57** | **2.46** | **2.13** | **743.07** | **1.68** | **-** | **5.74** | **2.92** | **523.03** | **3.30** | **1.33** | **56.83** | **1.25** | **11.45** | **-** | **18.15** | **1.80** | **21.15** | **1.71** |
| **SD** | | | | **2.94** | **15.98** | **3.84** | **112.45** | **0.11** | **0.45** | **590.27** | **-** | **-** | **1.25** | **0.61** | **202.52** | **0.25** | **0.09** | **3.24** | **0.16** | **2.05** | **-** | **2.07** | **-** | **2.63** | **0.29** |
| **Group 3** ML2006a4/RTV 40/20 mg/kg po b.i.d. | 1 | 8 - 10 | 2301 | 36.2 | 184 | 32.0 | 83.0 | 2.50 | 3.00 | 174 | BQL | BQL | 3.25 | 2.20 | 252 | BQL | 1.40 | 56.5 | 1.50 | 11.6 | BQL | 14.5 | NA | 20.4 | 1.78 |
| | 1 | 8 - 10 | 2302 | 40.8 | 197 | 45.0 | 105 | 2.55 | 2.90 | 464 | BQL | BQL | 3.85 | 2.65 | 582 | BQL | 1.42 | 60.9 | 1.75 | 10.5 | BQL | 15.0 | NA | 20.1 | 2.03 |
| | 1 | 8 - 10 | 2303 | 37.5 | 138 | 39.0 | 81.5 | 2.30 | 2.85 | 105 | BQL | BQL | 4.75 | 1.85 | 445 | BQL | 1.24 | 57.4 | 1.25 | 11.1 | BQL | 17.0 | NA | 20.0 | 1.88 |
| | 2 | 10 - 12 | 2304 | 30.57 | 153.6 | 97.8 | 247.2 | 2.28 | 1.68 | 420.87 | 1.92 | BQL | 5.76 | 2.43 | 545.88 | 3.66 | 1.347 | 52.62 | 1.05 | 8.70 | BQL | 16.2 | 1.74 | 22.05 | 1.39 |
| | 2 | 10 - 12 | 2305 | 34.29 | 133.5 | 48.0 | 112.8 | 2.40 | 2.04 | 261.30 | BQL | BQL | 10.41 | 3.00 | 431.94 | 2.58 | 1.254 | 58.38 | 1.11 | 14.79 | BQL | 18.6 | NA | 24.09 | 1.42 |
| | 2 | 10 - 12 | 2306 | 31.98 | 141.3 | 45.0 | 128.7 | 2.43 | 1.80 | 457.95 | BQL | BQL | 8.61 | 3.48 | 389.97 | 2.76 | 1.32 | 52.62 | 0.99 | 15.36 | BQL | 17.1 | NA | 20.64 | 1.55 |
| **Mean** | | | | **35.20** | **157.73** | **51.13** | **126.37** | **2.41** | **2.38** | **313.80** | **1.92** | **-** | **6.11** | **2.60** | **441.06** | **3.00** | **1.33** | **56.40** | **1.28** | **11.99** | **-** | **16.40** | **1.74** | **21.20** | **1.68** |
| **SD** | | | | **3.73** | **26.37** | **23.56** | **61.87** | **0.11** | **0.60** | **155.36** | **-** | **-** | **2.83** | **0.58** | **117.80** | **0.58** | **0.07** | **3.27** | **0.30** | **2.58** | **-** | **1.50** | **-** | **1.61** | **0.26** |
| **Group 4** NTV/RTV 40/20 mg/kg po b.i.d. | 1 | 8 - 10 | 2401 | 41.2 | 176 | 127 | 532 | 2.45 | 3.70 | 2410 | BQL | BQL | 5.00 | 3.75 | 1855 | BQL | 1.70 | 64.8 | 1.90 | 11.1 | BQL | 23.5 | NA | 23.6 | 1.75 |
| | 1 | 8 - 10 | 2402 | 39.4 | 211 | 40.5 | 112 | 2.50 | 2.55 | 446 | BQL | BQL | 3.45 | 2.20 | 395 | BQL | 1.32 | 58.1 | 1.75 | 11.9 | BQL | 19.0 | NA | 18.8 | 2.10 |
| | 1 | 8 - 10 | 2403 | 37.0 | 184 | 48.0 | 76.5 | 2.45 | 2.75 | 122 | BQL | BQL | 5.00 | 2.30 | 390 | BQL | 1.27 | 57.1 | 1.25 | 12.4 | BQL | 15.0 | NA | 20.1 | 1.84 |
| | 2 | 10 - 12 | 2404 | 31.71 | 181.8 | 56.1 | 513.3 | 2.25 | 2.40 | 2560.89 | 3.48 | BQL | 9.75 | 3.00 | 1094.88 | 8.49 | 1.272 | 55.35 | 1.53 | 14.94 | BQL | 23.4 | 5.01 | 23.64 | 1.34 |
| | 2 | 10 - 12 | 2405 | 35.70 | 177.6 | 34.5 | 90.6 | 2.31 | 2.76 | 406.83 | BQL | BQL | 4.47 | 2.25 | 276.48 | 2.43 | 1.149 | 60.54 | 1.05 | 9.75 | BQL | 18.9 | NA | 24.84 | 1.44 |
| | 2 | 10 - 12 | 2406 | 33.36 | 167.6 | 434.0 | 996.4 | 2.04 | 2.56 | 2872.68 | BQL | BQL | 6.12 | 2.92 | 1688.24 | 1.02 | 1.268 | 58.44 | 1.56 | 11.64 | BQL | 21.6 | NA | 25.08 | 1.33 |
| **Mean** | | | | **36.38** | **182.83** | **123.27** | **386.72** | **2.33** | **2.79** | **1469.68** | **3.48** | **-** | **5.63** | **2.74** | **949.94** | **3.98** | **1.33** | **59.04** | **1.51** | **11.95** | **-** | **20.23** | **5.01** | **22.66** | **1.63** |
| **SD** | | | | **3.57** | **14.89** | **155.87** | **365.65** | **0.17** | **0.47** | **1267.92** | **-** | **-** | **2.20** | **0.61** | **701.29** | **3.97** | **0.19** | **3.28** | **0.31** | **1.72** | **-** | **3.26** | **-** | **2.62** | **0.31** |

Values in red are outside the interval defined by mean±3SD of group 1
